# Complete biosynthesis of penicillin G in *Nicotiana benthamiana*

**DOI:** 10.64898/2026.05.08.723852

**Authors:** Abdul Rawoof, Yen Tung Lin, Sanjeevan Rajendran, Gaelle Antoine, Sameera Jayasundara, Yingqi Cai, Digar Singh, Payton Whitehead, Haley Dornberger, Sonya Mall, Ana P. Alonso, Michael C. Carroll, Elizabeth Skellam, Kent D. Chapman

## Abstract

Commercial penicillin production has relied on microbial fermentation for more than 80 years. Here, we engineered the plant, *Nicotiana benthamiana*, to produce penicillin G in its leaves by transient expression of up to seven fungal biosynthetic genes. Remarkably, all recombinant proteins localize to the analogous subcellular compartments without engineering signal peptide sequences or post-translational modification sites. Although non-ribosomal peptide synthetases occur widely in fungi and bacteria to produce a plethora of specialized metabolites, their evolutionary distribution does not extend to plants. Our results now open a new metabolic frontier for natural product synthesis, and offer possibilities to address global health concerns through an alternative biotechnology platform for fungal-derived pharmaceutical production.

## Main Text

Fungi are an important source of essential medicines including the β-lactam antibiotic penicillin G (also known as benzylpenicillin) (*1*). Since the discovery of penicillin in 1928, and the subsequent combined war-time efforts of both the UK and US to manufacture at scale (*2*), deep-vat fermentation has been the major approach for commercial production of natural and semi-synthetic β-lactam antibiotics. Fungi synthesize penicillin G from L−α-<u>a</u>minoadipic acid, <u>c</u>ysteine and <u>v</u>aline *via* the large multifunctional non-ribosomal peptide synthetase (NRPS), known as ACV synthase (ACVS) which is localized in small vacuoles and the cytosol (*3,4*). The ACV tripeptide product is converted to isopenicillin N by the 2-oxoglutarate-dependent dioxygenase isopenicillin N synthase (IPNS), localized in the cytoplasm (*5*) that catalyzes the formation of the β-lactam pharmacophore essential for bioactivity. Subsequently, the multifunctional isopenicillin *N*-acyltransferase (IAT), localized to the peroxisomes (*6*), removes the L−α-aminoadipyl side chain and replaces it with a phenylacetyl side chain (Figure 1). ACVS requires a post-translational modification by a phosphopantetheinyl transferase (PPTase) to be active in its *holo* form (*7*) and phenylacetyl-CoA ligase (PCL), localized to peroxisomes, is required for converting phenylacetic acid into phenylacetyl-CoA (*8,9*). Three transporters have been identified for shuttling intermediates between compartments (*10*): PenV, transports L-α-aminoadipic acid from the fungal vacuole to the cytoplasm and is located at the vacuolar membrane (*11*); PenM transports isopenicillin N from the cytoplasm to peroxisomes (*12*); and PaaT transports phenylacetic acid from the cytoplasm to peroxisomes, preventing self-toxicity (*13*). Finally, the transport of penicillin G from peroxisomes, to the outside of the cell proceeds *via* an unknown mechanism, but industrially engineered strains of fungi are reported to produce ∼ 50 g/L of penicillins by modern deep vat fermentation approaches (*14*), orders of magnitudes higher than the reported yields in original isolates (*15*).

**Fig. 1.**
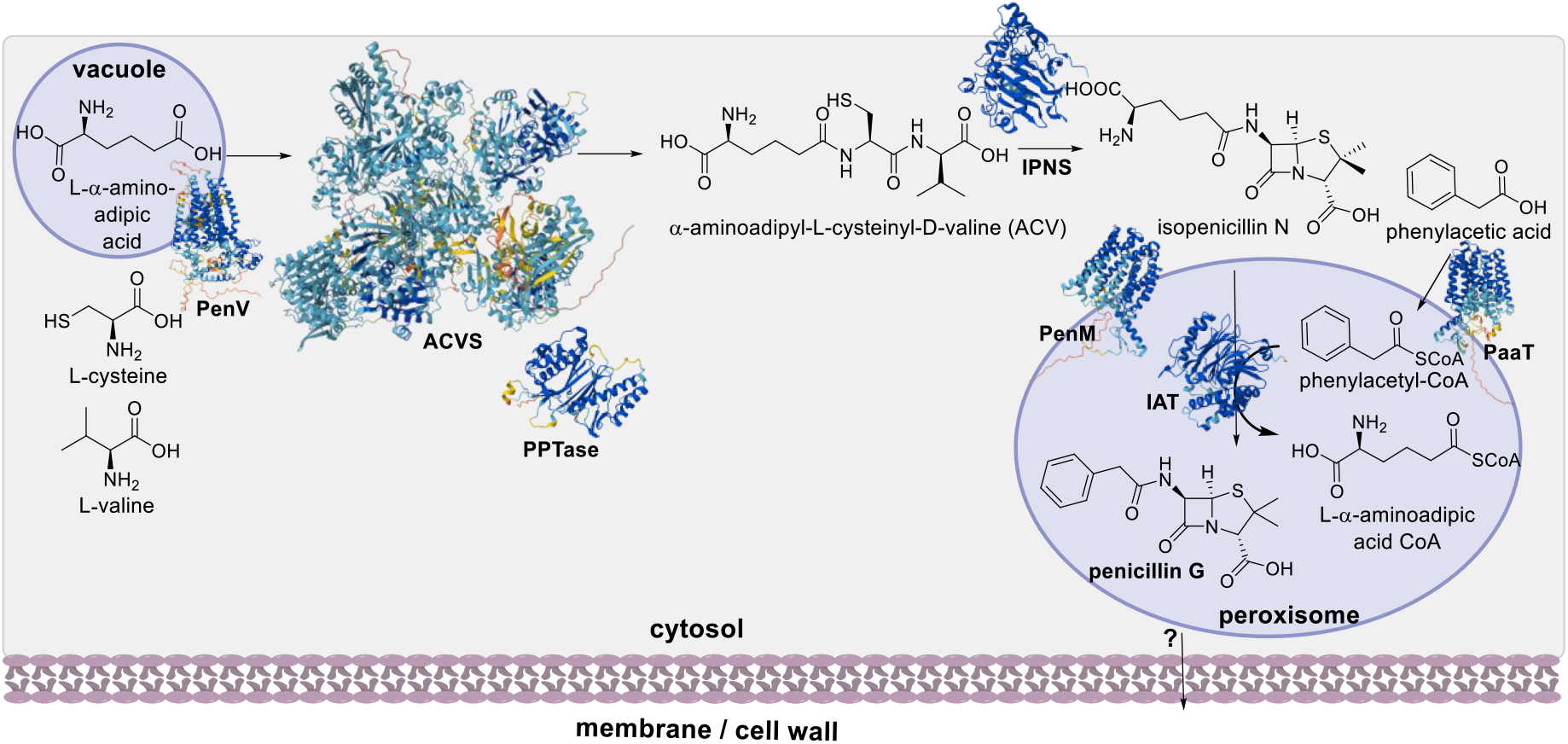
Biosynthesis of penicillin G in fungi. ACVS consists of three modules each containing an adenylation (A) domain and a thiolation (T) domain; two modules contain a condensation (C) domain. The A domains select and activate a specific amino acid *i*.*e*. L-α-amino-adipic acid, L-cysteine, and L-valine, which are transferred to the T domain, after post-translational modification with a phosphopantetheinyltransferase (PPTase). The C domain catalyzes condensation reactions between the amino acid precursors resulting in an elongated peptide chain. The third and final module contains an epimerization (E) domain, which interconverts the stereochemistry of L-valine to D-valine, in addition to a thioesterase (TE) domain which releases the α-aminoadipyl-L-cysteinyl-D-valine tripeptide product (ACV) *via* hydrolysis. Protein sequences of biosynthetic enzymes and transporters are listed in Table S4 and AlphaFold 3 (*16*) was used to generate structural models.

An emerging alternative to fermentation is molecular farming (or biopharming) where recombinant proteins, vaccines, and plant-made pharmaceuticals (PMPs) are produced in plants (*17*). Tobacco-related plants *i*.*e. Nicotiana* sp. are frequently used as a host platform due to high biomass yield, large leaf size, advanced PMP production methods, and large-scale production *via* greenhouses or even open fields (*18*). The benefits of plant-based expression systems over deep-vat fermentation include convenience, metabolic plasticity, lower capital investment into required infrastructure, and potential ease of scalability. Combined with transient expression approaches, such as *Agrobacterium*-mediated transformation, required genes can be rapidly introduced and propagated into the host plant, leading to PMP production in a few days (*19*). The lower cost associated with plant-based systems could be advantageous to lower-income countries which experience economic barriers for production of their medicines (*20*). While there are a wide range of recombinant proteins produced in plant systems, utilizing plants to synthesize bioactive molecules of microbial origin *via* recombinant, megasynthetase proteins has not been widely reported. A possible reason for this is that many of the essential medicines produced by microbes arise from large (>200 – 1000 kDa) multi-functional enzymes such as non-ribosomal peptide synthetases (NRPS) that are absent in plants (*21-23*).

Here, we reconstituted the complete production of penicillin G in the plant host *N. benthamiana via* the cytoplasmic NRPS megasynthetase, ACVS, and associated requisite pathway enzymes in their respective subcellular compartments. All pathway intermediates were identified using high-resolution mass spectrometry. Expansion of this proof-of-concept approach to other microbially-derived medicines and additional optimization may enable low- and middle-income countries to rapidly produce essential medicines and contribute to versatile management of emerging and future global health crises.

### Confocal fluorescence microscopy confirms analogous sub-cellular localization in plants and fungi

*Penicillium chrysogenum* is the well-studied industrial producer of penicillin, where the biosynthetic pathway consists of three genes *pcbAB, pcbC, penDE*, encoding ACVS, INPS, and IAT respectively (24). The genes *penV, penM*, and *paaT*, encoding the respective transporters, are located outside of this biosynthetic gene cluster (BGC) (*10-13*), as is *phl* encoding PCL (*8,9*), and *pc13g04050*, the native PPTase. NpgA is a universally utilized PPTase from *Aspergillus nidulans* used for post-translationally modifying megasynth(et)ases in non-filamentous fungal hosts (*25*). With the exception of *pcbAB, pc13g04050*, and *phl*, the codon-optimized genes, *npgA, pcbC, penDE, paaT*, and *penM*, were synthesized and individually cloned under the control of the UBQ10p or 35S promoter with a C- or N-terminal GFP in-frame fusion. The resulting chimeric gene fusions were co-expressed with either the cytoplasmic marker protein mCherry or the peroxisomal marker mCherry-SKL (containing the C-terminal Ser-Lys-Leu peroxisomal targeting signal) *via Agrobacterium*-mediated infiltration and transient transformation in *N. benthamiana* leaves.

Overlapping fluorescence distribution patterns confirmed the expected localization in the cytoplasm for Pc13g04050 (Fig. 2A), NpgA (Fig. 2B), ACVS (Fig. 2C,D), and IPNS (Fig. 2E). IAT was targeted to the plant peroxisomes as evidenced by its co-localization with the peroxisomal marker, mCherry-SKL (Fig. 2F). All localizations were quantified from multiple images by Fuji-based morphometric analysis and algorithmic scores supported the conclusion that these enzymes were in their correct subcellular compartments (Supplemental Figure S1). The ACVS protein appeared to be stabilized by co-expression with either fungal PPTase, since fluorescence signals from ACVS-GFP were barely detectable when expressed alone (Supplemental Fig. S2A); prolonged expression appeared to result in aggregated fluorescent ACVS structures even in the presence of PPTase (Supplemental Fig. S2B). The localization of PaaT appears to not be located exclusively to (or adjacent to) peroxisomes as expected but may also be associated with the ER-membrane based on the membrane-like fluorescent pattern and the possession of EAPR and KAIK ER-membrane retention signals (Supplemental Fig. S3). The sub-cellular localization of PenM was unclear as GFP-fusions to either end of the protein did not reveal the presence of a fluorescent signal, despite expression of coding RNAs (Supplemental Fig. S3). PenV was not included in the investigation since in plants L−α-aminoadipic acid is available in the cytoplasm *via* lysine catabolism (*26*).

**Fig. 2.**
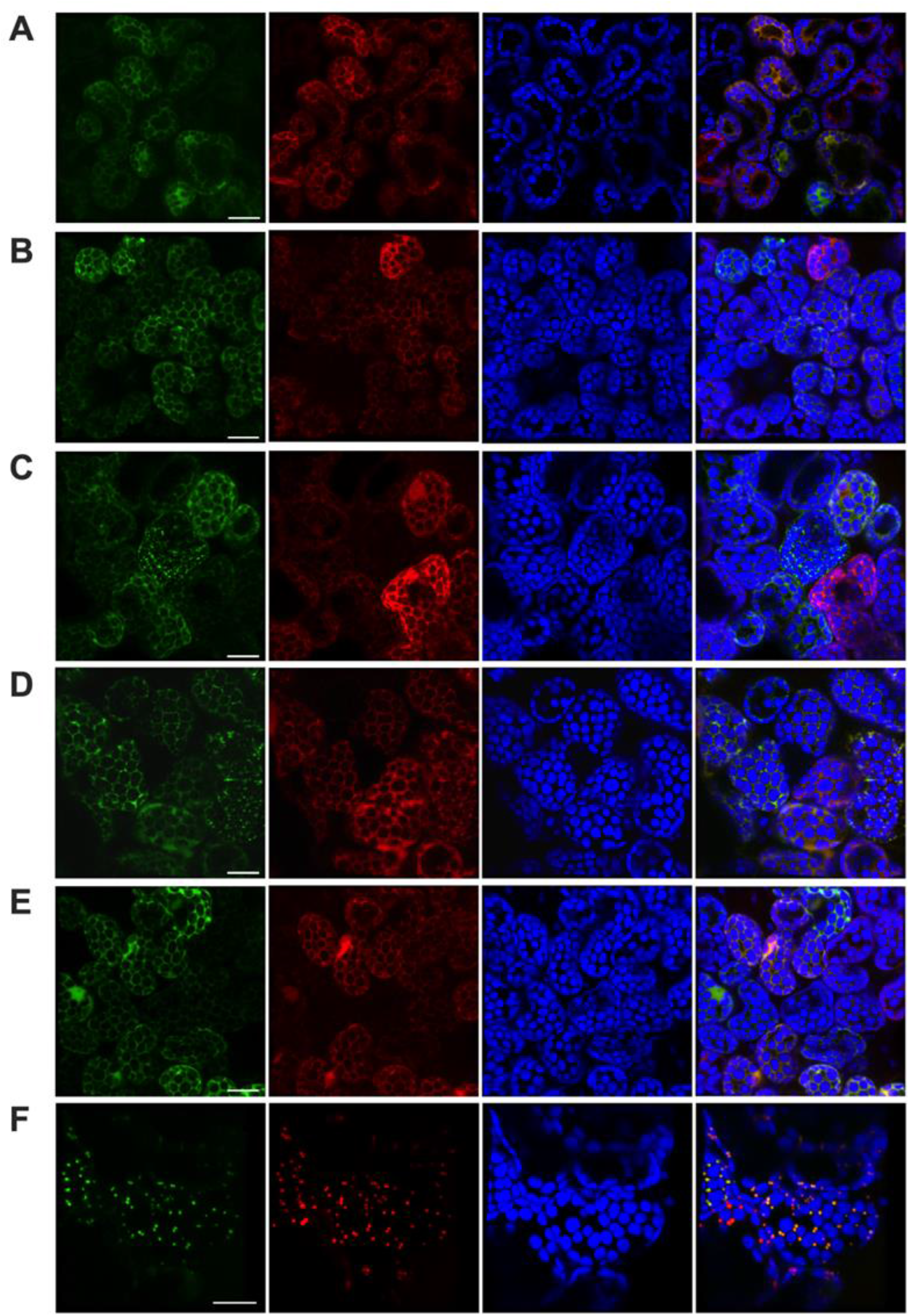
Subcellular localization of fungal biosynthetic enzymes in *N. benthamiana*. High resolution confocal images for subcellular localization of: A) NpgA-GFP localization (green), mCherry localization (red), chloroplasts (false-colored blue), and merged images (of NpgA-GFP, mCherry and chloroplasts); B) Pc13g04050-GFP localization, mCherry, chloroplasts, and merged images; C) ACVS-GFP (co-expressed with *npgA*, no tag), mCherry, chloroplasts, and merged images; D) ACVS-GFP (co-expressed with *Pc13g04050*, no tag), mCherry, and merged images; E) IPNS-GFP, mCherry, chloroplasts, and merged; F) GFP-IAT, mCherry-SKL marking the peroxisomes), chloroplasts, and merged images. Abbreviations: SKL = peroxisome targeting signal. Bar = 20 μm.

### ACV, isopenicillin N, and penicillin G production in plant tissues

With the correct localization of individual enzymes established, we performed a series of co-expression experiments to reconstruct the biosynthesis of ACV, isopenicillin N (IPN) and penicillin G (PenG) in *N. benthamiana* leaves (all utilizing the P19 viral protein to suppress transgene silencing (*27*)). Co-expression of ACVS with either NpgA or Pc13g04050 led to production of a metabolite with *m/z* 362.138±0.005 [M-H]^-^ in extracted ion chromatograms (XIC) corresponding to chemical formula C_14_H_24_N_3_O_6_S that matched the retention time and MS/MS fragmentation pattern of a commercial standard of ACV (Fig. S4A&B), although the co-expression of Pc13g04050 appeared considerably more efficient at producing activated, functional ACVS. No ACV was detected in mock-infected or P19-only controls. A small amount of ACV also could be detected in the absence of fungal PPTase, suggesting that the endogenous *N. benthamiana* enzyme(s) could activate the ACVS protein, albeit with less efficiency than the “native” fungal PPTase enzymes (Fig S4C). Mirror plots of MS fragmentations of the ACV commercial standard compared with the ACV metabolite extracted from the *N. benthamiana* leaves, confirmed the identity of the presumed metabolite (Fig S4D).

Co-expression of ACVS with either NpgA or Pc13g04050 and both IPNS and IAT led to production of a metabolite with *m/z* at 333.091 ± 0.005 [M–H]^−^ in XICs corresponding to a chemical formula of C_16_H_17_N_2_O_4_S that matched the retention time and MS/MS fragmentation pattern of a commercial standard of penicillin G (Fig. 3). No PenG was detected in mock-infected or P19-only controls. A very small amount of PenG was detected in the absence of fungal PPTase, supporting the observation that native *N. benthamiana* enzymes can activate ACVS (Fig. 3C). Highest amounts of PenG were recovered when the fungal *phl* gene was co-expressed with all pathway enzymes. However, endogenous CoA-ligases in *N. benthamiana* appear also to function adequately for PenG production to proceed in the absence of fungal *phl*. Mirror plots of the MS fragmentations of the PenG commercial standard compared with the PenG metabolite extracted from the *N. benthamiana* leaves confirmed the identity of the presumed metabolite (Fig 3D).

**Fig. 3:**
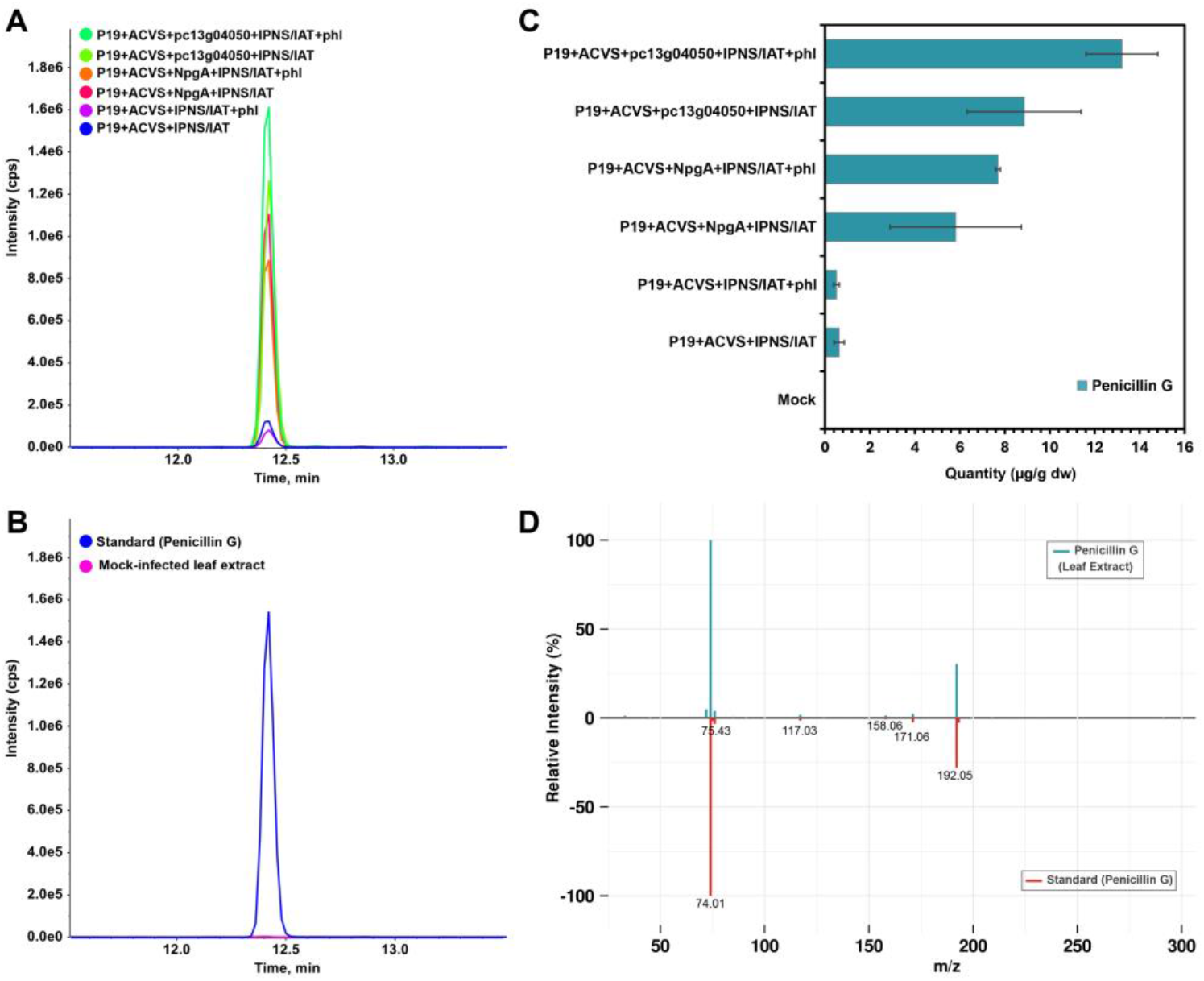
Production and quantification of PenG in *N. benthamiana* expressing penicillin G biosynthetic genes, analyzed by reverse-phase LC-MS/MS (TripleTOF 6600+, negative ESI). (A) Extracted ion chromatograms (XICs) of penicillin G, detected as [M–H]^−^ at *m/z* 333.091 ± 0.005 and eluting at 12.4 min, from leaves transiently expressing different combinations of pathway genes; (B) XIC comparison of an external penicillin G standard and a mock-infiltrated leaf extract (negative control), confirming the retention time and mass accuracy of detected penicillin G signal in planta, (C) Absolute penicillin G accumulation (µg g^−1^ dry weight) across the expression conditions shown in panel A, based on an external calibration curve (mean ± SD; *n* = 3), and (D) Mirror plot of MS/MS fragmentation spectra for penicillin G from leaf extracts compared with external penicillin G standard. Concordant fragment ions confirms the structural identity of penicillin G produced in *N. benthamiana*.

In addition to ACV and PenG production, a metabolite with *m/z* 358.107 [M-H]^-^ in XICs corresponding to a chemical formula of C_14_H_20_N_3_O_6_S, expected for IPN was detected in leaf extracts expressing fungal pathway enzymes and its formation was dependent upon expression of the fungal IPNS (Supplemental Fig. S5). Commercial standards for IPN were not available; however, the MS/MS fragmentation pattern of this molecule was consistent with *in silico* fragmentation of isopenicillin N obtained from the MassBank of North America (MoNA) (Supplemental Fig. S5D).

Reconstitution of PenG production (Fig.3), also resulted in accumulation of pathway intermediates ACV (Fig. S6) and IPN (Fig. S7), indicating that the complete pathway was fully functional. Conditions favoring the most PenG production, appeared also to favor accumulation of the highest amounts of intermediates (compare Fig. 3C with Fig. S6C and Fig. S7C). In mock infiltrations with buffer alone, or when Agrobacterium harboring P19 was infiltrated alone, none of the pathway metabolites—ACV, IPN, or PenG--could be detected using targeted or untargeted high-resolution electrospray ionization mass spectrometry (HRESIMS) confirming that they are indeed products of the infiltrated fungal enzymes (Fig. S4B, Fig. 3B, Fig. S5B, S6B and S7B). A representative total ion chromatogram of leaf extract (infiltrated with the entire pathway DNA coding sequences) along with selected XICs for ACV, IPN and PenG are shown in Supplemental Fig. S8 for context and complete comparisons. Quantification of ACV (Fig S4C, S6C) and PenG (Fig. 3C) were based on standard curves generated by chromatography and ESI of commercially available standards (Supplemental Figs. S9, S10). Additional care was exercised in compound identification, since fragmentation and MS/MS conditions can vary depending on matrix composition, analyte concentration, and fragmentation conditions (e.g., Supplemental Fig S11 for ACV).

We further tested the influence of fungal peroxisomal transporter PaaT on PenG accumulation (Fig. 4). Pen G was produced in the absence of PaaT, indicating that this transporter (or PenM) was not strictly required to complete the correct formation of PenG in peroxisomes. In other words, endogenous peroxisomal transporters could shuttle fungal intermediates into the peroxisomal matrix for IAT action. This is not particularly surprising given the promiscuity of plant peroxisomal transporters for various metabolic and hormone substrates (28). On the other hand, co-expression of PaaT enhanced the synthesis and accumulation of PenG by three-fold compared to without PaaT. PaaT transporter proteins having GFP fusions at either the N- or C-terminus (43-PaaT or 84-PaaT, respectively) were equally effective at enhancing the accumulation of PenG. By contrast, the co-expression of PenM fusion constructs with PenG pathway enzymes had no impact on overall PenG accumulation, and this was likely due to lack of detectable PenM fluorescent protein *in planta* (Fig. S3). Measurements of pathway intermediates, ACV (Fig. S12) and IPN (Fig. S13), showed that peroxisomal transporter activity did not influence these precursors to PenG. Finally, crude extracts from *N. benthamiana* leaves were tested via paper disk diffusion assays against *Staphylococcus saprophyticus* and *S. epidermidis* demonstrating clear zones of inhibition (Fig. S37).

**Fig 4:**
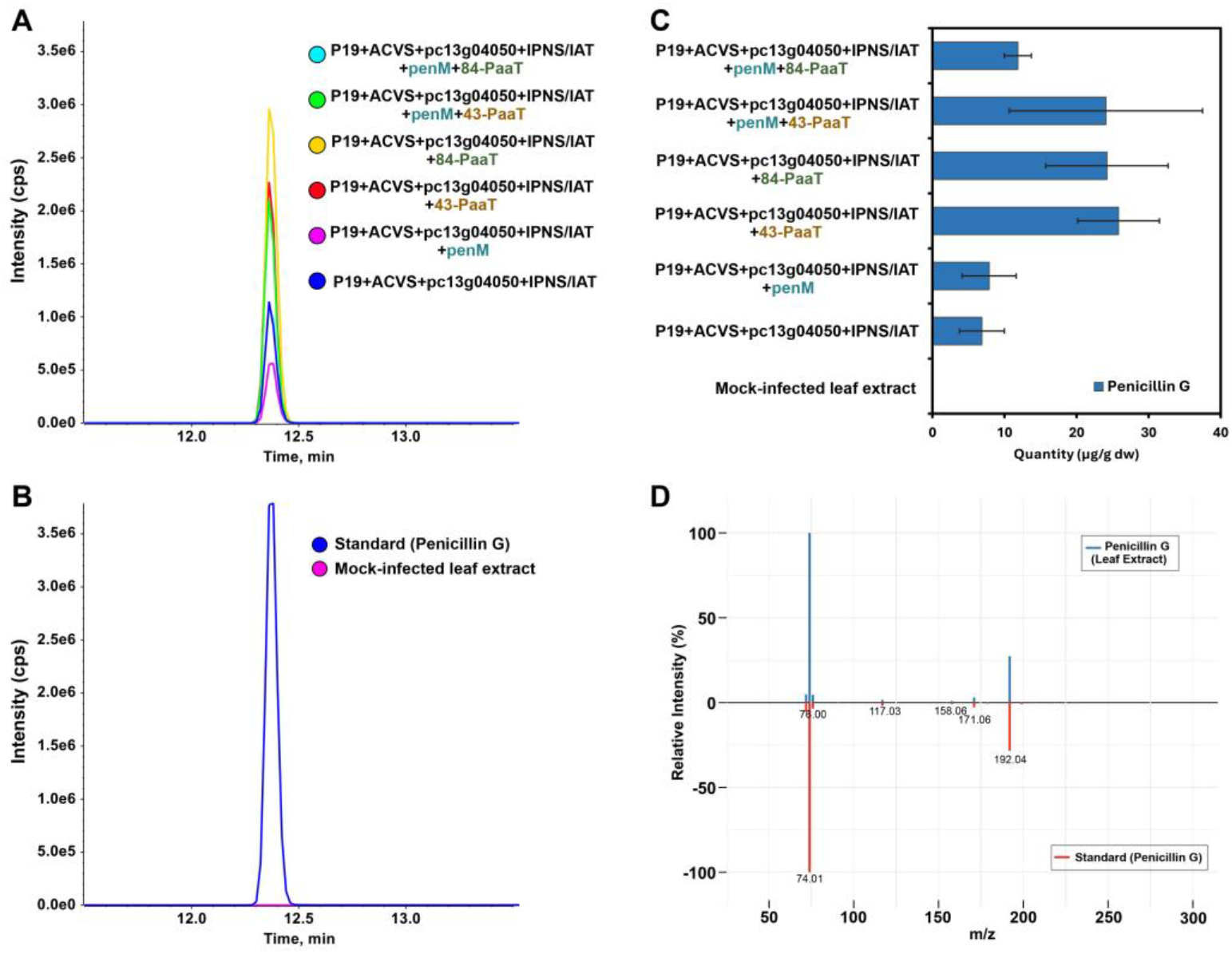
Endogenous peroxisomal transporters can support PenG production in plants, but fungal PaaT enhances accumulation of Pen G. LC-MS/MS-based detection and quantification of penicillin G in *N. benthamiana* leaf extracts expressing the penicillin G biosynthetic genes with and without transporters. (A) Extracted ion chromatograms (XICs) of penicillin G at a retention time (RT) of 12.4 min, detected as [M–H]^−^ at *m/z* 333.091 ± 0.005, from leaves transiently expressing the pathway genes either alone or in combination with the PenM and PaaT transporters, (B) XICs from mock-infiltrated leaf extract (negative control) and an external penicillin G standard, confirming RT and mass accuracy of penicillin G signal detected *in planta*, (C) Bar plot showing absolute accumulation of penicillin G (in µg g^−1^ dry weight) across sample groups shown in panel A and determined based on an external calibration curve (mean ± SD; *n* = 3; Fig. S14), and (D) MS/MS mirror plot comparing fragmentation patterns of penicillin G detected in planta with those of an external penicillin G standard, confirming structural identity of of penicillin G produced in *N. benthamiana*. [Note:84-PaaT, C-terminal GFP tag: 43-PaaT, N-terminal GFP tag; SD: standard deviation].

### Identification of plant defense / response compounds

To investigate *N. benthamiana*’s metabolic response to PenG, LC-MS/MS-based untargeted metabolomic profiling was performed on *N. benthamiana* leaf tissues infiltrated with 1 mg mL^−1^ PenG, water-infiltrated mock (MockH), and non-infiltrated healthy leaves (HL) collected at 1 and 3 days post-infiltration (D1 and D3). Comprehensive metabolite profiling, combining orthogonal separation techniques, hydrophilic interaction liquid chromatography (HILIC) and reverse-phase (RP) chromatography performed in both positive and negative ionization modes, identified a total of 2,254 high-confidence metabolic features, including 1,187 RP-derived and 1,067 HILIC-derived features (Data S1 and S2). Multivariate analysis using Partial Least Squares-Discriminant Analysis (PLS-DA) revealed clear separation between PenG-infiltrated samples and controls at both time points, with the first two principal components explaining 43.7% of the total variance. This result indicated that PenG induces a distinct metabolic shift in *N. benthamiana* (Fig. 5A). Of the 2,254 detected metabolic features, 420 were annotated, corresponding to 269 unique KEGG identifiers (Data S1). KEGG-based functional classification showed that approximately 8.3-9.1% of annotated metabolites were associated with carbohydrate and energy metabolism, while 32.6% were assigned to secondary metabolite biosynthesis (16.3%) and amino-acid metabolism (16.3%; Fig. S15A).

**Fig. 5:**
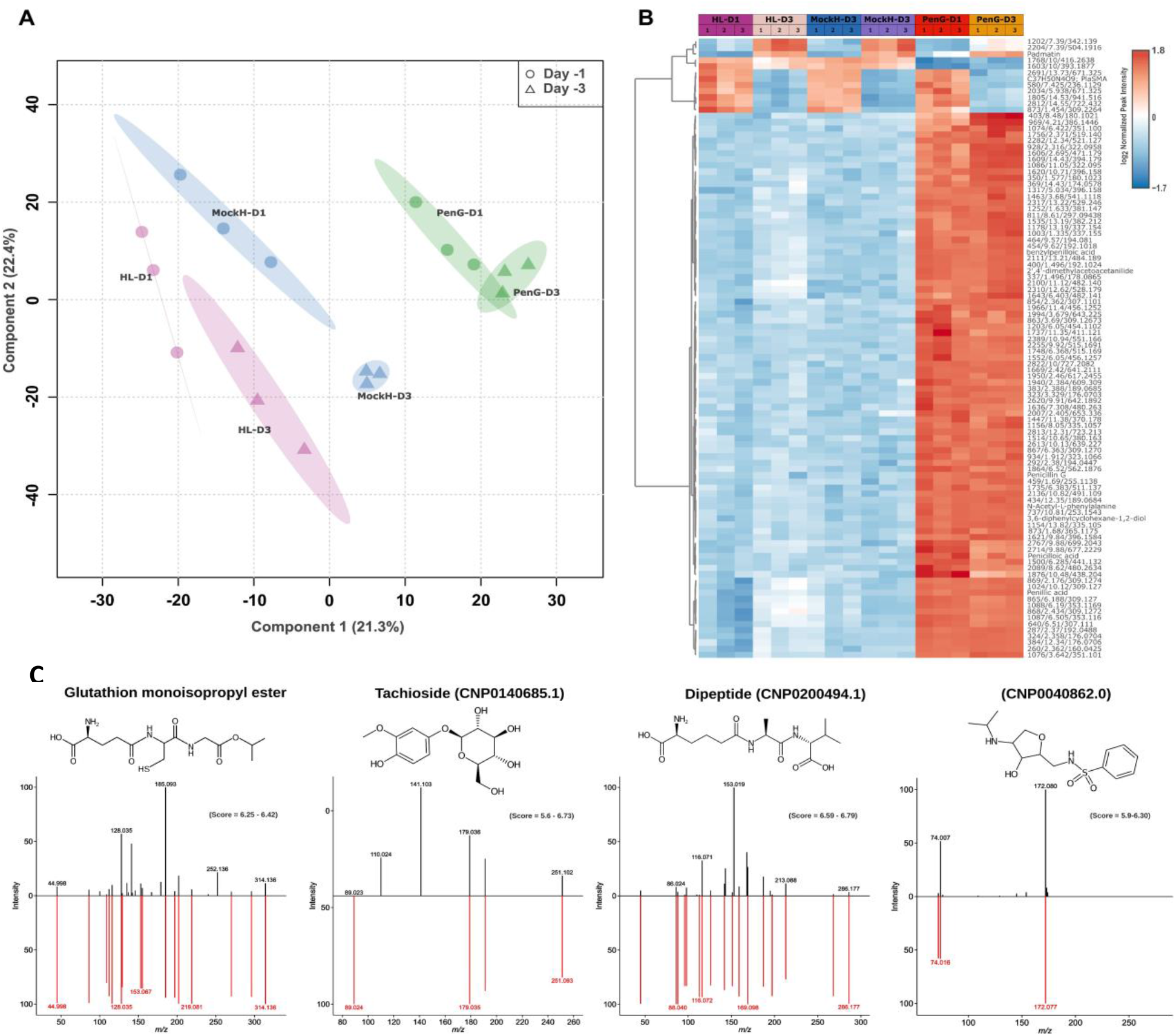
Plant response compounds identified by non-targeted metabolomics. (A) Partial least-square discriminant analysis (PLS-DA) between total metabolome of each sample group analyzed by HILIC and reverse-phase LC-MS/MS in both negative and positive ionization mode. Sample grouping showed distinct metabolic profile of samples infiltrated with penicillin G compared to samples mock and healthy leaves samples. Each dot in the plot represent metabolome of an individual sample within each group, and the shaded regions represent 95% confidence intervals (n=3); (B) heatmap showing top 100 differentially accumulated metabolites with distinct metabolic profile in penicillin G infiltrated leaf samples compared to mock and healthy leaf samples, and (C) structures of plant response compounds identified in leaf extracts after infiltration of penicillin G biosynthetic enzymes and corresponding mirror plots. [Abbreviations: HL = healthy leaf (no infiltration), Mock-H = infiltrated with water, PenG = infiltrated with penicillin G and D1, D3 = Day 1 and 3 of post infiltration].

Differential analysis identified 184 significantly differentially accumulated metabolites (DAMs) in at least one group comparison, based on thresholds of log_2_ fold change (log_2_FC) ≥ ±1, *p*-value, and FDR ≤ 0.01. Most DAMs showed increased accumulation in PenG-infiltrated samples compared to MockH or HL controls (Fig. 5B, Fig. S15B, and Data S3). Notably, the majority of DAMs corresponded to unknown or unannotated features (Fig. 5B), whereas only limited changes were observed in metabolites associated with plant defense or xenobiotic-like stress responses. For instance, N-acetyl-L-phenylalanine exhibited significantly higher levels (log_2_FC=7.9-8.8) in PenG-infiltrated samples compared to MockH and HL controls. Similarly, N-benzylformamide (log_2_FC=4.6-5.6) showed increased accumulation at D1 and D3 in PenG-infiltrated relative to controls. In addition, defense- or stress-related metabolites, such as N-acetyl-L-tyrosine, L-histidine, L-arginine, and 1-methyladenosine, showed approximately 2-fold higher levels specifically at D3 (PenG-D3 vs. MockH-D3) (Data S3).

In contrast to PenG infiltration, *N. benthamiana* leaves transiently expressing genes from the fungal PenG biosynthetic pathway accumulated a distinct subset of host metabolites absent from mock controls (Fig. 5C). *In silico* annotation using MS-Finder, supported by the COCONUT database (*29*), suggested that these metabolites belong to diverse classes of secondary compounds with antioxidant- and defense-related functions. These included the oligopeptide glutathione mono-isopropyl ester (MW 349.41), the phenolic glycoside tachioside (MW 302.10; CNP0140685.1), and the phenylalanine derived alkaloid N-[[3-hydroxy-4-(isopropylamino)tetrahydrofuran-2-yl]methyl]benzenesulfonamide (CNP0040862.0; MW 314.13). In addition, an unannotated dipeptide (CNP02000494.1; MW 331.17) was detected. All of these metabolites were exclusively observed in samples expressing the PenG pathway and were not detected in mock controls (Fig. 5C).

## Discussion

Our study demonstrates the first complete biosynthesis of penicillin G in a plant host *via* recombinant production of fungal proteins including a large (> 400 kDa) multi-functional NRPS enzyme and up to six additional biosynthetic proteins ranging from ∼ 37 – 60 kDa. These results are significant due to future biomedical applications, but also due to the expanded application of *N. benthamiana* in recombinant protein production and natural product biochemistry. To the best of our knowledge plants do not synthesize peptides using NRPS enzymes and instead utilize ribosomally synthesized and post-translationally modified peptides (RiPPs) (*30*).

Furthermore, our work demonstrates that fungal proteins localize to the expected sub-cellular compartment without manipulation of signal peptide sequences. The biosynthetic enzymes appear to interact efficiently based on only minor quantities of intermediates accumulating (Figs. S6C and S7C). Additionally, neither the fungal biosynthetic enzymes, transporters, or resulting peptide products are completely degraded by plant proteases, although future work should establish if quantities of bioactive molecules can be increased by silencing or knocking-out native proteases. An additional significant observation is that enzymes conserved across kingdoms *e*.*g*. PPTases (to some extent), metabolite transporters and aromatic acyl-CoA ligases appear to be capable of substituting for one another, opening up the possibility of engineering non-natural bioactive molecules by combining enzymes from different kingdoms to expand known chemical space *in planta*. The PenG extracted from infiltrated leaves retained bioactivity against gram-positive bacterial pathogens (Fig. S37), demonstrating a practical application of the platform. Furthermore, codon optimization of gene sequences was not always necessary, with some of the codon-optimized genes causing toxic-effects when being propagated through *E. coli*, potentially simplifying future applications.

We achieved maximum PenG yields of 25 μg g ^**−1**^ by including the transporter PaaT with all biosynthetic genes (Fig. 4C) which had more effect than including the CoA-ligase PCL (Fig. 3C). Although modest, these yields are comparable to the original yields of penicillin produced by filamentous fungi (*15*) and even the yields produced by engineered *Saccharomyces cerevisiae* (*25*). PenG degradation products were not detected indicating that the β-lactam ring is not hydrolyzed *in planta*. The yields we attained may be expected when using a heterologous host from a different kingdom which does not produce comparable biosynthetic enzymes (*30-32*). Although unlikely to supplant current fermentation production methods, we envision this plant-based methodology as a blueprint towards the supply of many additional fungal-derived medicines in an alternative, scalable, more environmentally friendly format than by energy intensive fermentation infrastructure. By utilizing existing technologies such as vacuum infiltration (*33*) and developing approaches to store bioactive molecules within plant cells, the efficiency and scale of production of fungal-derived metabolites may be further expanded.

Although PenG has no known molecular target in plants, both direct infiltration and endogenous biosynthesis elicited measurable metabolic responses in *N. benthamiana*. Direct infiltration of PenG triggered plant defense- and stress-related metabolic responses (Fig. 5B; Data S1). The increased accumulation of *N*-acetyl-L-phenylalanine suggests enhanced phenylalanine turnover and a reprogramming of aromatic amino acid metabolism (Fig. 5B; Data S3). In plants, such responses are linked to phenylpropanoid-associated defense pathways and stress-induced metabolic reallocation (*34,35*). Similarly, higher levels of *N*-benzylformamide following PenG-infiltration (Fig. 5B; Data S3) suggest diversion of aromatic amino acid-derived intermediates toward secondary metabolite biosynthesis, consistent with stress-associated metabolic remodeling previously described in cotton (*36*). The modest increases in additional amino acid-related metabolites, including L-histidine, L-arginine, and *N*-acetyl-L-tyrosine, further support a broader stress-associated adjustment of nitrogen and amino acid metabolism (Data S3). L-histidine and L-arginine have previously been reported to play roles in plant stress and defense responses, including signaling and nitric oxide-mediated pathways (*37-39*). In *Arabidopsis*, the modified nucleoside 1-methyladenosine has been shown to regulate stress tolerance through modulation of mRNA stability (*40*). Its increased accumulation following PenG infiltration in *N. benthamiana* may therefore reflect activation of stress-associated post-transcriptional regulatory processes. Importantly, these metabolic changes occurred without major perturbation of central metabolism, suggesting selective activation of defense-associated pathways.

In contrast, transient expression of the fungal PenG biosynthetic pathway led to the accumulation of metabolites not detected in mock controls, including glutathione mono-isopropyl ester, phenolic glycosides, and phenylalanine-derived alkaloid-like compounds (Fig. 5C). Our results indicate that these compounds are not a result of the *Agrobacterium* strains or infiltration process but either the biosynthetic enzymes or bioactive molecules themselves since they were not reported in similar metabolomic studies of *N. benthamiana* after *Agrobacterium*-mediated transformation. The presence of glutathione-related metabolites suggests enhanced redox buffering and detoxification capacity, a characteristic feature of intracellular xenobiotic or biosynthetic stress (*41*). Phenolic glycosides and other aromatic secondary metabolites are widely associated with cellular protection, sequestration, and stress tolerance in plants (*42*). Understanding these metabolic fluxes and defense responses is crucial for developing *N. benthamiana* as a safe and non-toxic PMP platform in future. Collectively, these findings highlight the considerable metabolic plasticity of *N. benthamiana* and support its suitability as a resilient platform for biosynthesis of fungal-derived natural products.

In summary, we have reconstructed the complete biosynthetic pathway for penicillin G in a plant host, demonstrating that fungal enzymes are naturally sorted and located to the correct sub-cellular compartments without engineering signal peptide sequences or post-translational modification sites. Our findings provide a blueprint towards large-scale production of fungal-derived PMPs, which could also be expanded to include agricultural products, and offer an alternative method for producing high-value bioactive molecules from fungi.

## Supporting information

Auxilliary Supplemental Information

## Acknowledgments

We would like to thank Christophe Cocuron from UNT’s BioAnalytical Facility for HRMS analysis, Doug Whitten at Michigan State University’s Proteomics Facility for peptide sequencing analysis, and Dr. Syeda Alam from UNT for providing *Staphylococcus* strains.

## Author contributions

Conceptualization: ES, KDC

Methodology: ES, KDC, APA, SR, GA, AR, YTL

Investigation: ES, KDC, APA, SR, GA, AR, YTL, SJ, YC, DS, PW, HD, SM

Visualization:

Funding acquisition: ES, KDC, MC, APA

Project administration: ES, KDC

Supervision: ES, KDC, APA

Writing – original draft: ES, KDC

Writing – review & editing: ES, KDC, APA, AR, YTL, SR, GA, SJ, YC, PW

## Data and materials availability

All data are available in the main text or the supplementary materials

