## Supplementary Information for "Complete biosynthesis of penicillin G in *Nicotiana benthamiana*"

**The PDF file includes:**

Materials and Methods
Figs. S1 to S38
Tables S1 to S8
References 43 - 54

**Materials and Methods**

Materials purchased and suppliers

Media ingredients, supplements, antibiotics, reagents, and analytical grade chemicals were purchased from Sigma Aldrich, Thermo Fisher Scientific, PhytoTech. The solvents used for HPLC were LCMS grade. General molecular biology procedures were performed as standard and molecular biology kits were used according to the manufacturer's protocols (*e.g.* QIAGEN, NEB, Zymo, Invitrogen). Analytical PCR was performed using OneTaq polymerase and preparative PCR for cloning procedures was performed using Q5 polymerase manufactured by NEB. Restriction endonucleases were also purchased from NEB. Plasmids and PCR products were sequenced using the Plasmidsaurus sequencing service. Chemical reference standards were purchased from MedChem Express, TCI America, Olchemim, Cayman Chemicals and Benchchem.

Organisms used in this study and their suppliers are listed in Table S1; media used for their cultivation is listed in Table S2. Oligonucleotides were purchased from IDT (Table S3), synthetic genes (Table S4) and gene fragments (Table S5) were purchased from TWIST Bioscience or Gene Universal. A complete description of plasmids used in this study is listed in Table S6 and plasmid maps are provided in Figures S16 – S26.

Cultivation conditions

Culture media and conditions used for microbes used in this study are described in Table S1 and Table S2.

***N. benthamiana (for Agrobacterium infiltration and transformation)***: *N. benthamiana* plants used in transient expression experiments to generate Penicillin G and its upstream metabolites were germinated and grown in soil 28 °C under 16-h/8-h day-night cycles. The first 3 fully expanded leaves from the top of 4 to 5-week-old *N. benthamiana* plants were infiltrated with the *Agrobacterium tumefaciens* strain GV3101 harboring the appropriate binary vectors. After infiltration, plants were incubated at 22-24 °C under 16-h/8-h day-night cycles. Infiltrated leaves were collected 2-6 days post inoculation depending on experiments.

***N. benthamiana growth conditions and sample preparation for untargeted metabolomics in response to PenG***

*N. benthamiana* seeds were germinated on a sterile filter paper soaked with water. After one week, the germinated seeds were transferred to pots containing autoclaved coco-peat and vermiculite (3:1) mixture supplied with fertilizer Osmocot (~1.5 mL per pot). The plants were kept and grown in growth chamber at 22-24 °C with 16/8 h day/light cycle and 65% of relative humidity (RH). After 3-4 weeks, fully expanded first 3 leaves were infiltrated with 1 mg/mL of solvent (water as mock), and 1 mg/mL of Penicillin G. The infiltrated leaves were collected at 1 and 3 days of post infiltration (DPI). The fully expanded healthy leaves (HL) were also collected from plants with no infiltration, as control. At least three leaves from each biological replicate (n=3) were collected. The collected leaf samples were immediately frozen in liquid nitrogen and lyophilized for two days at -80 °C. The lyophilized leaves were homogenously ground to a fine powder using the 2010 Geno/Grinder (SPEX SamplePrep LLC, NJ, USA) with two 8 mm stainless steel beads for sample disruption. The dried leaf samples were subsequently used for penicillin G quantification and untargeted metabolomics analysis using ultra-high performance liquid chromatography (UHPLC) with high resolution mass spectrometer (HRMS).

Chemical extraction methods

For UHPLC-HRMS analysis, metabolites were extracted using a chloroform:methanol:water (CMW)–based protocol adapted from previously published methods (*43,44*). Briefly, approximately 10–12 mg of homogenized, dry *Nicotiana benthamiana* leaf tissue was transferred to a 2 mL microcentrifuge tube (USA Scientific, FL, USA) and suspended in 1 mL of CMW extraction solvent (1:2.5:1, v/v/v). Then the extraction solvent was supplemented with the following internal standards: lidocaine (0.5 nmol), sulfamethoxazole (2 nmol), reserpine (2.5 nmol), HEPES (5 nmol), acyclovir (5 nmol), and $^{13}C$-glycine (300 nmol). In parallel, extraction blanks (CMW buffer only) and internal standard–only (IS) controls (CMW supplemented with internal standards only) were prepared and processed alongside the biological samples; penicillin G (2 nmol) and/or ACV (13 nmol) of ACV reference standards were spiked exclusively into the internal standard–only (IS) controls. Samples were briefly vortexed to ensure uniform suspension and homogenized using a Thermomixer C (Eppendorf, NY, USA) at 4 °C for 10 min with agitation at 1500 rpm, followed by centrifugation at 17,000 × g for 5 min at 4 °C. The resulting supernatant was transferred to a new 2 mL microcentrifuge tube, after which 400 µL of ice-cold water was added to induce phase separation. The mixture was vortexed for 10 s and centrifuged again under the same conditions. The upper phase was collected into a new 1.5 mL microcentrifuge tube. For parallel analysis of polar and semi-polar metabolites through untargeted approach by LC–MS/MS, the collected upper phase was divided into two equal aliquots of 500 µL each. [Note that for *N. benthamiana* leaf extracts from transient expression experiments, the upper phase was not split and was entirely used for reverse phase (RP) analysis.] Methanol was removed using a SpeedVac concentrator (ThermoScientific, MA, USA) at 30 °C for 20 min. The samples were subsequently flash-frozen in liquid nitrogen and lyophilized for 3–4 h at -80 °C to remove residual water.

Method for plant extract preparation and the bacterial growth inhibition assay

Infiltrated *N. benthamiana* leaves were collected into a 50 mL Falcon tube and lyophilized for 2 nights. The lyophilized leaf tissue was transferred into homogenization tubes prefilled with 2.7 mm glass beads, and 2 mL of methanol was added. Tissues were disrupted for 30 seconds using a Glen Mills Mini-Beadbeater-16. Homogenized tissue was incubated at 4 °C fridge for 1 hour. The supernatant and leaf debris were separated by centrifugation. The supernatant was transferred to the new tube, and the methanol was evaporated completely under a stream of N2 before being re-suspended in 1 mL of sterile water. The resulting extract was filtered using a 0.22 mm filter prior to the bacterial growth inhibition assay. The two gram-positive bacteria, *Staphylococcus saprophyticus* and *Staphylococcus epidermidis*, were cultured in liquid LB medium, and the starting culture densities were adjusted to optical densities ($OD_{600}$) of 0.44 and 0.35, respectively. A uniform bacterial lawn was prepared by spreading 150 µL on the surface of solid LB medium plates. To evaluate the functional activity of PenG in plant extracts, 25 µL of the plant extracts and known concentrations of penicillin G were pipetted directly onto sterile blank paper disks. These impregnated disks were placed onto the seeded bacterial plates. The plates were incubated at 37 °C for 16 hours. Following incubation, the zones of inhibition were measured and recorded.

Chromatography methods

Metabolomic analyses were performed using an ultra-high-performance liquid chromatography system coupled to a high-resolution mass spectrometer (UHPLC–HRMS; Exion LC system interfaced with a TripleTOF 6600+, AB Sciex, MA, USA), following an established method (*43*). Chromatographic separations were carried out using either a reverse-phase (RP) column (Kinetex F5, 150 × 2.1 mm, 2.6 µm, with F5 guard column, 10 × 2.1 mm; Phenomenex, Torrance, CA, USA) or a hydrophilic interaction liquid chromatography (HILIC) column (ACQUITY Premier BEH Amide, 150 × 2.1 mm, 1.7 µm, with matching VanGuard FIT pre-column, 5 × 2.1 mm, 1.7 µm; Waters, Milford, MA, USA). The autosampler temperature was maintained at 10 °C for all analyses, while column temperatures were set to 20 °C for RP and 35 °C for HILIC separations. A second set of internal standards was prepared separately for chromatographic reconstitution and consisted of *trans*-zeatin-$d_5$ (1 nmol), MOPS (2 nmol), ampicillin (2 nmol), 9-phenanthrol (2 nmol), 5-fluorocytosine (10 nmol), $^{13}$C-benzoic acid (10 nmol), *trans*-cinnamic acid-$d_6$ (10 nmol), and $^{13}$C-fumarate (50 nmol). These standards were dissolved in two different solvent systems: methanol:water (20:80, v/v) for reverse-phase (RP) chromatography and acetonitrile:water (80:20, v/v) for hydrophilic interaction liquid chromatography (HILIC). For each upper phase extract, one dried aliquot was reconstituted in 100 µL of methanol:water (20:80, v/v) for reverse-phase (RP) analysis, while the second aliquot was reconstituted in 100 µL of acetonitrile:water (80:20, v/v) for HILIC. Both biological sample extracts and IS controls were reconstituted using the respective solvent systems containing the corresponding second set of internal standards, whereas extraction blanks were resuspended using the same solvent without internal standards. Reconstituted extracts were sonicated for 5 min, incubated in a thermomixer at 1750 rpm for 5 min, and centrifuged at 17,000 × g for 5 min at 25 °C. Subsequently, 100 µL of the supernatant was transferred to LC–MS/MS glass vial containing an insert (Agilent, CA, USA).

Quality control (QC) samples were prepared by pooling 2 µL from each biological sample into a separate LC–MS/MS vial. Each sample extract, QC, and corresponding blank was injected into the LC–MS/MS system equipped with RP and HILIC columns, with injection volumes of 3 µL and 4 µL, respectively. *N. benthamiana* leaf extracts from transient experiments were only prepared for RP LC-MS/MS analysis, while samples infiltrated with penicillin G compounds were prepared for untargeted metabolomics using RP & HILIC for LC-MS/MS analysis.

For RP chromatography, metabolite separation was achieved at a flow rate of 0.2 mL/min using a gradient system composed of solvent A (0.25% v/v formic acid and 5 mM ammonium formate in water) and solvent B (acetonitrile containing 0.1% formic acid, v/v). The RP gradient profile consisted of an initial hold with 0% B for 2 min, followed by linear increase (0-80%) of B over next 16 min and then rapidly increase to 95% B at 18.1 min and maintained until 21 min, after which the B was decreased to 0% at 21.1 min and column returns to starting conditions and re-equilibrated until 25 min with 0% B.

HILIC separations were conducted at a flow rate of 0.3 mL/min using gradient system consisting of solvent A (0.2% v/v formic acid and 25 mM ammonium formate in water) and solvent B (90% aqueous acetonitrile containing 0.15% v/v formic acid and 10 mM ammonium formate). The HILIC gradient began with 100% B for 2 min, followed by a gradual decrease from 100-70% B at 6 min, 40% B at 9.35 min, and 30% B at 11 min. The gradient was subsequently returned to 100% B by 13.5 min and held until 20 min to ensure adequate column re-equilibration.

Following chromatographic separation, data-independent acquisition was performed using Sequential Window Acquisition of All Theoretical Mass Spectra (SWATH-MS) (*43*). To determine precursor ion isolation windows, preliminary data-dependent acquisition (DDA) experiments were conducted on pooled QC samples in both positive and negative ionization modes for RP and HILIC analyses. Based on these data, 36 variable SWATH windows were defined to cover a precursor ion mass range of m/z 50–1250, with every scan accumulation time of 200 ms per window. Fragment ion spectra were acquired over an *m/z* range of 30–1250 using an accumulation time of 25 ms per window, resulting in a total cycling time of approximately 1.15 s. Electrospray ionization (ESI) source and quadrupole parameters were optimized to ensure consistent ionization and efficient ion transmission across the mass-spectra analyzer. Curtain gas pressure was maintained at 35 psi for all analyses. Source temperatures were set to 550 °C for RP and 600 °C for HILIC separations. IonSpray voltages were adjusted according to ionization polarity and chromatographic mode, ranging from −4500 V (RP) to −3500 V (HILIC) in negative mode and from 3500 V (HILIC) to 5000 V (RP) in positive mode. Nebulizing and heating gas pressures were operated within ranges of 50-60 psi and 60-80 psi, respectively. Declustering potential and collision energy were set to ±50 V and ±30 V, respectively, with a collision energy spread of 10 V applied across all modes to improve fragmentation consistency. To account for potential mass drift during LC-MS/MS runs, an atmospheric-pressure chemical ionization (APCI) calibrant solution was applied after every eight injections, alternating between positive and negative ionization modes. Instrument control and data acquisition were performed using Analyst TF software (v1.8.1; AB Sciex, MA, USA).

Metabolite quantification methods

To quantify penicillin G and ACV from *N. benthamiana* leaf extracts following transient expression, the external calibration curves of penicillin G and ACV were generated using their reference standards prepared at four different concentrations (Fig. S10 & S14). Standard solutions were analyzed using RP LC-MS/MS analysis at negative ESI mode under the same analytical conditions as the samples. Calibration curves were generated by linear regression of peak areas versus concentration. 9-phenanthrol was used as an internal standard across all the samples to correct for analytical variability.

**# Eq. 1:** $$Correction\ factor\ (CF) = \frac{Peak\ Area\ 9 - phenanthrol_{in\ sample}}{Average\ Peak\ Area\ 9 - phenanthrol_{in\ IS\ samples}}$$

**# Eq. 2:** $$Corrected\ Peak\ Area\ (CPA) = \frac{PeakArea_{of\ analyte\ in\ sample}}{CF_{of\ analyte\ in\ Sample}}$$

**# Eq. 3:** $$Quantity\ (\mu g) = \frac{CPA}{Slope}$$

**# Eq. 4:** $$\mu g\ per\ g\ (dry\ weight) = \frac{Quantity\ (\mu g)}{sample\ dry\ weight\ (gram)}$$

UHPLC-HRMS Data Processing

The raw data files generated with Analyst TF (v1.8.1; *.wiff) were imported and processed using MS-DIAL (v5.2.240424.3) (*45*) with SWATH as acquisition method. The mass tolerance was set to 0.01 Dalton (Da; MS1) and 0.025 Da (MS2) and retention time (RT) ranges were defined according to chromatographic mode (HILIC: 1-20 min; RP: 1-18 min). Mass scan ranges were *m/z* 50-1250 for MS1 and *m/z* 30-1250 for MS/MS. Retention time correction was performed using an internal standard. For peak detection, minimum peak height of 1000, mass slice width of 0.1, and linear weighted moving-average smoothing as level 3, were used. Peaks spanning < 5 scans were excluded, and spectral deconvolution was performed with sigma window of 0.5 with exclusion of signals following the precursor ion. The isotopic ions were retained up to 5, while MS/MS abundance was kept at zero. The metabolite annotation was performed using an in-house spectral library from BioAnalytical Facility (BAF) at University of North Texas, MassBank of North America (MoNA; https://mona.fiehnlab.ucdavis.edu/), the Global Natural Product Social Molecular Networking (GNPS; https://gnps.ucsd.edu/), and the RIKEN MSn spectral database for phytochemical (ReSpect; http://spectra.psc.riken.jp/).

Identification criteria included mass tolerances of 0.01 Da (MS1) and 0.025 Da (MS2), RT tolerances of 1 min (in-house) and 100 min (public libraries), and direct, weighted, and reverse dot product scores of 0.5. Annotations were accepted when at least one reference spectrum matched, with matched peak ratios ≥ 20% (in-house) or ≥ 30% (public). RT was incorporated into scoring for in-house library. For positive mode adducts $[M+H]^+$, $[M+Na]^+$, $[M+K]^+$, $[M+H-H_2O]^+$, $[M+H-2H_2O]^+$, $[2M+H]^+$ were selected, while for negative mode the selected adducts were $[M-H]^-$, $[M-H_2O-H]^-$, $[M+Na-2H]^-$, $[M+FA-H]^-$, $[2M-H]^-$, $[2M+FA-H]^-$. Peak/feature alignment was performed using a QC sample injected mid-sequence, with tolerances of 0.1 min (RT) and 0.015 Da (MS1). Features with sample max and blank mean ratio < 5 or lacking MS/MS spectra were removed. Post alignment, the data curation was performed using MS-CleanR module in MS-DIAL and ghost peaks, peaks with incorrect *m/z* values, peaks detected in blanks (blank ratio ≥ 0.15), and peaks with high relative standard deviation (RSD ≥ 25%) were removed (*43,46*). All peak integrations and alignments were manually inspected. Finally, the aligned raw peak data from MS-DIAL was exported into excel files and further processed for manual curation and amalgamation. The normalization was performed using the internal standard 9-phenanthrol. For each feature, the standard ratio (SR) was calculated using the ratio of average normalized peak from IS samples and from all biological. Then features with SR > 0.15, signal-to-noise ratio (S/N) < 10, within-group RSD > 100%, or RSD >25% across biological groups were excluded. A second curation step incorporated annotation status and removed duplicate features based on annotation, annotated features with RSD <100% were retained, while features with total-score < 1 were labeled as unknown features. The normalized peak data were used for downstream analysis.

Extracted-ion Chromatogram and MS/MS Fragmentation Analysis

Fungal metabolites, including δ-(L-α-aminoadipyl)-L-cysteinyl-D-valine (ACV), isopenicillin N, and penicillin G, were assessed in *N. benthamiana* leaf extracts from transient expression experiments, using reverse-phase LC–MS/MS operated in negative ESI mode. Metabolite identification and confirmation was based on extracted-ion chromatograms (XICs) together with MS/MS fragmentation analysis. The XICs for ACV and penicillin G were generated using PeakView (AB Sciex, MA, USA) with a mass extraction window of ±0.005 Da. Compound identities were confirmed by concordance in retention time and accurate mass by comparing the XICs of their corresponding reference standard (ACV/Penicillin G). Additionally, MS/MS spectra of ACV and penicillin G were further compared with the detected spectra from their reference standards, and spectral similarity was visualized using mirror-plots generated with the ggplot2 package (*47*) in R (https://www.r-project.org/). For isopenicillin N, the XIC was detected based on its [M-H]- precursor ion *m/z*, and the corresponding MS/MS spectra were interpreted using *in silico* fragmentation analysis against the fragmentation patterns reported in public databases (PubChem, ChEBI, DrugBank) using MS-FINDER. ACV and penicillin G were also independently evaluated using MS-Finder to provide complementary *in silico* support for metabolite identification and confirmation in *N. benthamiana* leaf extracts. Only *in silico* identified spectra with high (Score 6.5–6.9) or very high confidence (Score ≥7.0) were used to confirm the detected MS/MS spectra of ACV tripeptide, isopenicillin N, and penicillin G in *N. benthamiana* leaf extracts.

*In silico* structural annotation of LC-MS/MS features using MS-Finder

For annotation of unknown features, their respective MS/MS spectral data from each chromatographic and ionization mode were exported as .msp files and analyzed using MS-Finder (v3.61) (*48*). The candidate structure and annotation of MS/MS fragments for each feature *in silico* were searched against the background databases provided in MS-Finder, including HMDB,

YMDB (yeast), PlantCyc, Natural product databases (UNPD, KNApSAcK, COCONUT, NPA, and NANPDP) and biomolecules databases (ChEBI and PubChem). MS-Finder assigns a score (0-10) to each predicted annotations of a feature and annotations with score between 6.0–6.5, >6.5–<7.0, and ≥7.0 were considered as moderate, high and very high confidence, respectively. For untargeted data, the MS-Finder based annotations (name, alignment ID, ontology, formula, InChIKey, and SMILES) were incorporated for unknown features to the data from MS-DIAL using their respective information matched based on alignment ID, *m/z*, and RT in excel.

Statistical Data Analysis

The normalized data was analyzed using MetaboAnalyst (v6.0) (*49*). Before analysis, log10 transformation, autoscaling with no additional normalization parameters were applied for data processing. Multivariate analysis, including principal component analysis (PCA) and Partial Least Squares-Discriminant analysis (PLS-DA), were performed to evaluate overall variability across the different sample groups. Univariate statistical tests, including Student's t-test and one way ANOVA followed by Tukey's HSD post hoc test, were applied to identify metabolites that were differentially accumulated among sample groups.

Plasmid construction methods

For *pcbAB* the gene was reassembled from native or synthetic fragments (~ 2 – 4 kb) using yeast homologous recombination. Smaller genes (< 5kb) were ordered directly in entry vectors and transferred to expression plasmids / destination vectors using LR recombination (Gateway) or GoldenBraid cloning techniques.

***pcbAB* (native) gene assembly:** The *pcbAB* gene was amplified in three overlapping fragments from *P. rubens* cDNA template, omitting the stop codon. The *eGFP* gene was amplified from the pMDC84 vector and used as a fourth fragment, positioned downstream of and in-frame with the *pcbAB* native sequence. Each fragment was designed with 30 bp overlapping regions using primers listed in Table S3 and assembled into the pDEST/URA/Ub10 vector using yeast recombination.

Transformation methods

***E. coli*:** Chemically competent *E. coli* cells (Invitrogen) were thawed on ice. Aliquots of 50 µL were mixed with 1–5 ng of plasmid DNA, gently flicked to mix, and incubated on ice for 30 minutes. Heat shock was performed at 42 °C for 45 seconds, followed by immediate cooling on ice for 5 minutes. SOC medium (250 µL) was added, and the cells were incubated at 37 °C with shaking at 220 RPM for 1 hour. Transformants were plated on LB agar containing the relevant antibiotic and incubated overnight at 37 °C. Colonies were screened by PCR and final constructs were verified by plasmid sequencing

**Yeast:** Chemically competent *Saccharomyces cerevisiae* cells were prepared following the protocol provided with the Zymo Research Yeast Competent Cell Preparation Kit. In brief, a single yeast colony was inoculated into YPD medium and incubated at 30 °C with shaking at 220 RPM until cultures reached mid-log phase (OD 600 ≈ 0.7–1.0). Cells were harvested by centrifugation, washed, and processed according to kit instructions to obtain chemically competent cells. For transformation, equimolar amounts of linear DNA were mixed with 100 µL of competent yeast cells. The mixture was incubated with the supplied transformation reagents and subjected to heat shock as described in the Zymo Research protocol. Transformed cells were plated on selective

SM-Ura agar and incubated at 29 °C for 2–3days to allow colony formation. Individual yeast colonies were inoculated in 10 mL SM-Ura liquid medium at 29 °C overnight. Plasmids were extracted using the Zymoprep Yeast Plasmid Miniprep Kit (Zymo Research) and transformed into chemically competent *E. coli* TOP10 cells (Invitrogen).

**Agrobacterium**: The recombinant plant expression plasmid was diluted approximately to 10 ng/µL, and 1 µL of the diluted plasmid DNA was mixed with 50 µL of competent *Agrobacterium tumefaciens* GV3101 cells in a chilled 1.5 mL microcentrifuge tube, then gently transferred to a pre-chilled electroporation cuvette. Electroporation was performed at 2.4 kV (Eppendorf E-porator). Immediately after the pulse, 950 µL of LB medium was added, and the suspension was gently mixed, and the mixture was transferred to a 2 mL microcentrifuge tube for recovery. Transformed cells were incubated at 28 °C for 1 hour before plating 50 µL onto LB agar containing 50 µg/mL Gentamycin, 25 µg/ml Rifampicin, and the appropriate antibiotic used to select the introduced plasmids. Plates were incubated at 28 °C for 48 hours to allow colony formation.

***Agrobacterium tumefaciens*-mediated transient expression in *N. benthamiana* leaves**:
Agrobacteria harboring the appropriate binary vectors were grown in a 2 mL starter culture of LB supplemented with 50 µg/mL Gentamycin, 25 µg/mL Rifampicin, and the appropriate antibiotics overnight at 28 °C. The next day, a portion of the starter culture was inoculated in LB medium (1: 100 dilution ratio) containing 50 µg/mL Gentamycin, 25 µg/mL Rifampicin, 10 mM MES, 25 µM acetosyringone, and the appropriate antibiotics, and was incubated overnight at 28 °C. Cells were harvested by centrifugation, then resuspended in infiltration buffer containing 10 mM $MgCl_2$, 10 mM MES (pH 5.6), 500 µM acetosyringone, 0.5% glucose, and poloxamer 188 (1%, w/v) dissolved in sterile water, and incubated at 28 °C for 3 hours. Before leaf infiltration, the final $OD_{600}$ of each culture in the infiltration mixtures ranged from 0.2 to 0.6, depending on the experiment. The infiltration procedures were previously described (*50*). For all infiltrations, *A. tumefaciens* harboring the tomato bushy stunt virus gene *P19* was included to suppress gene silencing and enhance transient protein expression in *N. benthamiana* leaves (*51*). The infiltration experiments are summarized in Table S7 and the resultant LCMS chromatograms are shown in Fig. S4 – S8, and S12 – S13.

<u>PCR and RT-PCR methods</u>

***E. coli* colony PCR**: Individual *E. coli* colonies were picked and dipped into the PCR master mix as the template. PCR amplifications were performed using primers specific to the target regions. Cycling conditions included an initial denaturation at 98 °C for 5 min, followed by 30–35 cycles of denaturation, annealing, and extension steps appropriate to the primer–amplicon set. PCR products were analyzed by gel electrophoresis.

**Yeast colony PCR**: Single yeast colonies were picked and resuspended PCR mixture as a template. PCR amplifications were performed using primers specific to the target regions. Cycling conditions included an initial denaturation at 98 °C for 5 min, followed by 30–35 cycles of denaturation, annealing, and extension steps appropriate to the primer–amplicon set. PCR products were analyzed by gel electrophoresis.

***P. rubens* RT-PCR**: Fungal mycelia were harvested by filtration and ground in liquid nitrogen. Total RNA was extracted using the Qiagen RNeasy Mini Kit, following the manufacturer's protocol, including the recommended homogenization step for filamentous fungi. RNA concentration and purity were assessed spectrophotometrically. To eliminate DNA contamination, 5 µg total RNA was digested with 2 µL of TURBO DNase (Thermo Fisher Scientific), incubating at 37 °C for 120 minutes per the supplier's guidelines. The RNA was further purified using the Monarch RNA Cleanup Kit (New England Biolabs) and eluted in RNase-free water. For cDNA synthesis, 1000 ng of DNase-treated RNA was used as template in RT-PCR reactions with the LunaScript® RT SuperMix Kit (New England Biolabs) following the standard protocol, yielding cDNA suitable for downstream applications.

***N. benthamiana* RT-PCR**:
The expression of transgenes in *N.benthamiana* leaf tissue was confirmed at the transcriptional level using qualitative RT-PCR (Fig. S3, S28 – S29). The Agrobacterium-infiltrated and mock-infiltrated leaves were collected and frozen in liquid nitrogen. RNA was extracted using the QIAGEN RNeasy Plant mini kit (QIAGEN) and treated with TURBO DNase (Thermo Fisher Scientific) to eliminate DNA contamination. One microgram of DNase-treated RNA was used to synthesize cDNA using OneTaq RT-PCR kit (New England Biolabs). The synthesized cDNA was diluted to 2-fold. One microliter of the diluted cDNA was used as the template for PCR to confirm the presence of transgene transcripts using Promega GoTaq Green Master Mix (Promega). PCR reactions were set up following the manufacturer's protocol. Cycling conditions included an initial denaturation at 95 °C for 2 min, followed by 30 cycles of denaturation, annealing, and extension steps appropriate to the primer-amplicon set. PCR products were analyzed by gel electrophoresis.

**Gel electrophoresis**: PCR products and DNA fragments were resolved using 0.8% agarose gels prepared in 1× TAE buffer containing SYBR Safe DNA Gel Stain (Thermo Fisher Scientific). Electrophoresis was carried out at 140V until adequate separation was achieved and gels were visualized under UV illumination.

<u>Protein extraction and Western blot assays</u>
Agrobacterium-infiltrated *N. benthamiana* leaves expressing the native *pcbAB* gene were used for protein identification 4 days post-infiltration. For each treatment, 100 mg of leaf tissue was collected, frozen in liquid nitrogen, and ground using a Retsch MM400 Mixer Mill in a 2 mL screw cap microtube. The ground leaf tissue was mixed with 400 µL Urea SDS sample buffer (8 M Urea, 2% SDS, 20% Glycerol, 0.1M Tris-HCl, pH 6.8, 0.004% Bromophenol Blue, 0.1M DTT) and incubated at 95 °C for 10 min. The supernatant was separated from tissue debris by centrifugation and transferred into a new microtube. Fifteen microliters of the supernatant were loaded onto a 10% TGX Stain-Free polyacrylamide gel (Bio-Rad) to perform gel electrophoresis. The gel was run at 50 V for 30 mins to concentrate proteins into a thin line in the stacking gel, and subsequently the voltage was increased to 150 V to separate proteins in the resolving gel until the migration front reached the bottom of the gel. The gel was imaged using ChemiDoc XRS imaging system (Bio-Rad) before transferring proteins. The proteins were transferred to a PVDF membrane by using the Trans-Blot Turbo Transfer System at 25 °C and 2.5 V for 10 min (Bio-Rad). The membrane was blocked in 5% non-fat milk in TBST buffer (0.2 M NaCl, 50 mM Tris pH 7.6, and 0.1% Tween-20) for one hour, and incubated with the anti-GFP rabbit monoclonal antibody (Boster Bio) at a 1:2000 dilution at 4 °C overnight. The second antibody was HRP-conjugated anti-rabbit IgG antibody (Thermo Fisher Scientific) at a 1:10,000 dilution. The incubation was at room temperature for one hour. To visualize proteins on western blots, Clarity Western ECL

Blotting substrate (Bio-Rad) and ChemiDoc XRS imaging system (Bio-Rad) were used, following the manufacturer's protocol (Fig. S29).

**Protein identification**

To prepare protein samples for peptide sequencing to identify the presence of the heterologous ACVS proteins in Agrobacterium-infiltrated *N. benthamiana* leaves, 25 μL of protein extracts prepared as described above were loaded onto a 10% TGX Stain-Free polyacrylamide gel to separate proteins by molecular weight. The gel was stained with Coomassie blue dye (Bio-Rad) following the manufacturer's protocol. The bands of six putative ACVS proteins (see Fig. S30) were sliced with a clean razor blade, transferred into a 1.5 mL microtube containing 5% acetic acid, and stored at 4 °C before shipping. Peptides were analyzed by LC-MS/MS at the Michigan State University Proteomics Facility using standard data-dependent acquisition workflows. The analysis data for the submitted samples were in a *.sf3 file that can be opened with Scaffold Viewer (Proteome Software). The "Proteins" function in the viewer located the identified peptides from the samples on the full-length ACVS protein sequence and showed % coverage information (Fig. S31 – S36).

Confocal microscopy

Confocal fluorescence microscopy was performed using a Zeiss LSM 710 confocal laser-scanning microscope (CLSM; retrofitted with an AIRYSCAN head for enhanced resolution) to visualize the subcellular localization of heterologous proteins in Agrobacterium-infiltrated *N. benthamiana* leaves. Images were obtained with a 63x water immersion objective lens (Numerical Aperture = 1.0). Images of leaf cells were collected as Z-stacks with 15-26 optical sections taken 0.5 μm apart and saved as 1,024 x 1,024-pixel digital images. GFP and chlorophyll autofluorescence were excited with a 488 nm laser, and mCherry with a 563 nm laser. Emission spectra of GFP, mCherry, and chlorophyll were collected from 500 to 540 nm, 590-640 nm and 640-720 nm, respectively. Leaf cells expressing a single fluorescence protein (GFP or mCherry) were imaged using the previously described settings to confirm that there was no detectable overlap between different signal collection spectra. Images were taken from at least 2 separate infiltrated leaves across at least 3 individual infiltration experiments. To quantify colocalization of GFP-fused proteins and mCherry, the Coloc2 plugin in ImageJ Fiji (version 2.17.0) was used to evaluate pixel intensity correlation between two fluorophores using two metrics: Pearson's correlation coefficient and Manders' overlap coefficients ($M_1$ and $M_2$), and the statistical significance of the colocalization was evaluated by the Costes statistical significance test. For each treatment, eight images were used to analyze.

**Fig. S1.**

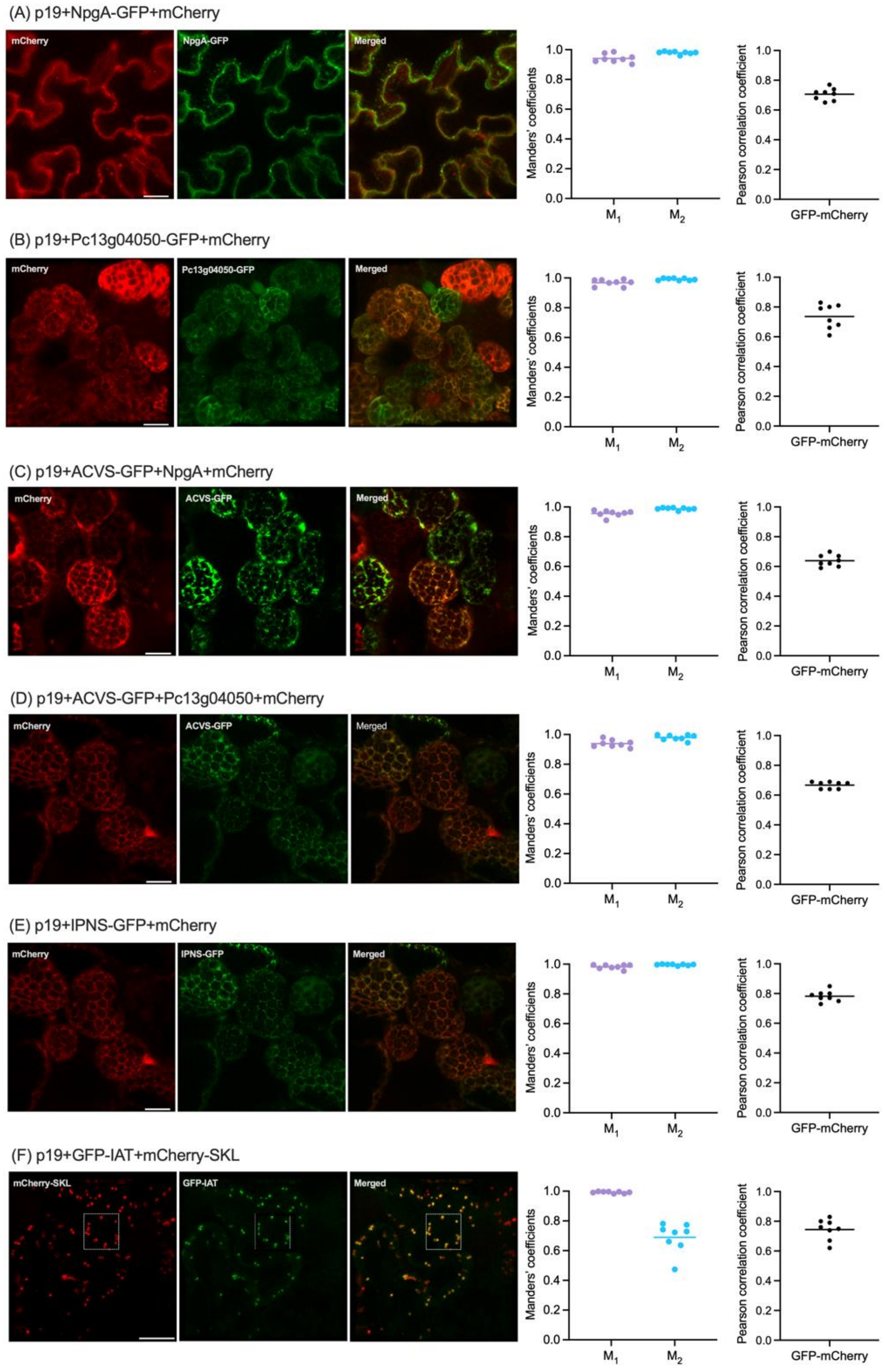

(A) p19+NpgA-GFP+mCherry
mCherry
NpgA-GFP
Merged
Manders' coefficients
Pearson correlation coefficient
1.0
0.8
0.6
0.4
0.2
0.0
M1
M2
GFP-mCherry
(B) p19+Pc13g04050-GFP+mCherry
Pc13g04050-GFP
(C) p19+ACVS-GFP+NpgA+mCherry
ACVS-GFP
(D) p19+ACVS-GFP+Pc13g04050+mCherry
(E) p19+IPNS-GFP+mCherry
IPNS-GFP
(F) p19+GFP-IAT+mCherry-SKL
mCherry-SKL
GFP-IAT

**Colocalization analysis of GFP-fused proteins and mCherry in the infiltrated *N. benthamiana* leaves.** Representative confocal images show the expression and subcellular localization of GFP-fused proteins and mCherry in leaf cells across different treatments. The corresponding merged images are also shown. Manders' overlap coefficients and Pearson's correlation coefficient were used to quantify the colocalization between GFP and mCherry across treatments using Fiji's Coloc2 plugin (n = 8 / treatment), as shown in the graphs. Manders' overlap coefficients: the fraction of the total fluorescence of one channel overlapping with the other. $M_1$: the portion of mCherry overlapped with GFP. $M_2$: the portion of GFP overlapped with mCherry. Pearson's correlation coefficient: the linear correlation of pixel intensities between GFP and mCherry. White dashed boxes indicate the colocalization areas analyzed. SKL, corresponds to serine, lysine, leucine at the C-terminus, the peroxisome targeting signal type 1. Bar = 20 μm.

**Fig. S2.**

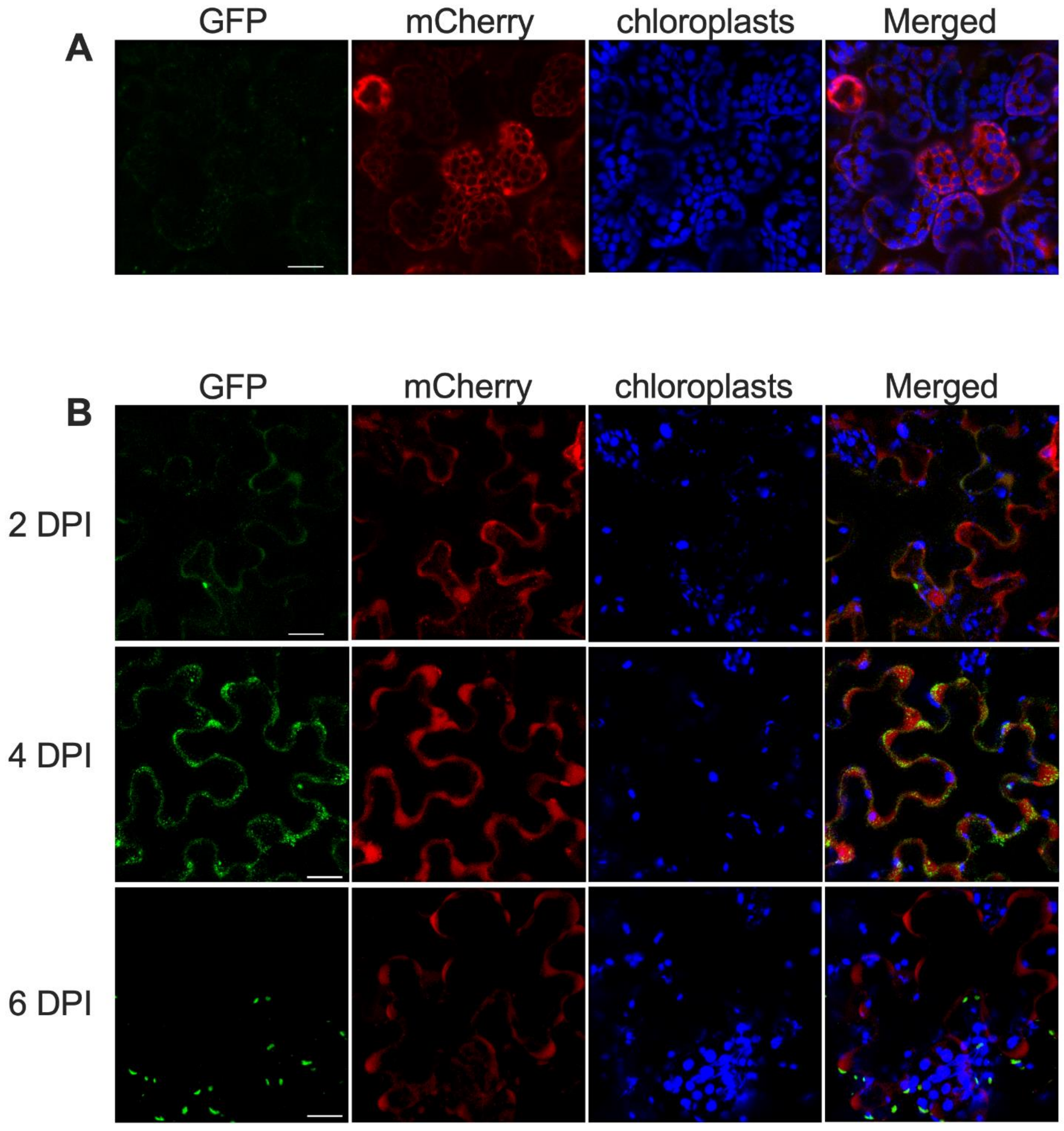

**Subcellular localization of ACVS-GFP enzyme *N. benthamiana*.** (A) Confocal images for subcellular localizations of ACVS-GFP, co-expressed with mCherry in the absence of PPTase. Chloroplasts are false colored blue (chlorophyll fluorescence). The corresponding merged images are also shown in the last panel of the row. (B) Time-course of ACVS-GFP protein signal at 2, 4, and 6 days post infiltration (co-expressed with PPTase, Pc13g04050). Confocal images show the subcellular distribution of ACVS-GFP and mCherry. Chloroplasts are false colored blue (chlorophyll fluorescence). The corresponding merged images for each time point are also shown in the last column of each row. Abbreviations: DPI = days post infiltration. Bar = 20 μm.

**Fig. S3.**

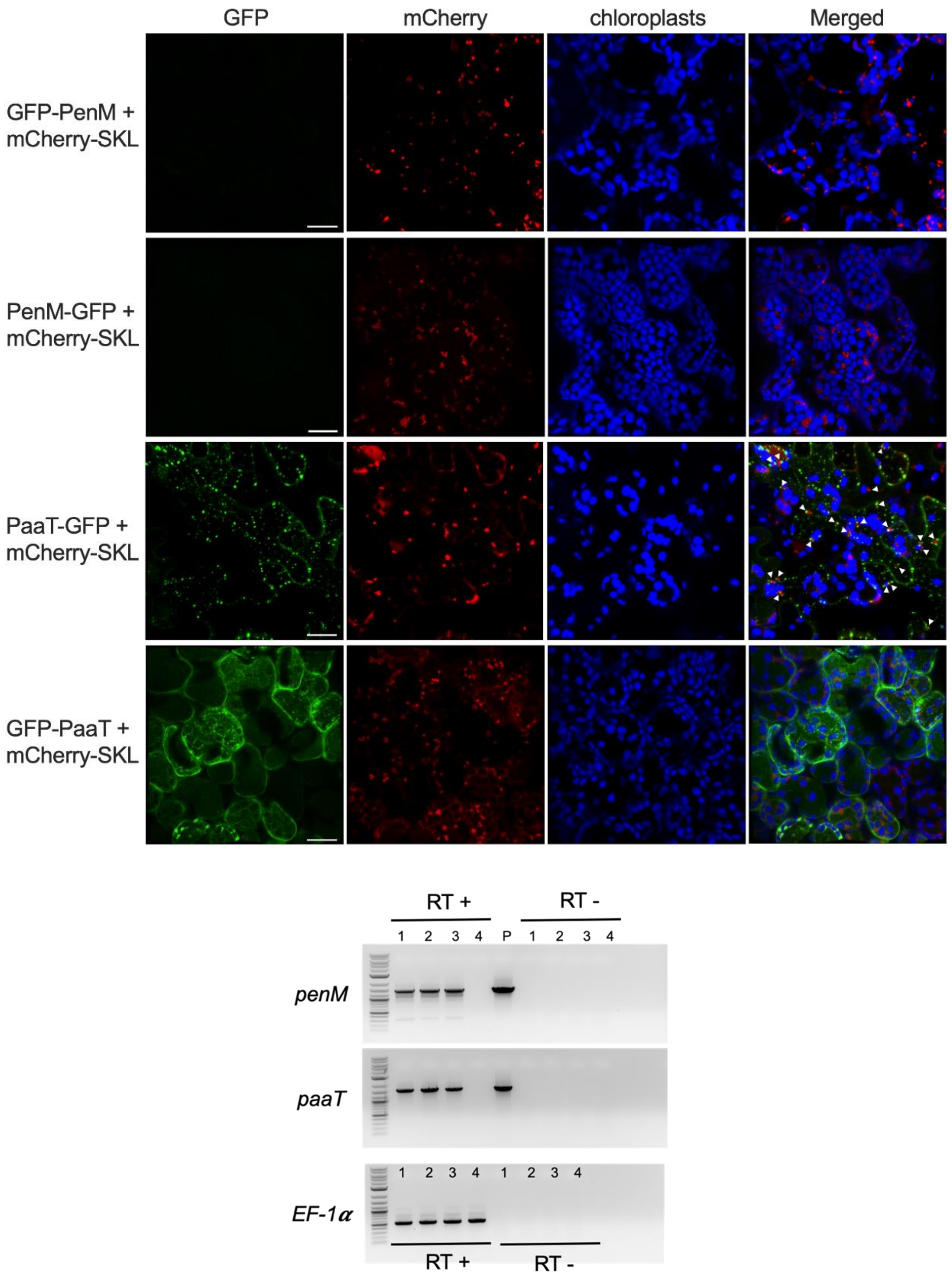

**Production of fungal transporters in *N. benthamiana*.** Top composite: Confocal images for the analysis of GFP-tagged PenM and PaaT transporters co-expressed mCherry-SKL (chloroplast fluorescence is false-colored blue). The corresponding merged images are also shown in the last column. GFP-PenM and GFP-PaaT: the GFP reporters are fused at the N-terminus of proteins. PenM-GFP and Paat-GFP: GFP reporters are fused at the C-terminus. Arrowhead: the regions where PaaT transporters occur on or just adjacent to peroxisomes. SKL is the peroxisomal targeting signal type 1. Bar = 20 μm. Lower composite: Confirmation of expression of the *penM* and *paaT* genes in infiltrated *N. benthamiana* leaves via RT-PCR analysis. Lane 1-3: RNA samples were isolated from leaves infiltrated with *paaT* or *penM* genes, along with the gene silencing suppressor p19; three samples represent RNA from three individual infiltrated leaves. Lane 4: The RNA sample was isolated from the infiltrated leaf with the p19 gene alone. *EF-1$\alpha$* served as the reference gene showing the consistency of RNA isolation and cDNA synthesis across all samples. RT+: RT-dependent cDNAs were used as templates for PCR. RT-: RNAs were used as templates for PCR (no RT) P: plasmids as the positive controls. M: 1 kb DNA ladder.

**Fig. S4.**

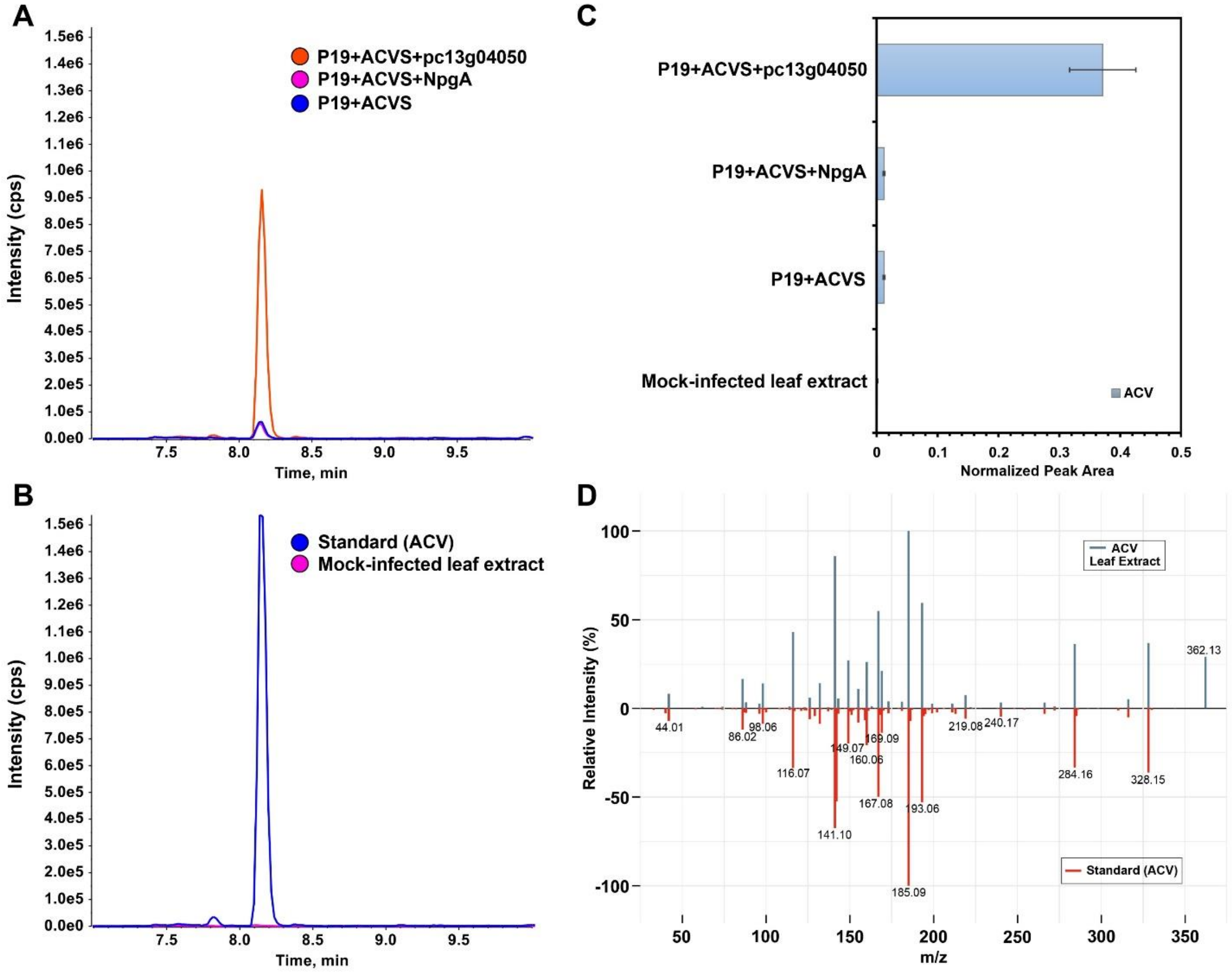

**ACV production without and with PPTases.** Detection and confirmation of *in planta*-produced penicillin biosynthetic pathway intermediate δ-(L-α-aminoadipyl)-L-cysteinyl-D-valine (ACV) using LC-MS/MS analysis. (A) Extracted ion chromatograms (XICs) of ACV at retention time (RT) 8.4 min, detected as $[M\text{-}H]^-$ at *m/z* 362.138±0.005, from *N. benthamiana* leaves transiently expressing the non-ribosomal peptide synthetase ACVS alone or co-expressed with the phosphopantetheinyl transferases (PPTases) NpgA or pc13g04050. (B) XIC comparison of an external ACV standard and a mock-infiltrated leaf extract (negative control), confirming RT and mass accuracy of detected ACV signal *in planta*, (C) Relative accumulation of ACV across the sample groups as shown in panel A, presented as normalized peak area (mean±SD; n=3), (D) Mirror plot comparison of MS/MS fragmentation spectra of ACV detected in leaf extracts (top, blue) and the external ACV standard (bottom, red), demonstrating matching fragment ions and confirming the molecular identity of the produced first-step intermediate. LC-MS/MS analysis was performed using reverse-phase chromatography in negative electrospray ionization (ESI) mode.

**Fig. S5.**

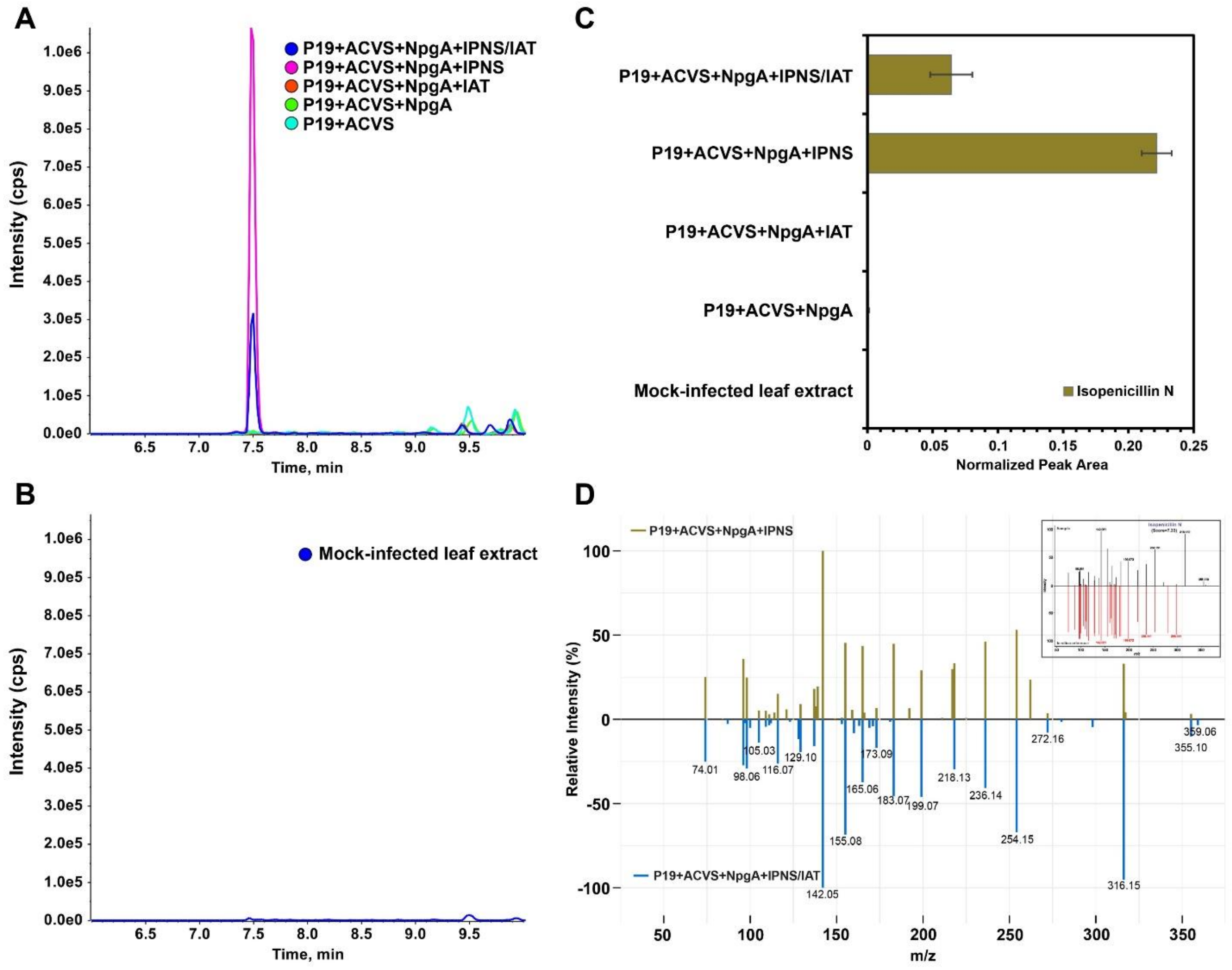

**Isopenicillin N (IPN) synthesis is dependent on co-expression of ACVS and IPNS.** Detection and confirmation of *in planta* biosynthesis IPN in *N. benthamiana* using reverse-phase LC-MS/MS (TripleTOF 6600, negative ESI) analysis. (A) Extracted ion chromatograms (XICs) showing IPN detection at retention time (RT) 7.6–7.7 min, detected as $[M\text{-}H]^-$ at *m/z* 358.107 ± 0.005, exclusively in *N. benthamiana* leaves transiently co-expressing the ACV-forming enzyme ACVS, the phosphopantetheinyl transferase NpgA, and isopenicillin N synthase (IPNS). IPN is not detected upon co-expression with isopenicillin N acyltransferase (IAT), in the absence of IPNS, or when ACVS ± PPTase is expressed alone. (B) XIC of mock-infected leaf extract as negative control with absence of IPN. (C) Bar-plot showing IPNS-dependent accumulation of IPN across sample groups as shown in panel A, presented as normalized peak area (mean ± SD; $n = 3$). (D) Mirror plot showing spectral concordance of IPN detected in plant extracts; inset shows high similarity (score=7.33/10) between *in planta* and *in-silico* predicted MS/MS fragmentation of IPN analyzed using MS-Finder, supporting structural assignment.

**Fig. S6.**

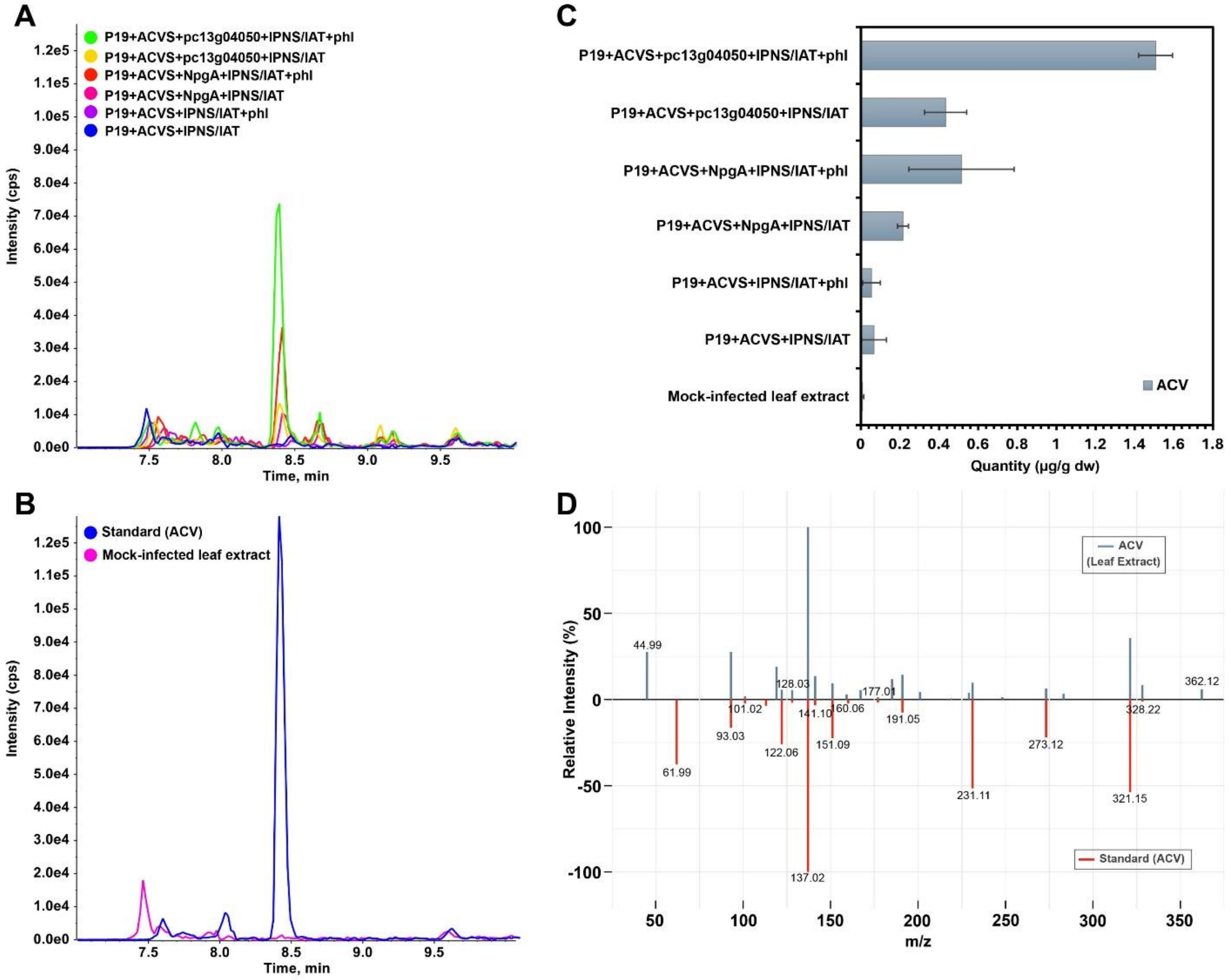

**Accumulation of ACV intermediates when Pen G pathway genes are co-expressed.** Detection and quantification of δ-(L-α-aminoadipyl)-L-cysteinyl-D-valine (ACV) in *N. benthamiana* using reverse-phase LC-MS/MS (TripleTOF 6600, negative ESI). (A) Extracted ion chromatograms (XICs) of ACV at RT 8.4 min, detected as $[M-H]^-$ at *m/z* 362.138 ± 0.005, from leaves expressing different combinations of penicillin G biosynthetic genes, (B) XICs from a mock-infiltrated extract (negative control) and an external ACV standard, confirming retention time and mass accuracy, (C) Absolute ACV quantification (µg $g^{-1}$ dry weight) across the expression conditions shown in panel A, determined using an external calibration curve (mean ± SD, n = 3), and (D) MS/MS mirror plot comparing ACV from leaf extracts with an external standard, showing matching fragment ions that verify ACV identity *in planta*.

**Fig. S7.**

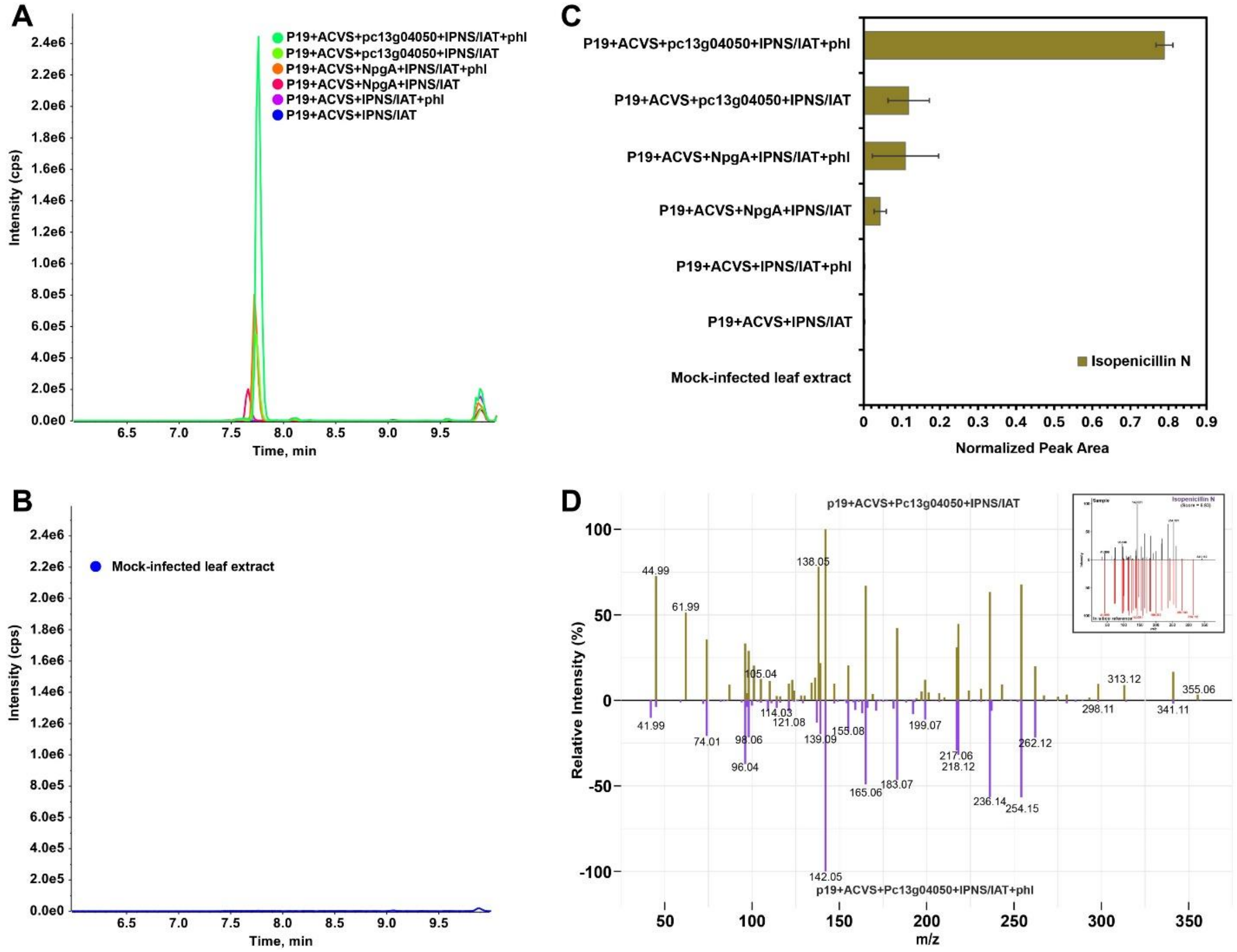

**Accumulation of IPN intermediates when Pen G pathway genes are co-expressed.** Detection and characterization of ispenicillin N in *N. benthamiana* leaf extracts expressing complete penicillin G biosynthetic pathway genes, analyzed by revere-phase LC-MS/MS (TripleTOF 6600+, negative ESI). (A) Extracted ion chromatograms (XICs) of isopenicillin N at retention time (RT) 7.6-7.7 min, detected as $[M–H]^-$ at *m/z* 358.107 ± 0.005, from leaves transiently expressing different combinations of pathway genes; (B) XICs from a mock-infiltrated leaves extract (negative control), showing no detectable signal at the mass or RT of isopenicillin N, (C) Relative accumulation of isopenicillin across the samples groups as shown in panel A, presented as normalized peak area (mean ± SD; n = 3), and (D) MS/MS spectra of isopenicillin N obtained from representative samples, shown as a mirror plot. Fragment ions matched with *in silico* fragmentation pattern predicted by MS-Finder (inset; score=6.83/10), supporting structural assignment.

**Fig. S8.**

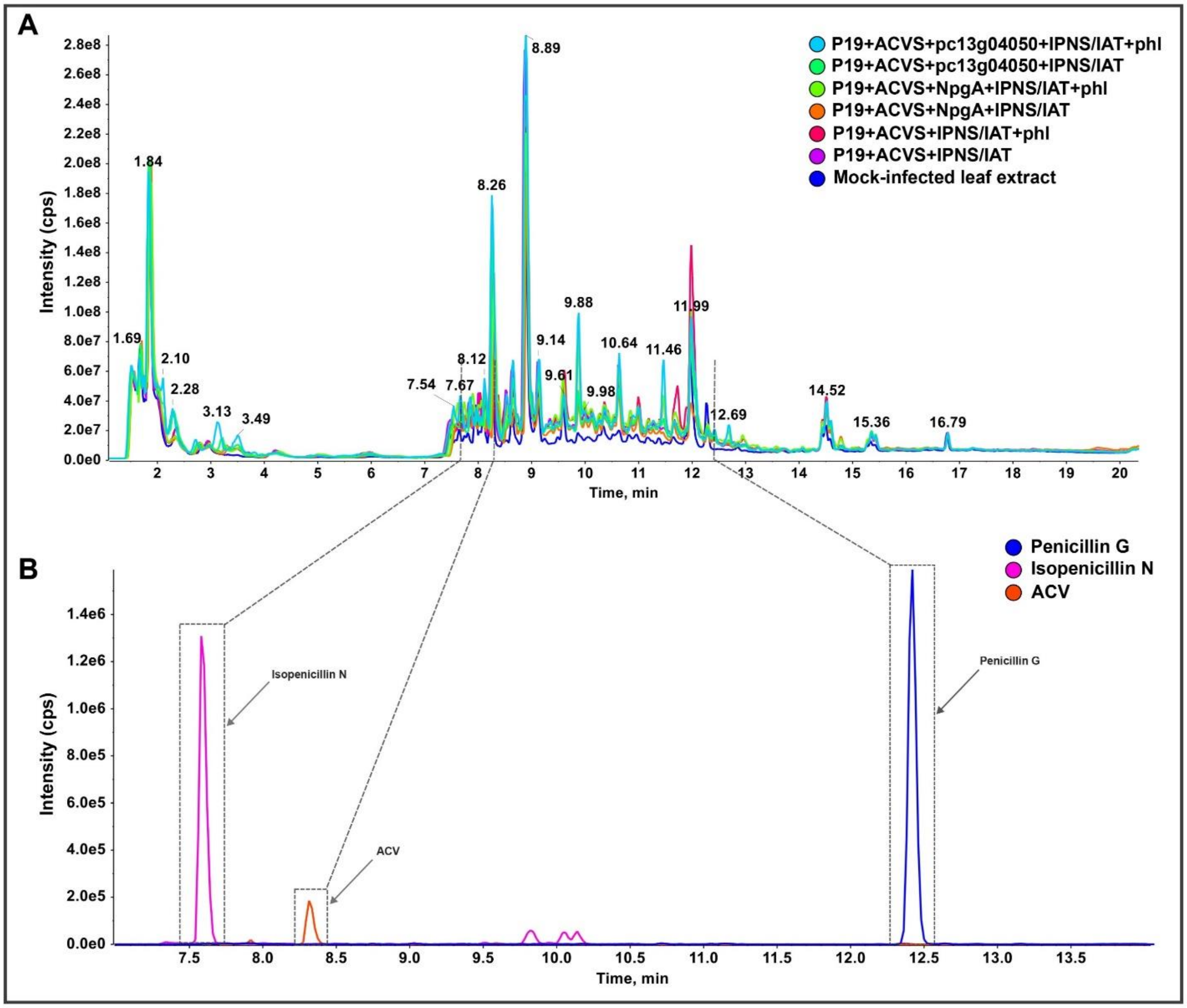

**LC-MS/MS analysis of penicillin G pathway reconstruction in *Nicotiana benthamiana*.** (A) Total ion chromatograms (TICs) showing overlaid metabolic profiles of samples from *N. benthamiana* leaf extracts transiently expressing different combinations of penicillin G biosynthetic pathway genes. Samples were analyzed by LC-MS/MS on a TripleTOF 6600+ system using reverse-phase separation under negative electrospray ionization (ESI) mode. (B) Extracted ion chromatograms (XICs) of pathway intermediates and the final product, detected as their $[M\text{-}H]^-$ ions. δ-(L-α-aminoadipyl)-L-cysteinyl-D-valine (ACV; *m/z* 362.138 ± 0.005, RT 8.4 min), isopenicillin N (*m/z* 358.107 ± 0.005, RT 7.6-7.7 min), and penicillin G (*m/z* 333.091 ± 0.005, RT 12.4 min) were selectively extracted to confirm their identities, retention times (RT), demonstrating successful *in-planta* reconstruction of the penicillin G biosynthetic pathway.

**Fig. S9.**

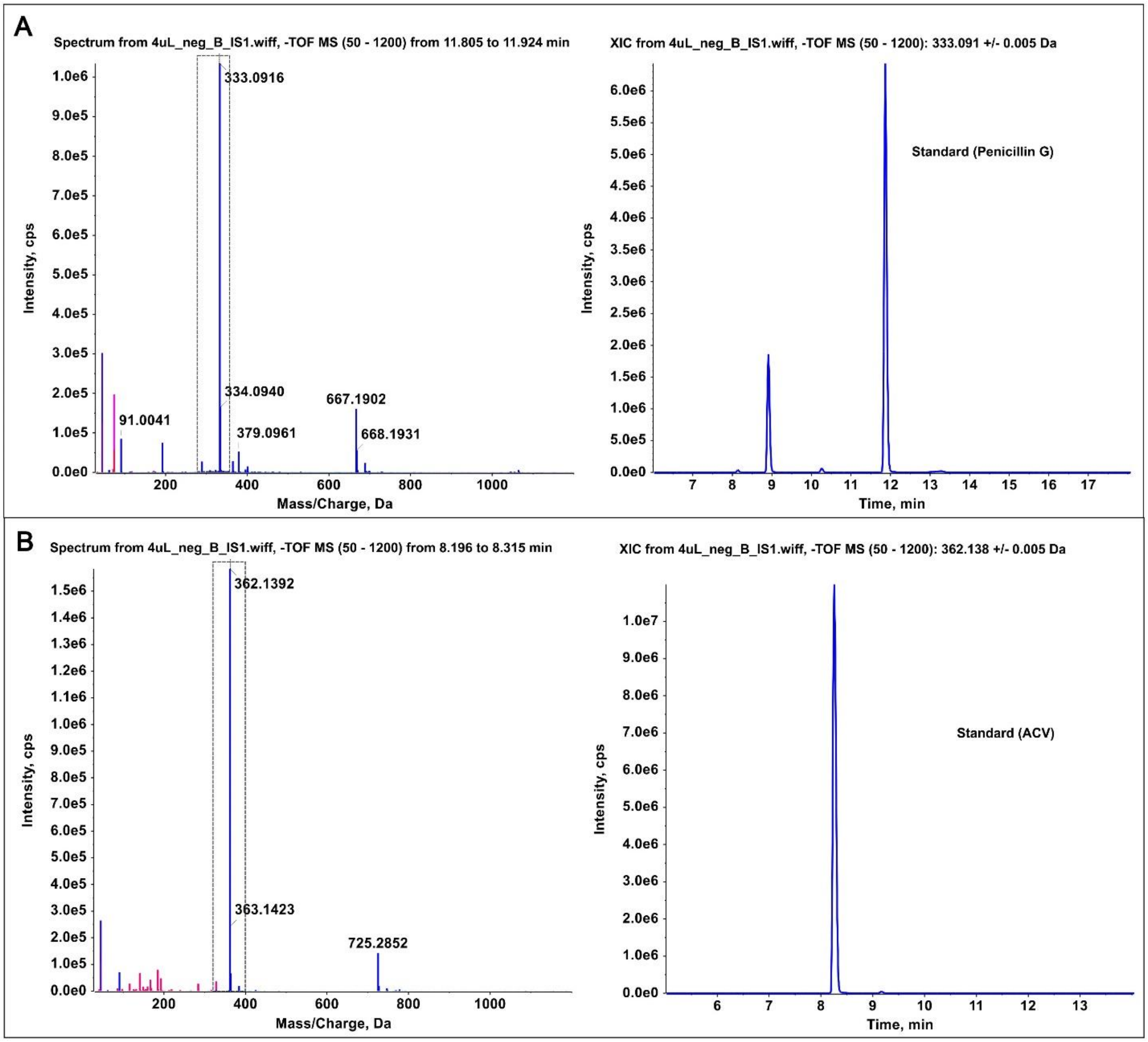

**Reverse-phase (RP) liquid chromatography tandem mass spectrometry (LC-MS/MS) analysis of reference standards in negative ESI mode.** Mass spectra (left) and extracted ion chromatograms (XIC; right) of (A) Penicillin G, showing the precursor ion $[M\text{-}H]^-$ at *m/z* 333.091 ± 0.005 at retention time (RT) 11.8 min, (B) ACV tripeptide, showing the precursor ion [M-H]- at *m/z* 362.1392 ± 0.005 at RT 8.4 min.

**Fig. S10.**

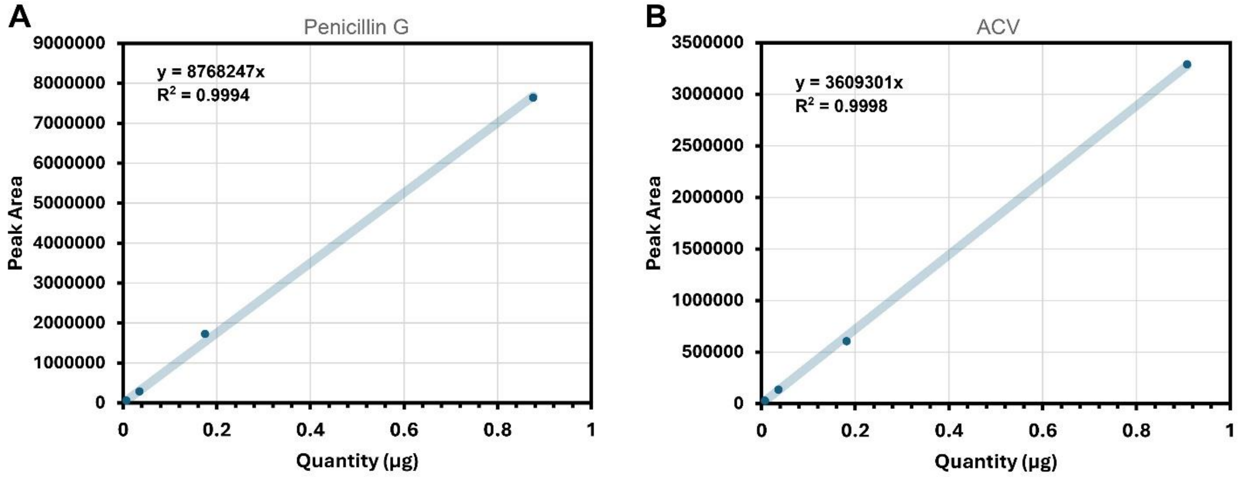

**Calibration curves for (A) penicillin G and (B) ACV**, constructed using four 5-fold quantities (0.00028, 0.0014, 0.007, and 0.035 µg). Peak areas were measured by LC-MS/MS on a TripleTOF 6600+ operated in negative electrospray ionization (ESI) mode. Linear regression yielded coefficients of determination ($R^2$) of 0.9994 for penicillin G and 0.9998 for ACV, demonstrating strong linearity across the calibration range. Note: These calibration curves were used for ACV and PenG quantification for experiments with PenG biosynthetic pathway enzymes; Fig 3, Fig. S4 & S6)

**Fig. S11.**

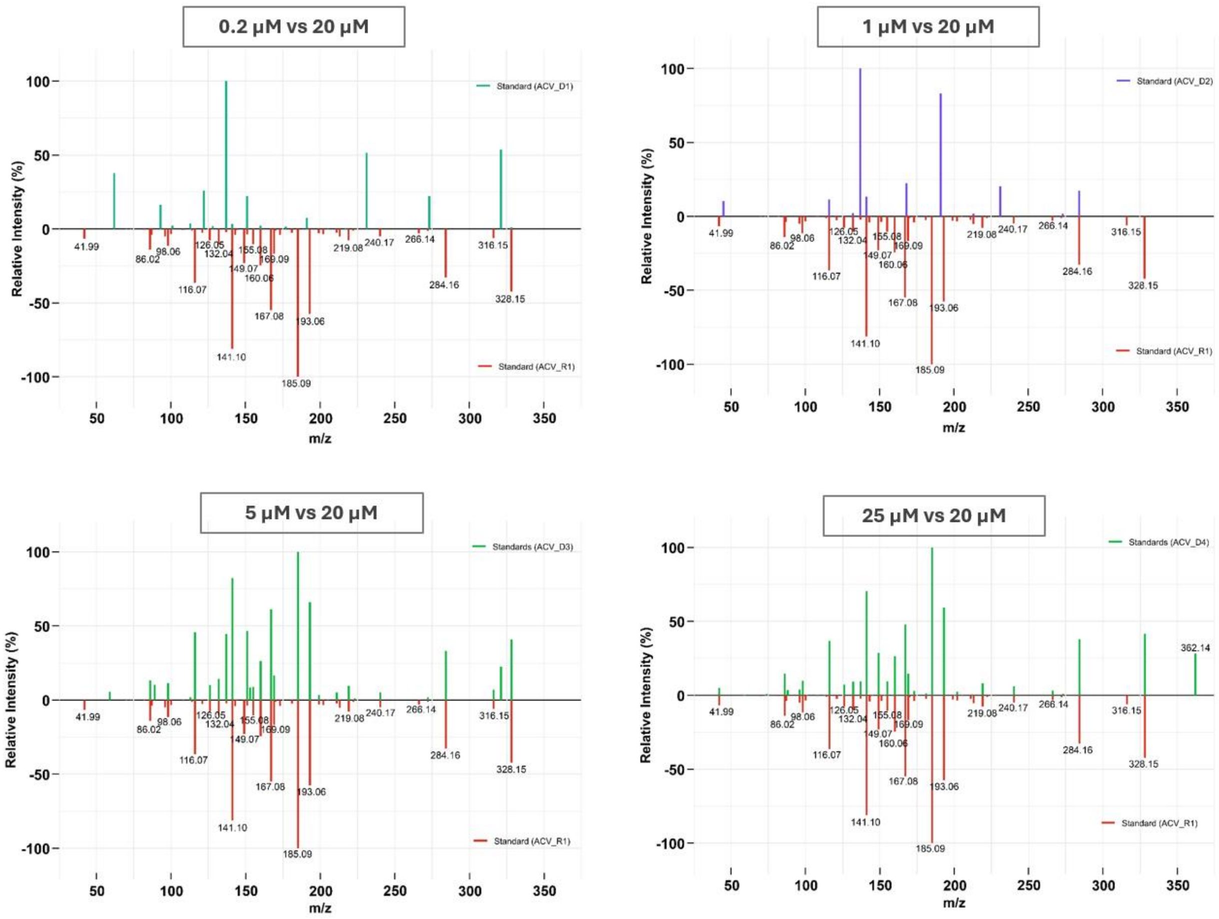

**Mirror plots comparing the MS/MS fragmentation spectra of ACV** standards at 0.2, 1, 5, and 25 µM (top) with the 20 µM reference ACV spectrum (bottom). The MS/MS spectra were acquired using LC-MS/MS on a TripleTOF 6600+ system operated in reverse-phase negative electrospray ionization (ESI) mode. Relative fragment ion intensities (%) are plotted as a function of *m/z* to assess reproducibility of ACV fragmentation across concentrations.

**Fig. S12.**

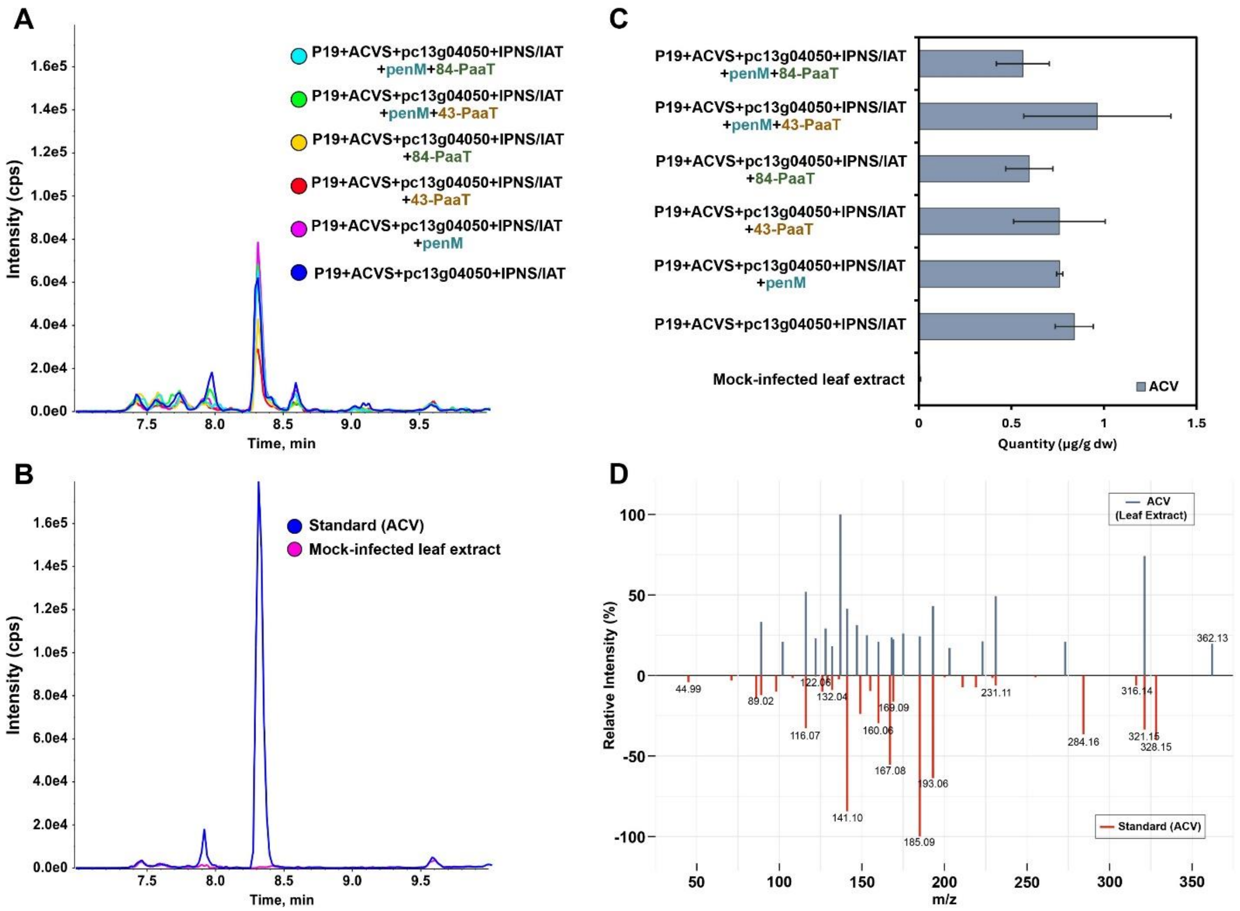

**Accumulation of ACV intermediates when PenG pathway genes and transporters are co-expressed.** LC-MS/MS-based detection and quantification of δ-(L-α-aminoadipyl)-L-cysteinyl-D-valine (ACV) in *N. benthamiana* leaf extracts expressing penicillin G biosynthetic pathway genes with and without transporters. (A) Extracted ion chromatograms (XICs) of ACV at retention time (RT) of 8.4 min, detected as $[M–H]^-$ at *m/z* 362.138 ± 0.005 from leaves transiently expressing pathway genes either alone or in combination with the PenM and PaaT transporters, (B) XICs from a mock-infiltrated leaf extract (negative control) and an external ACV standard, confirming RT alignment and mass accuracy, (C) Absolute ACV quantification (µg $g^{-1}$ dry weight) across the sample groups as shown in panel A, determined using an external calibration curve (mean ± SD, n = 3), and (D) MS/MS mirror plot comparing fragmentation patterns of ACV detected *in planta* to those of an external standard, showing matching fragment ions and confirming ACV identity. [Note:84-PaaT, C-terminal GFP tag: 43-PaaT, N-terminal GFP tag; SD: standard deviation].

**Fig. S13.**

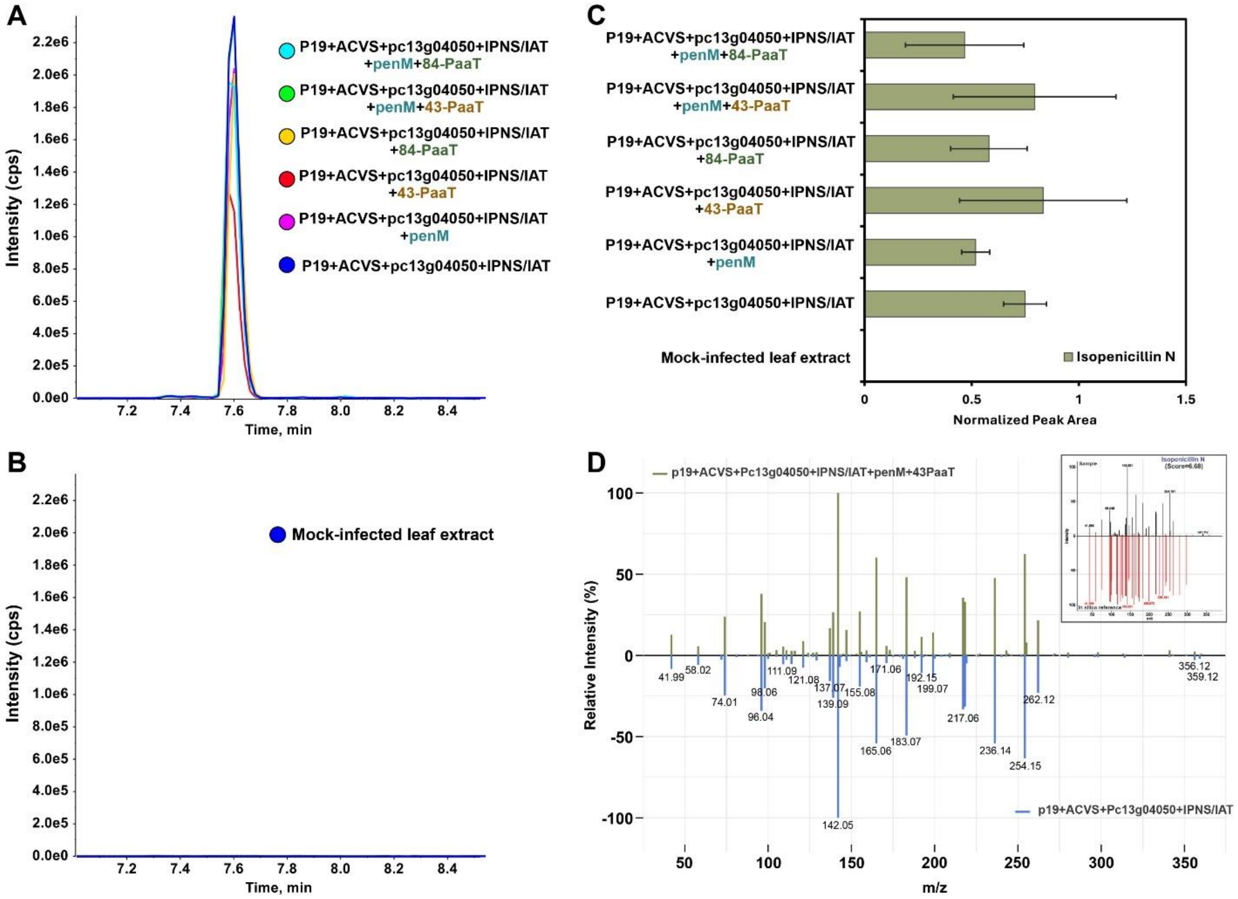

**Accumulation of IPN intermediates when PenG pathway genes and transporters are co-expressed.** LC-MS/MS based detection and characterization of isopenicillin N (IPN) in *N. benthamiana* leaf extracts expressing penicillin G biosynthetic genes. (A) Extracted ion chromatograms (XICs) of IPN at retention time (RT) 7.6 min, detected as $[M–H]^{-}$ at *m/z* 358.107 ± 0.005, from leaves transiently expressing pathway genes in combination with the PenM and PaaT transporters; (B) XICs from a mock-infiltrated leaf extract (negative control), (C) relative accumulation of IPN across the sample groups as shown in panel A, presented as normalized peak area (mean ± SD; n = 3), and (D) MS/MS mirror plot showing matching fragment ions for IPN detected *in planta,* in samples expressed with (top) and without (bottom) transporters; inset shows *in-silico* MS/MS fragmentation of IPN predicted using MS-Finder, matched against IPN spectra from databases (score=6.83/10), supporting structural assignment. [Note:84-PaaT, C-terminal GFP tag: 43-PaaT, N-terminal GFP tag; SD: standard deviation].

**Fig. S14.**

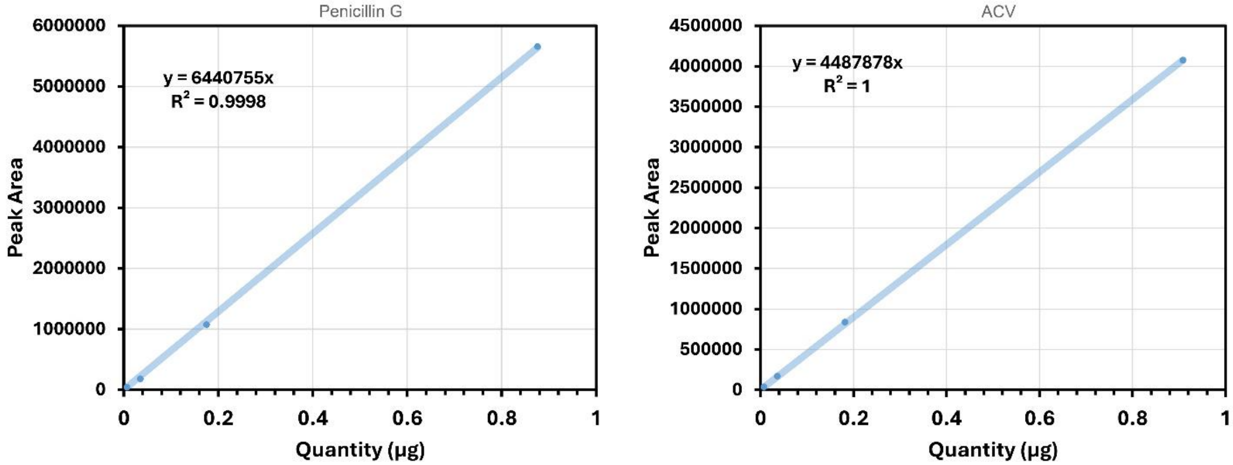

**Calibration curves for (A) penicillin G and (B) ACV, constructed using four 5-fold quantities (0.00028, 0.0014, 0.007, and 0.035 µg).** Peak areas were measured by LC-MS/MS on a TripleTOF 6600+ operated in negative electrospray ionization (ESI) mode. Linear regression yielded coefficients of determination ($R^2$) of 0.9998 for penicillin G and 1.0 for ACV, demonstrating strong linearity across the calibration range. Note: These calibration curves were used for ACV and PenG quantification without and with transporters, Fig 4, Fig S12)

**Fig. S15.**

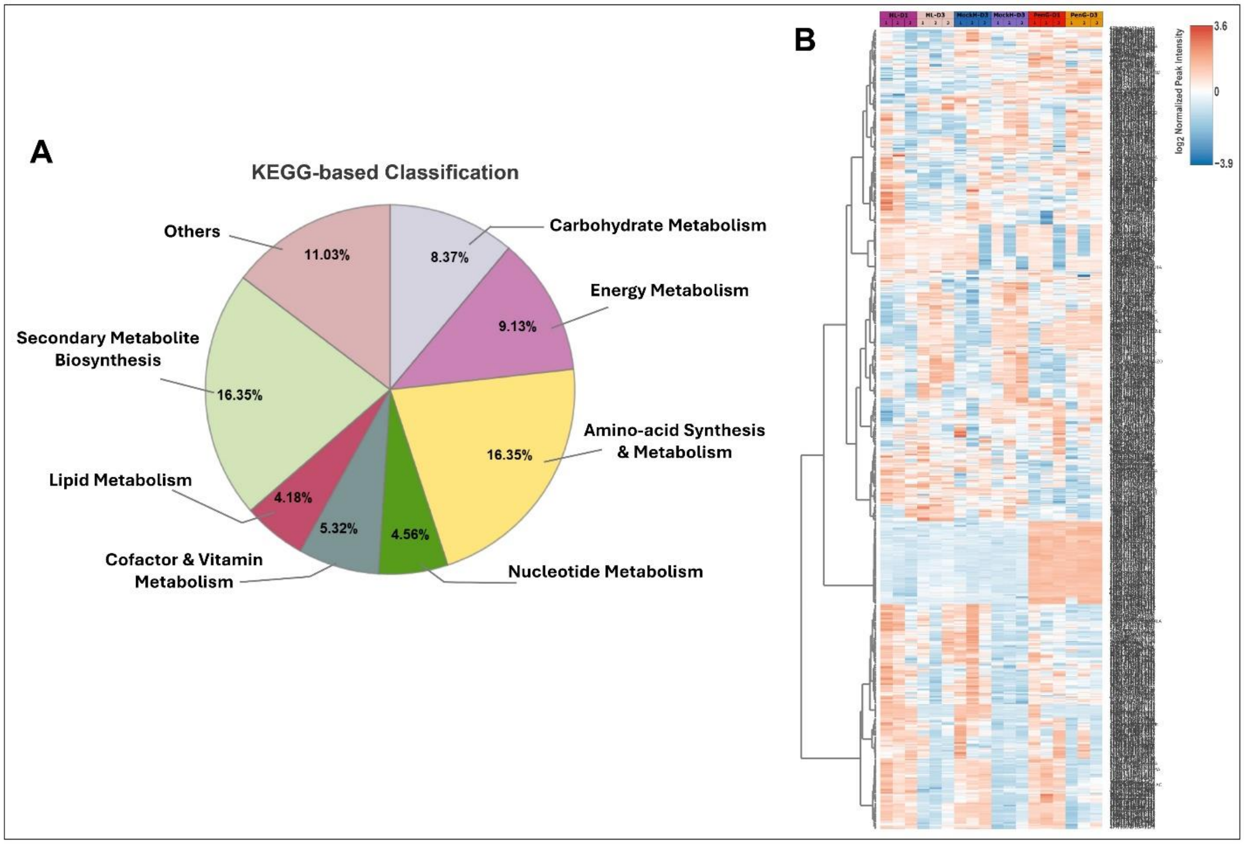

**Annotation and metabolic profiles of features identified using un-targeted metabolomics approach using reverse-phase (RP) and HILIC liquid chromatography tandem mass spectrometry (LC-MS/MS) analysis in *N. benthamiana* leaf samples infiltrated with penicillin G compound.** (A) Pie-chart showing KEGG-based classification of known metabolic features and (B) heatmap showing differential metabolic profiles of all identified features across penicillin infiltrated, mock and healthy leaves samples. [Abbreviations: HL = healthy leaf (no infiltration), MockH = infiltrated with water, PenG = infiltrated with penicillin G, and D1, D3 = Day 1 and 3 of post infiltration].

**Fig. S16.**

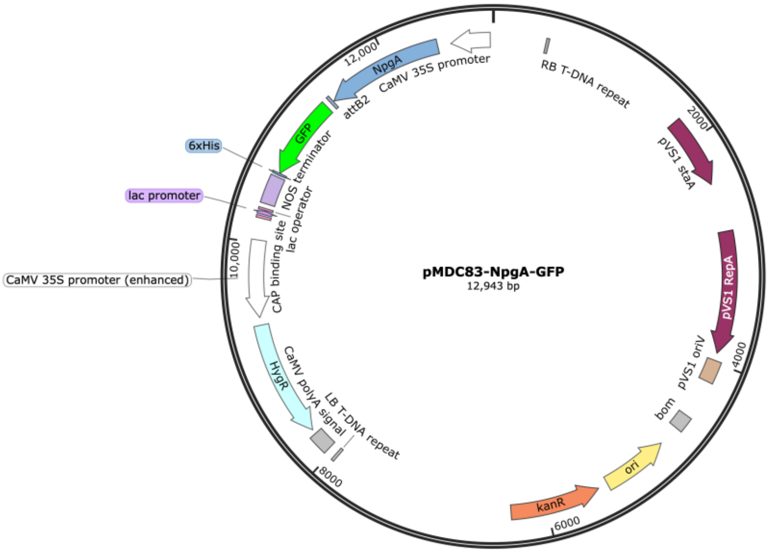

Plasmid map of pMDC83-NpgA-GFP used for confocal imaging and metabolite analysis.

**Fig. S17.**

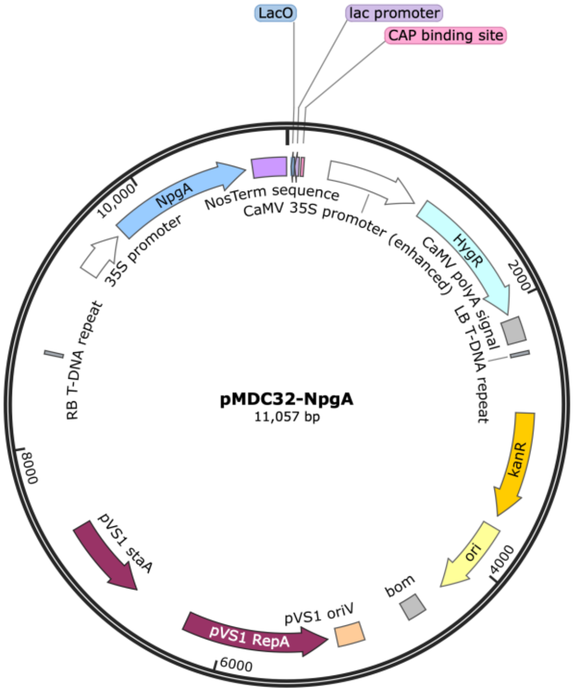

Plasmid map of pMDC32-NpgA used for metabolite analysis.

**Fig. S18.**

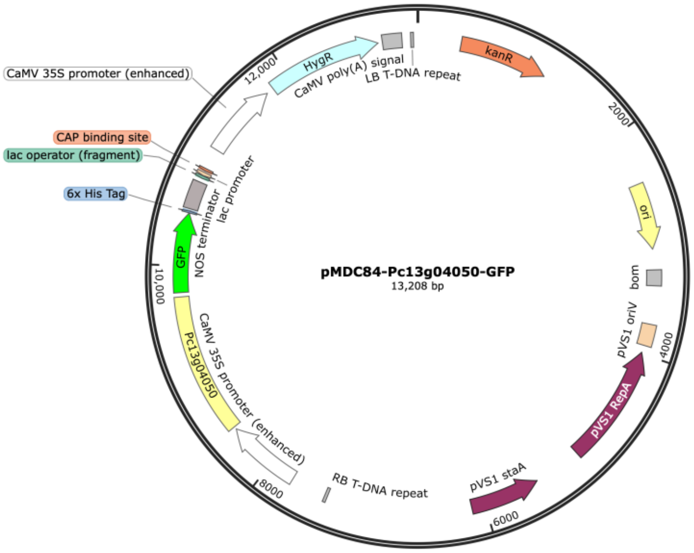

Plasmid map of pMDC84-Pc13g04050-GFP used for confocal imaging.

**Fig. S19.**

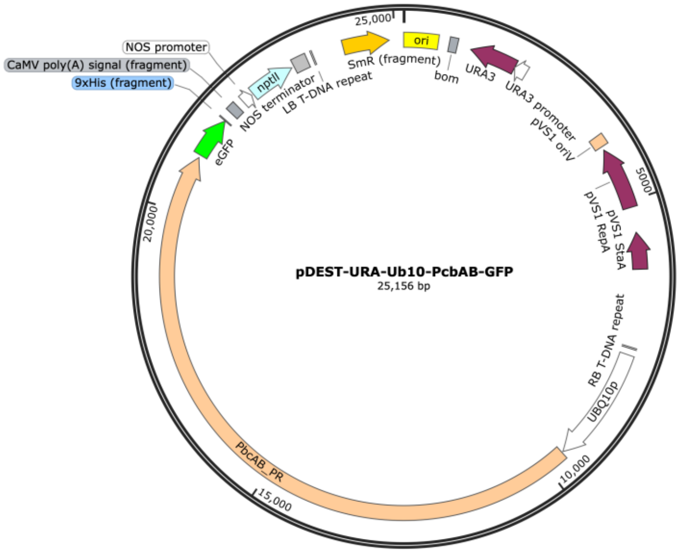

Plasmid map of pDEST-URA-Ub10-PcbAB-GFP used for confocal imaging and metabolite analysis. Here the native *pcbAB* sequence was used, cloned from *P. rubens*.

**Fig. S20.**

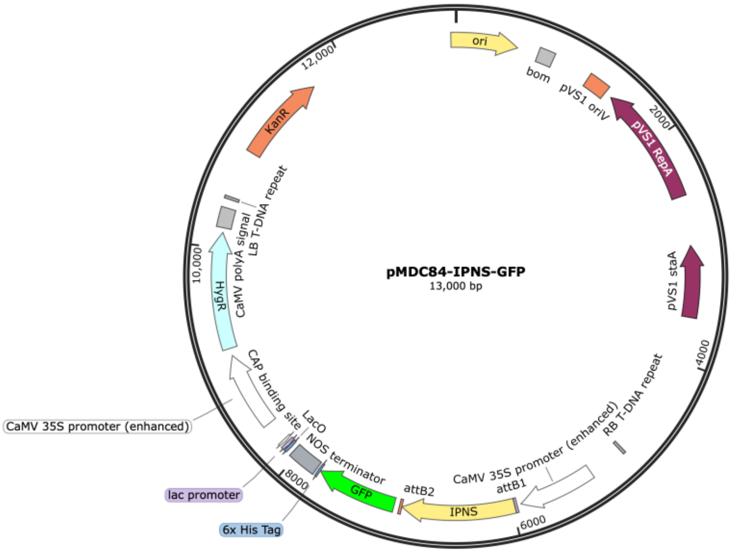

Plasmid map of pMDC84-IPNS-GFP used for confocal imaging.

**Fig. S21.**

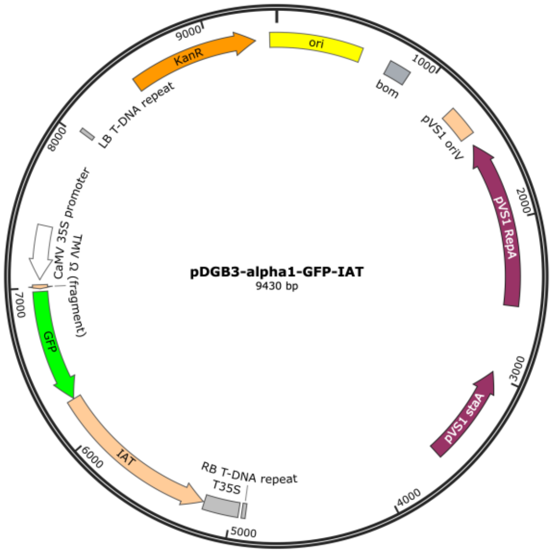

Plasmid map of pDGB3-alpha1-GFP-IAT used for confocal imaging.

**Fig. S22.**

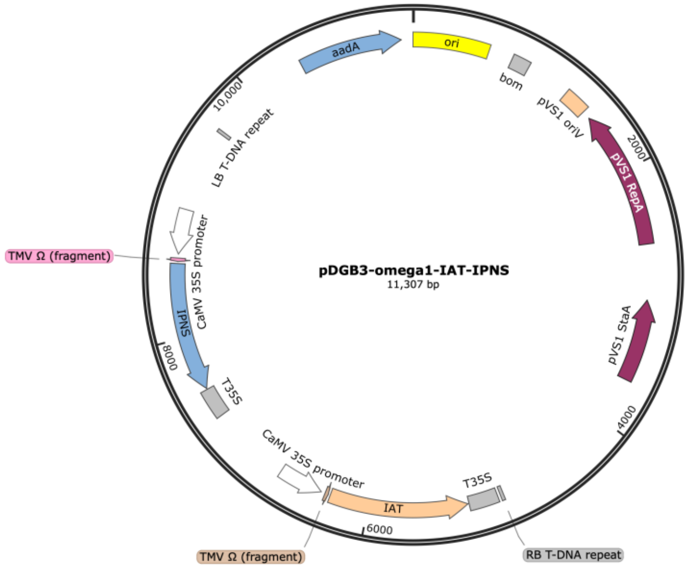

Plasmid map of pDGB3-omega1-IAT-IPNS with two expression cassettes on the T-DNA, where each cassette contains a CaMV35S promoter and T35S terminator, used for metabolite analysis.

**Fig. S23.**

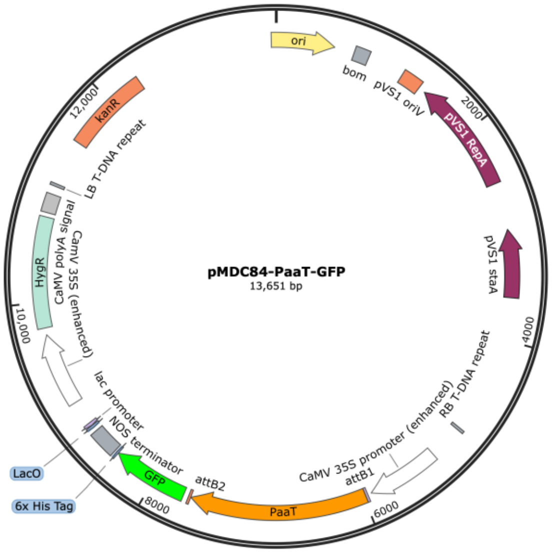

Plasmid map of pMDC84-PaaT- used for confocal imaging and metabolite analysis.

**Fig. S24.**

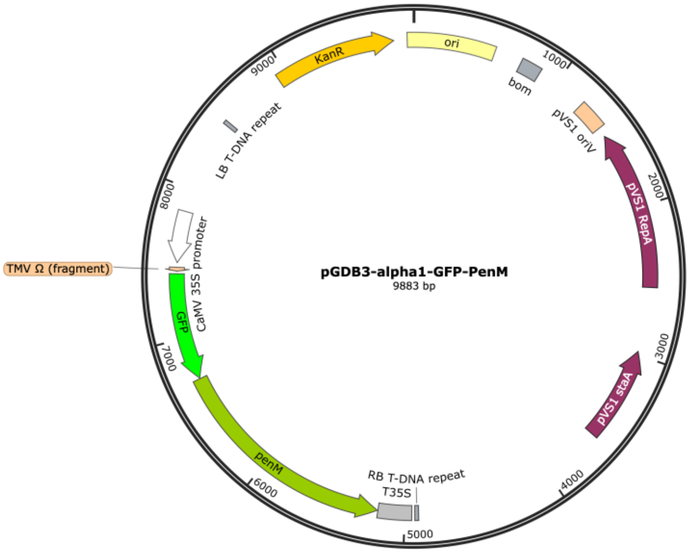

Plasmid map of pDGB3-alpha1-GFP-PenM used for confocal imaging and metabolite analysis.

**Fig. S25.**

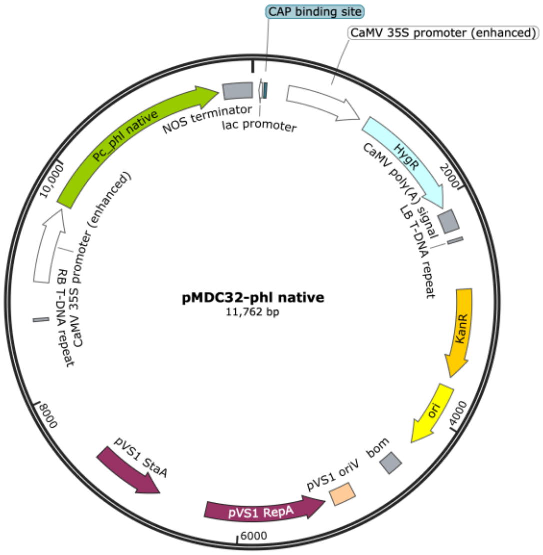

Plasmid map of pMDC32-phl (native sequence) used for metabolite analysis.

**Fig. S26.**

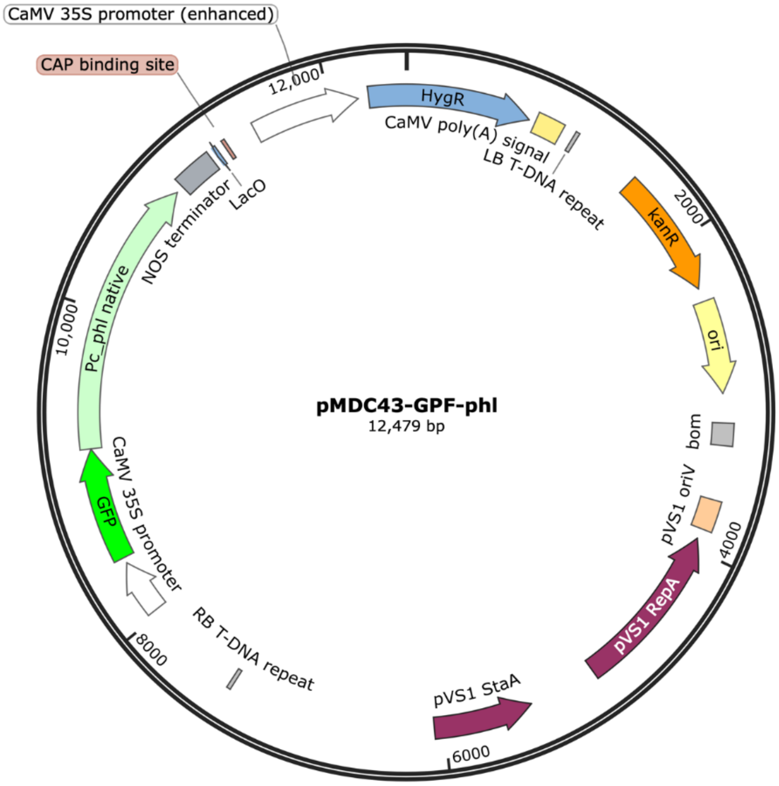

Plasmid map of pMDC43-GFP-phl used for confocal imaging

**Fig. S27.**

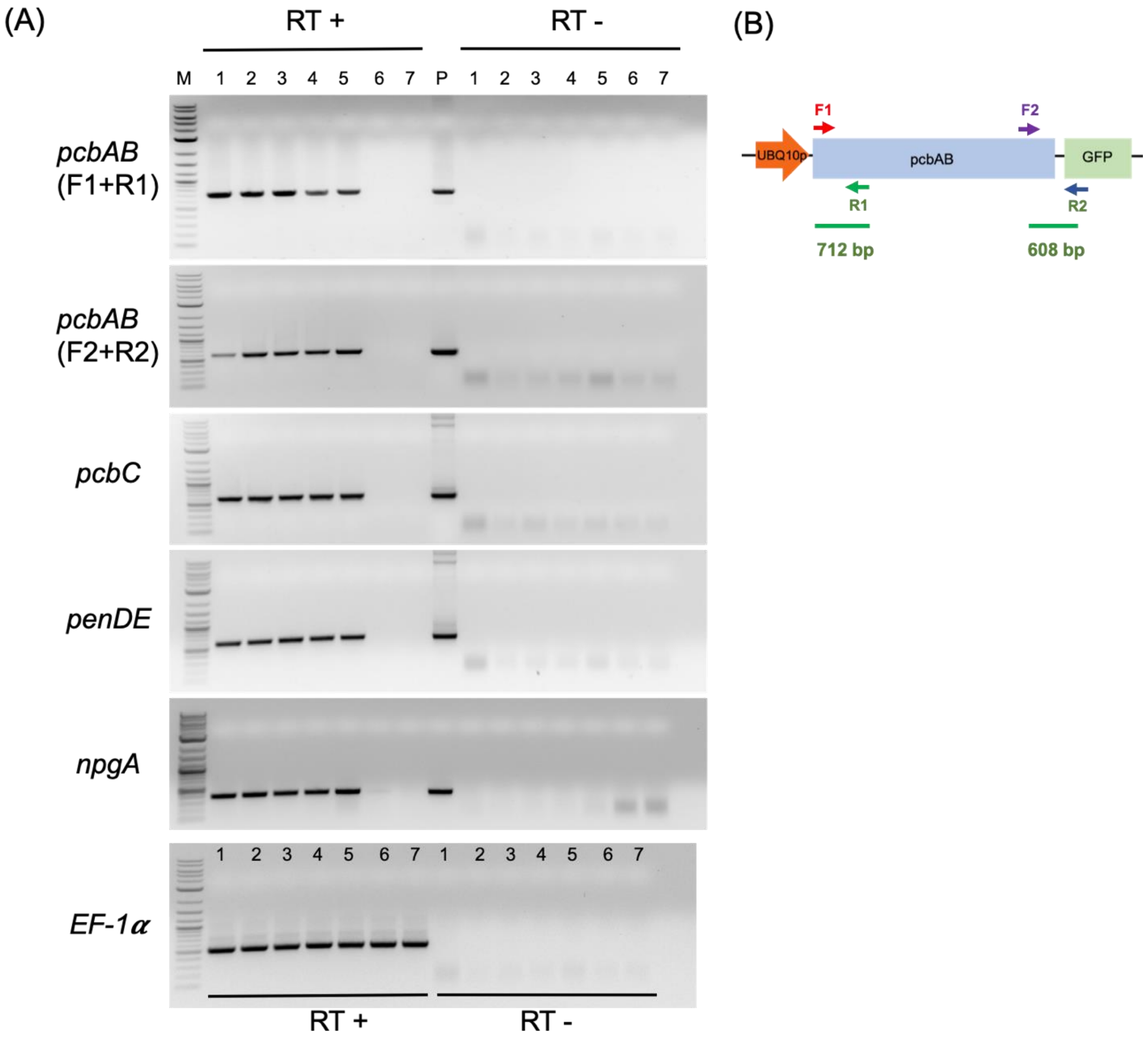

Confirmation of the expression of Penicillin G biosynthetic genes in infiltrated *N. benthamiana* leaves. (A) RT-PCR analysis to confirm the expression of *pcbAB*, *pcbC*, *penDE,* and *npgA* genes. *EF-1α* served as the reference gene showing the consistency of RNA isolation and cDNA synthesis across all samples. Lane 1-5: RNA samples were isolated from leaves co-infiltrated with *pcbAB*, *pcbC*, *penDE*, and *npgA* genes along with the gene silencing suppressor p19; five samples represent five individual infiltrated leaves. Lane 6 and 7: RNA samples were isolated from two individual infiltrated leaves with the p19 alone. RT+: Synthesized cDNAs were used as templates for PCR. RT-: DNA-free RNAs were used as templates for PCR. P: plasmids as the positive controls. M: 1 kb DNA ladder. (B) Schematic representation of primer design to target pcbAB transcripts. F1 and F2: forward primers; R1 and R2: reverse primers.

**Fig. S28.**

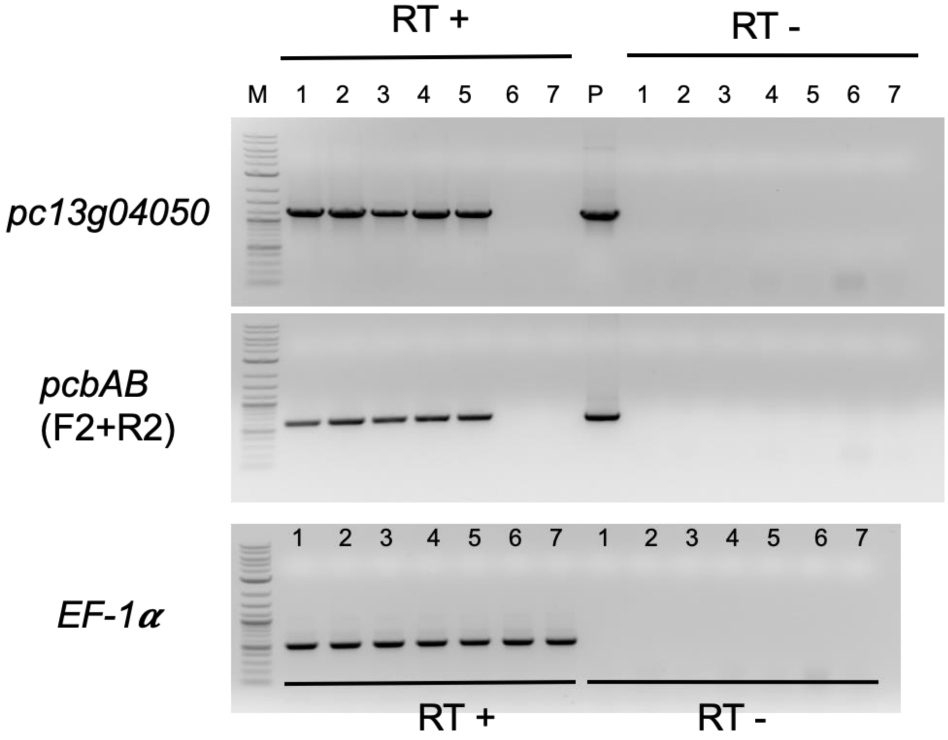

Confirmation of the expression of *pc13g04050* and *pcbAB* genes in infiltrated N.*benthamiana* leaves. (A) RT-PCR analysis to confirm the expression of *pcbAB* and *pc13g04050* genes. *EF-1α* served as the reference gene showing the consistency of RNA isolation and cDNA synthesis across all samples. Lane 1-5: RNA samples were isolated from leaves co-infiltrated with *pcbAB*, pc13g04050, and *mCherry* genes along with gene silencing suppressor p19; five samples represent five individual infiltrated leaves. Lane 6 and 7: RNA samples were isolated from two individual infiltrated leaves with the p19 gene alone. RT+: Synthesized cDNAs were used as templates for PCR. RT-: DNA-free RNAs were used as templates for PCR. P: plasmids as the positive controls. M: 1 kb DNA ladder.

**Fig. S29.**

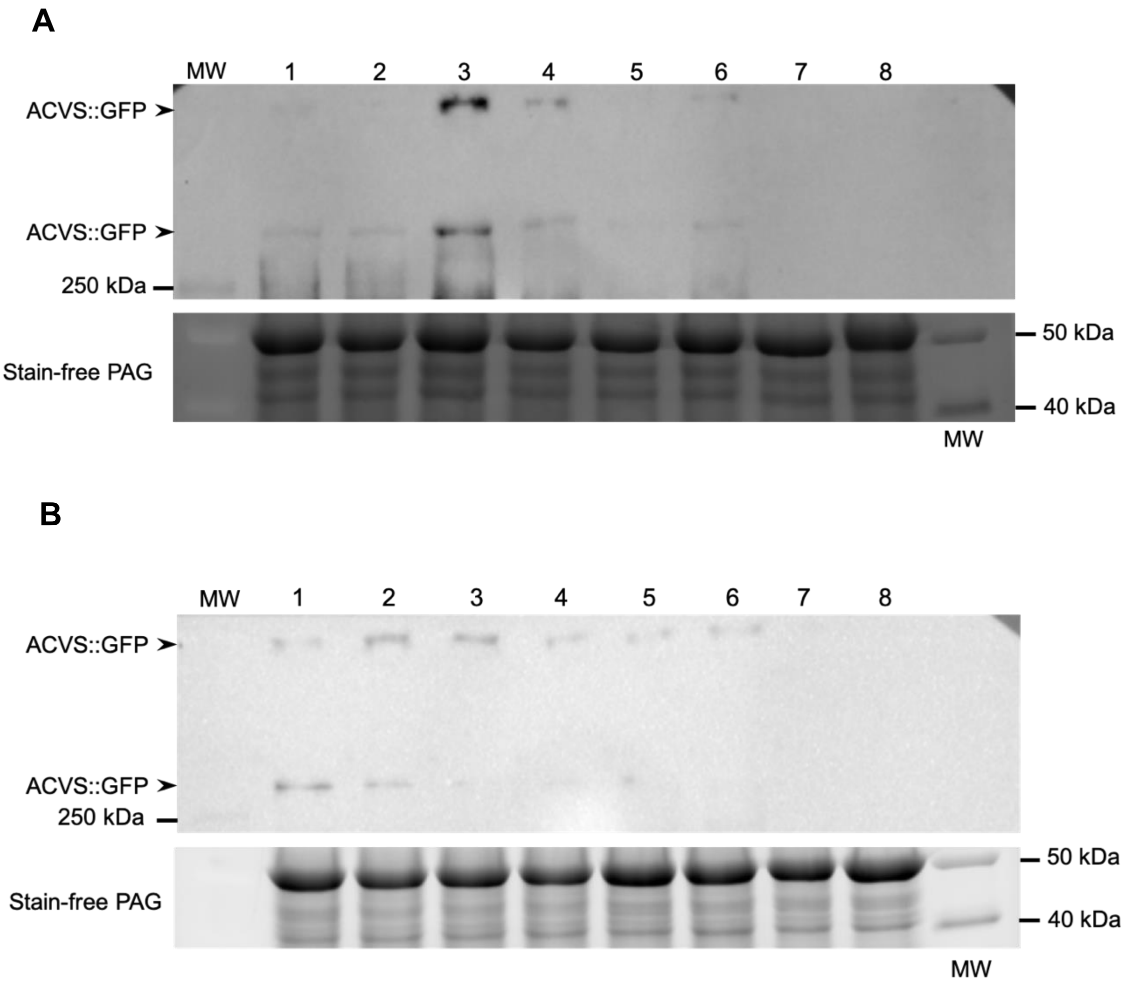

Western blot analysis for the identification of ACVS::GFP fusion proteins expressed in *N. benthamiana* leaves. Lanes1-6 in (A) and (B) show the presence of 453 kDa ACVS::GFP cross-reacting bands, which are above the 250 kDa band of the protein standards. (A) Lanes 1-5 are five independent protein samples extracted from leaves co-infiltrated with *pcbAB-gfp*, *pc13g04050*, and gene silencing suppressor p19. Lane 6 is the protein sample extracted from leaves co-infiltrated with *pcbAB-gfp* and gene silencing suppressor p19. Lane 7 is the p19 only treatment, and Lane 8 is a mock treatment. (B) Lanes 1-5 are five independent protein samples extracted from leaves co-infiltrated with *pcbAB-gfp*, *npgA*, and gene silencing suppressor p19. Lane 6 is the protein sample extracted from leaves co-infiltrated with *pcbAB-gfp* and gene silencing suppressor p19. Lane 7 is the p19 only treatment, and Lane8 is a mock treatment. Stain-free PAG, the stain-free polyacrylamide gel to show the protein amount in each lane.

**Fig. S30.**

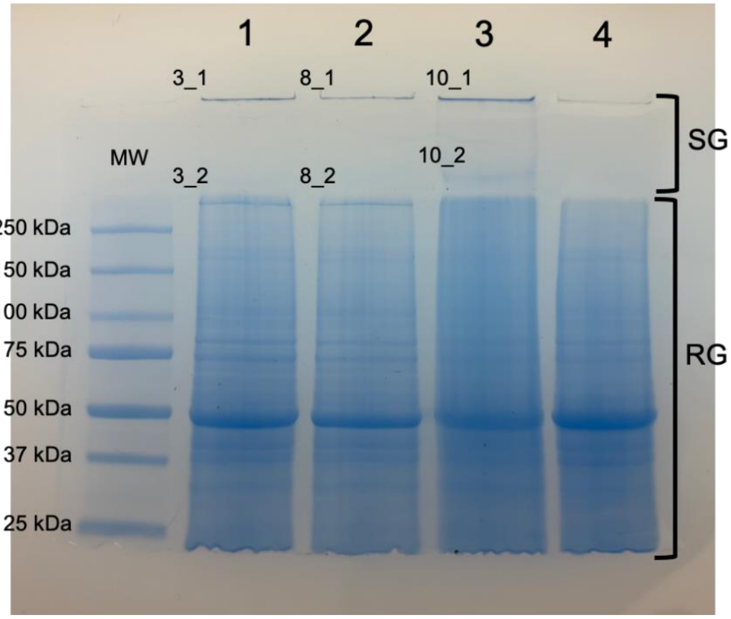

Coomassie Blue stained 10% SDS-PAGE gel showing proteins separated by size via gel electrophoresis. Lane 1 and 2: two independent protein samples were extracted from leaves co-infiltrated with *pcbAB-gfp*, *pc13g04050*, and gene silencing suppressor p19. Lane 3: the protein sample was extracted from leaves co-infiltrated with *pcbAB-gfp*, *npgA*, and p19. Lane 4: p19 alone treatment. Band 3-1, 3-2, 8-1, 8-2, 10-1, and 10-2: gel slices were sent for peptide sequencing. 3-1, 8-1, and 10-1: gel slices were located at the bottom of the stacking gel (SG) wells. 3-2 and 8-2 gel slices were located at the top of the resolving gel (RG) and near the SG-RG border; 10-2 was at the bottom of SG.

**Fig. S31.**

| | | | | |
|---|---|---|---|---|
| MTQLKPPNGT | TPIGFSATTS | LNASGSSSVK | NGTIKPSNGI | FKPSTRDTMD |
| PCSGNAADGS | IRVRFRGGIE | RWKECVNQVP | ER**CDLSGLTT** | **DSTRYQLAST** |
| **GFGDASAAYQ** | **ER**LMTVPVDV | HAALQELCLE | RRVSVGSVIN | FSVHQMLKGF |
| GNGTHTITAS | LHR**EQNLQNS** | **SPSWVVSPTI** | **VTHENR**DGWS | VAQAVESIEA |
| GR**GSEKESVT** | **AIDSGSSLVK** | MGLFDLLVSF | VDADDAR**IPC** | **FDFPLAVIVR** |
| ECDANLSLTL | RFSDCLFNEE | TICNFTDALN | ILLAEAVIGR | **VTPVADIELL** |
| **SAEQK**QQLEE | WNNTDGEYPS | SKRLHHLIEE | VVERHEDKIA | VVCDER**ELTY** |
| **GELNAQGNSL** | **AR**YLR**SIGIL** | **PEQLVALFLD** | **KSEK**LIVTIL | GVWKSGAAYV |
| PIDPTYPDER | VRFVLDDTKA | R**AIIASNQHV** | **ER**LQREVIGD | RNLCIIR**LEP** |
| **LLASLAQDSS** | **KFPAHNLDDL** | **PLTSQQLAYV** | **TYTSGTTGFP** | **K**GIFK**QHTNV** |
| **VNSITDLSAR** | YGVAGQHHEA | ILLFSACVFE | PFVRQTLMAL | VNGHLLAVIN |
| DVEK**YDADTL** | **LPFIR**RHSIT | YLNGTASVLQ | EYDFSDCPSL | NR**IILVGENL** |
| **TEAR**YLALRQ | RFKNR**ILNEY** | **GFTESAFVTA** | **LKIFDPESTR** | KDTSLGRPVR |
| NVKCYILNPS | LKR**VPIGATG** | **ELHIGGLGIS** | **K**GYLNRPELT | PHR**FIPNPFQ** |
| **TDCEKQLGIN** | **SLMYK**TGDLA | RWLPNGEVEY | LGRADFQIKL | RGIR**IEPGEI** |
| **ETMLAMYPR**V | R**TSLVVSK**KL | RNGPEETTNE | HLVGYYVCDS | ASVSEADLLS |
| FLEKKLPRYM | IPTRLVQLSQ | IPVNVNGKAD | LR**ALPAVDIS** | **NSTEVR**SDLR |
| GDTEIALGEI | WADVLGARQR | SVSRNDNFFR | LGGHSITCIQ | LIARIRQRLS |
| VSISVEDVFA | TRTLER**MADL** | **LQNK**QQEKCD | KPHEAPTELL | EENAATDNIY |
| LANSLQQGFV | YHYLK**SMEQS** | **DAYVMQSVLR** | YNTTLSPDLF | QRAWK**HAQQS** |
| **FPALR**LRFSW | EK**EVFQLLDQ** | **DPPLDWRFLY** | **FTDVAAGAVE** | **DRKLEDLRR**Q |
| DLTERFKLDV | GRLFRVYLIK | HSENRFTCLF | SCHHAILDGW | SLPLLFEKVH |
| ETYLQLLHGD | NLTSSMDDPY | TRTQRYLHAH | REDHLDFWAG | VVQKINER**CD** |
| **MNALLNER**SR | **YKVQLADYDQ** | **VQEQR**QLTIA | LSGDAWLADL | RQTCSAQGIT |
| LHSILQFVWH | AVLHAYGGGT | HTITGTTISG | R**NLPILGIER** | **AVGPYINTLP** |
| **LVLDHSTFKD** | **KTIMEAIEDV** | **QAK**VNVMNSR | GNVELGRLHK | TDLK**HGLFDS** |
| **LFVLENYPNL** | **DK**SR**TLEHQT** | **ELGYSIEGGT** | **EKLNYPLAVI** | **AR**EVETTGGF |
| TVSICYASEL | FEEVMISELL | HMVQDTLMQV | ARGLNEPVGS | LEYLSSIQLE |
| QLAAWNATEA | EFPDTTLHEM | FENEASQKPD | K**IAVVYEETS** | **LTYR**ELNERA |
| NRMAHQLR**SD** | **VSPNPNEVIA** | **LVMDK**SEHMI | VNILAVWK**SG** | **GAYVPIDPGY** |
| **PNDR**IQYILE | DTQALAVIAD | SCYLPR**IKGM** | **AASGTLLYPS** | **VLPANPDSKW** |
| **SVSNPSPLSR** | **STDLAYIIYT** | **SGTTGRPK**GV | TVEHHGVVNL | QVSLSKVFGL |
| RDTDDEVILS | FSNYVFDHFV | EQMTDAILNG | QTLLVLNDGM | RGDKERLYRY |
| IEKNR**VTYLS** | **GTPSVVSMYE** | **FSR**FKDHLR**R** | **VDCVGEAFSE** | **PVFDK**IRETF |
| HGLVINGYGP | TEVSITTHKR | **LYPFPER**RMD | KSIGQQVHNS | TSYVLNEDMK |
| R**TPIGAVGEL** | **YLGGEGVVR**G | YHNRADVTAE | R**FIPNPFQSE** | **EDKR**EGRNSR |
| LYKTGDLVR**W** | **IPGSSGEVEY** | **LGR**NDFQVKI | RGLR**IELGEI** | **EAILSSYHGI** |
| **K**QSVVIAKDC | REGAQK**FLVG** | **YYVADAALPS** | **AAIR**RFMQSR | LPGYMVPSRL |
| ILVSK**FPVTP** | **SGK**LDTK**ALP** | **PAEEESEIDV** | **VPPR**SEIERS | LCDIWAELLE |
| MHPEEIGIYS | DFFSLGGDSL | KSTKLSFMIH | ESFNRAVSVS | ALFCHRTVEA |
| QTHLILNDAA | DVHEITPIDC | NDTQMIPVSR | AQERLLFIHE | FENGSNAYNI |
| DAAFELPGSV | DASLLEQALR | GNLARHEALR | TLLVK**DHATG** | **IYLQKVLSPD** |
| **EAQGMFSVNV** | **DTAK**QVER**LD** | **QEIASLSQHV** | **FR**LDDELPWE | ARILKLESGG |
| LYLILAFHHT | CFDAWSLK**VF** | **EQELRALYAA** | **LQK**TK**SAANL** | **PALK**AQYKEY |
| ALYHRRQLSG | DRMRNLSDFW | LR**KLIGLEPL** | **QLITDRPR**PV | QFK**YDGDDLS** |
| **IELSKK**ETEN | LRGVAKRCKS | SLYVVLVSVY | CVMLASYANQ | SDVSVGIPVS |
| HRTHPQFQSV | IGFFVNLVVL | RVDISQSAIC | GLIRRVMK**EL** | **VDAQLHQDMP** |
| **FQEVTKLLQV** | **DNDPSRHPLV** | **QNVFNFESR**A | NGEHDARSED | EGSLAFNQYR |
| PVQPVDSVAK | **FDLNATVTEL** | **ESGLRVNFNY** | **ATSLFNKSTI** | **QGFLHTYEYL** |
| **LRQLSELSAE** | **GINEDTQLSL** | **VRPTENGDLH** | **LPLAQSPLAT** | **TAEEQKVASL** |
| **NQAFER**EAFL | AAEK**IAVVQG** | **DRALSYADLN** | **GQANQLAR**YI | QSVSCIGADD |
| GIALMLEKSI | DTIICILAIW | K**AGAAYVPLD** | **PTYPPGRVQL** | **ILEEIKAKAV** |
| **LVHSSHASK**C | ERHGAKVIAV | DSPAIETAVS | QQSAADLPTI | ASLGNLAYII |
| FTSGTSGKPK | **GVLVEQK**AVL | LLRDALRERY | FGRDCTKHHG | VLFLSNYVFD |
| FSVEQLVLSV | LSGHK**LIVPP** | **AEFVADDEFY** | **RMASTHGLSY** | **LSGTPSLLQK** |
| IDLAR**LDHLQ** | **VVTAAGEELH** | **ATQYEK**MRRR | FNGPIYNAYG | VTETTVYNII |
| AEFTTNSIFE | NALR**EVLPGT** | **RAYVLNAALQ** | **PVPFDAVGEL** | **YLAGDSVTRG** |
| **YLNQPLLTDQ** | **R**FIPNPFCK**E** | **EDIAMGR**FAR | LYKTGDLVRS | RFNR**QQQPQL** |
| **EYLGR**GDLQI | KMRGYR**IEIS** | **EVQNVLTSSP** | **GVR**EGAVVAK | **YENNDTYSRT** |
| **AHSLVGYYTT** | **DNETVSEADI** | **LTFMK**ARLPT | YMVPSHLCCL | EGALPVTING |
| KLDVR**RLPEI** | **INDSAQSSYS** | **PPR**NIIEAKM | CRLWESALGM | ERCGIDDDLF |
| KLGGDSITSL | HLVAQIHNQV | GCKITVRDIF | EHRTARALHD | HVFMKDSDR**S** |
| **NVTQFR**TEQG | PVIGEAPLLP | IQDWFLSKAL | QHPMYWNHTF | YVR**TPELDVD** |
| **SLSAAVRDLQ** | **QYHDVFR**MRL | KR**EEVGFVQS** | **FAEDFSPAQL** | **R**VLNVKDVDG |
| SAAVNEILDG | WQSGFNLENG | PIGSIGYLHG | YEDRSARVWF | SVHHMAIDTV |
| SWQILVR**DLQ** | **TLYR**NGSLGS | KGSSFRQWAE | AIQNYKASDS | ERNHWNK**LVM** |
| **ETASSISALP** | **TSTGSR**VRLS | RSLSPEK**TAS** | **LIQGGIDR**QD | VSVYDSLLTS |
| VGLALQHIAP | TGPSMVTIEG | HGR**EEVDQTL** | **DVSRTMGWFT** | **TMYPFEIPRL** |
| **STENIVQGVV** | **AVSER**FRQVP | AR**GVGYGTLY** | **GYTQHPLPQV** | **TVNYLGQLAR** |
| KQSKPKEWVL | AVGDNEFEYG | LMTSPEDKDR | SSSAVDVTAV | CIDGTMIIDV |
| DSAWSLEESE | QFISSIEEGL | NKILDGRASQ | QTSR**FPDVPQ** | **PAETYTPYFE** |
| **YLEPPR**QGPT | LFLLPPGEGG | AESYFNNIVK | RLRQTNMVVF | NNYYLHSKR**L** |
| **RTFEELAEMY** | **LDQVR**GIQPH | GPYHFIGWSF | GGILAMEMSR | RLVASDEKIG |
| FLGIIDTYFN | VRGATRTIGL | GDTEILDPIH | HIYNPDPANF | QR**LPSATDR**I |
| VLFK**AMRPNN** | **KYESENQR**R**L** | **YEYYDGTR**LN | GLDSLLPSDS | DVQLVPLTDD |
| THFSWVGNPQ | QVEQMCATIK | EHLARYLIAW | RASR**GGPVPV** | **EK**MSKGEELF |
| TGVVPILVEL | DGDVNGHK**FS** | **VSGEGEGDAT** | **YGK**LTLKFIC | TTGKLPVPWP |
| TLVTTLTYGV | QCFSRYPDHM | KRHDFFK**SAM** | **PEGYVQER**TI | FFKDDGNYKT |
| R**AEVKFEGDT** | **LVNR**IELKGI | DFKEDGNILG | HKLEYNYNSH | NVYIMADKQK |
| NGIKANFKTR | HNIEDGGVQL | ADHYQQNTPI | GDGPVLLPDN | HYLSTQSALS |
| KDPNEKRDHM | VLLEFVTAAG | ITHGMVELYK | HHHHHH | |

Amino acid coverage showing parts of the ACVS reference protein sequence (3-1) are identified by detected peptides from MS/MS. Segments with identified peptides are marked in yellow, and modified amino acids are in green. The sequence coverage percentages of 3-1 is 38%.

**Fig. S32.**

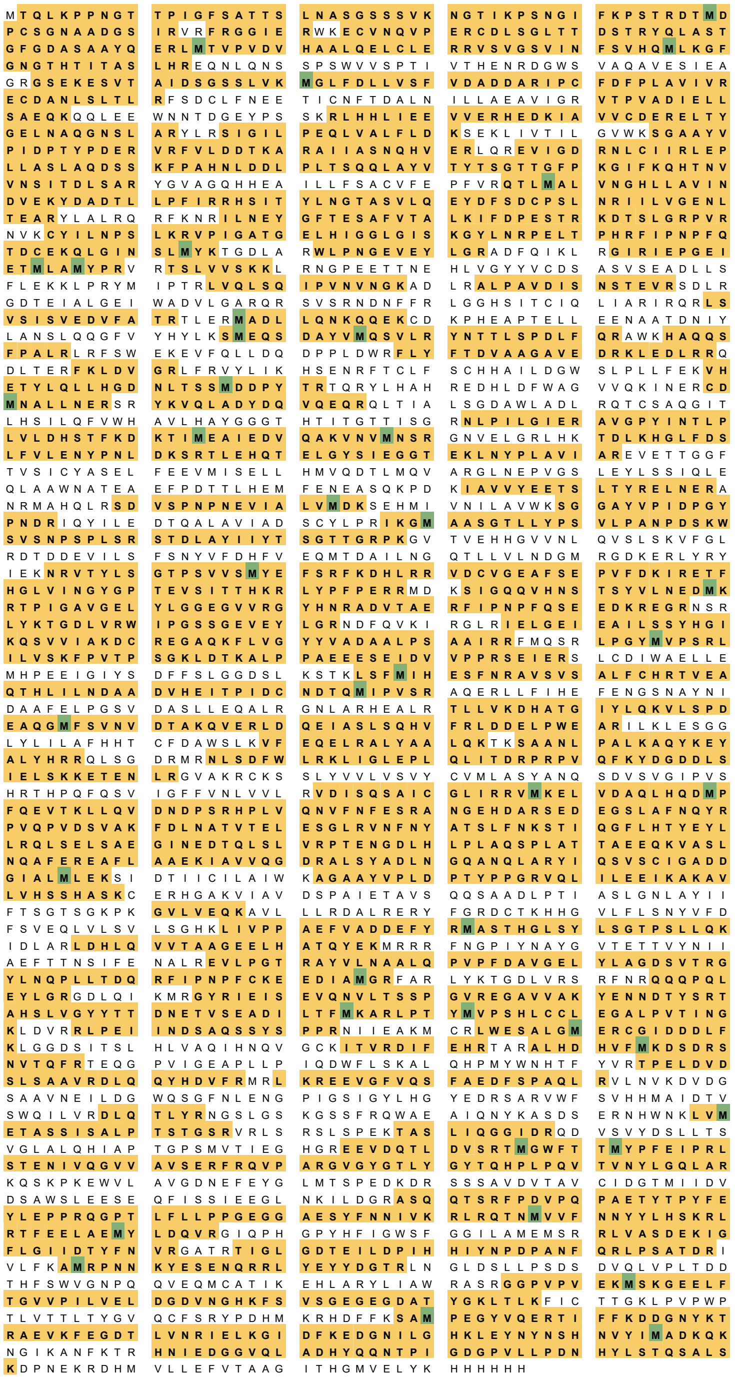

Amino acid coverage showing parts of the ACVS reference protein sequence (3-2) are identified by detected peptides from MS/MS. Segments with identified peptides are marked in yellow, and modified amino acids are in green. The sequence coverage percentages of 3-2 is 62%.

**Fig. S33.**

```
MTQLKPPNGT TPIGFSATTS LNASGSSSVK NGTIKPSNGI FKPSTRDTMD
PCSGNAADGS IRVRFRGGIE RWKECVNQVP ERCDLSGLTT DSTRYQLAST
GFGDASAAYQ ERLMTVPVDV HAALQELCLE RRVSVGSVIN FSVHQMLKGF
GNGTHTITAS LHREQNLQNS SPSWVVSPTI VTHENRDGWS VAQAVESIEA
GRGSEKESVT AIDSGSSLVK MGLFDLLVSF VDADDARIPC FDFPLAVIVR
ECDANLSLTL RFSDCLFNEE TICNFTDALN ILLAEAVIGR VTPVADIELL
SAEQKQQLEE WNNTDGEYPS SKRLHHLIEE VVERHEDKIA VVCDERELTY
GELNAQGNSL ARYLRSIGIL PEQLVALFLD KSEKLIVTIL GVWKSGAAYV
PIDPTYPDER VRFVLDDTKA RAIIASNQHV ERLQREVIGD RNLCIIRLEP
LLASLAQDSS KFPAHNLDDL PLTSQQLAYV TYTSGTTGFP KGIFKQHTNV
VNSITDLSAR YGVAGQHHEA ILLFSACVFE PFVRQTLMAL VNGHLLAVIN
DVEKYDADTL LPFIRRHSIT YLNGTASVLQ EYDFSDCPSL NRIILVGENL
TEARYLALRQ RFKNRILNEY GFTESAFVTA LKIFDPESTR KDTSLGRPVR
NVKCYILNPS LKRVPIGATG ELHIGGLGIS KGYLNRPELT PHRFIPNPFQ
TDCEKQLGIN SLMYKTGDLA RWLPNGEVEY LGRADFQIKL RGIRIEPGEI
ETMLAMYPRV RTSLVVSKKL RNGPEETTNE HLVGYYVCDS ASVSEADLLS
FLEKKLPRYM IPTRLVQLSQ IPVNVNGKAD LRALPAVDIS NSTEVRSDLR
GDTEIALGEI WADVLGARQR SVSRNDNFFR LGGHSITCIQ LIARIRQRLS
VSISVEDVFA TRTLERMADL LQNKQQEKCD KPHEAPTELL EENAATDNIY
LANSLQQGFV YHYLKSMEQS DAYVMQSVLR YNTTLSPDLF QRAWKHAQQS
FPALRLRFSW EKEVFQLLDQ DPPLDWRFLY FTDVAAGAVE DRKLEDLRRQ
DLTERFKLDV GRLFRVYLIK HSENRFTCLF SCHHAILDGW SLPLLFEKVH
ETYLQLLHGD NLTSSMDDPY TRTQRYLHAH REDHLDFWAG VVQKINERCD
MNALLNERSR YKVQLADYDQ VQEQRQLTIA LSGDAWLADL RQTCSAQGIT
LHSILQFVWH AVLHAYGGGT HTITGTTISG RNLPILGIER AVGPYINTLP
LVLDHSTFKD KTIMEAIEDV QAKVNVMNSR GNVELGRLHK TDLKHGLFDS
LFVLENYPNL DKSRTLEHQT ELGYSIEGGT EKLNYPLAVI AREVETTGGF
TVSICYASEL FEEVMISELL HMVQDTLMQV ARGLNEPVGS LEYLSSIQLE
QLAAWNATEA EFPDTTLHEM FENEASQKPD KIAVVYEETS LTYRELNERA
NRMAHQLRSD VSPNPNEVIA LVMDKSEHMI VNILAVWKSG GAYVPIDPGY
PNDRIQYILE DTQALAVIAD SCYLPRIKGM AASGTLLYPS VLPANPDSKW
SVSNPSPLSR STDLAYIIYT SGTTGRPKGV TVEHHGVVNL QVSLSKVFGL
RDTDDEVILS FSNYVFDHFV EQMTDAILNG QTLLVLNDGM RGDKERLYRY
IEKNRVTYLS GTPSVVSMYE FSRFKDHLRR VDCVGEAFSE PVFDKIRETF
HGLVINGYGP TEVSITTHKR LYPFPERRMD KSIGQQVHNS TSYVLNEDMK
RTPIGAVGEL YLGGEGVVRG YHNRADVTAE RFIPNPFQSE EDKREGRNSR
LYKTGDLVRW IPGSSGEVEY LGRNDFQVKI RGLRIELGEI EAILSSYHGI
KQSVVIAKDC REGAQKFLVG YYVADAALPS AAIRRFMQSR LPGYMVPSRL
ILVSKFPVTP SGKLDTKALP PAEEESEIDV VPPRSEIERS LCDIWAELLE
MHPEEIGIYS DFFSLGGDSL KSTKLSFMIH ESFNRAVSVS ALFCHRTVEA
QTHLILNDAA DVHEITPIDC NDTQMIPVSR AQERLLFIHE FENGSNAYNI
DAAFELPGSV DASLLEQALR GNLARHEALR TLLVKDHATG IYLQKVLSPD
EAQGMFSVNV DTAKQVERLD QEIASLSQHV FRLDDELPWE ARILKLESGG
LYLILAFHHT CFDAWSLKVF EQELRALYAA LQKTKSAANL PALKAQYKEY
ALYHRRQLSG DRMRNLSDFW LRKLIGLEPL QLITDRPRPV QFKYDGDDLS
IELSKKETEN LRGVAKRCKS SLYVVLVSVY CVMLASYANQ SDVSVGIPVS
HRTHPQFQSV IGFFVNLVVL RVDISQSAIC GLIRRVMKEL VDAQLHQDMP
FQEVTKLLQV DNDPSRHPLV QNVFNFESRA NGEHDARSED EGSLAFNQYR
PVQPVDSVAK FDLNATVTEL ESGLRVNFNY ATSLFNKSTI QGFLHTYEYL
LRQLSELSAE GINEDTQLSL VRPTENGDLH LPLAQSPLAT TAEEQKVASL
NQAFEREAFL AAEKIAVVQG DRALSYADLN GQANQLARYI QSVSCIGADD
GIALMLEKSI DTIICILAIW KAGAAYVPLD PTYPPGRVQL ILEEIKAKAV
LVHSSHASKC ERHGAKVIAV DSPAIETAVS QQSAADLPTI ASLGNLAYII
FTSGTSGKPK GVLVEQKAVL LLRDALRERY FGRDCTKHHG VLFLSNYVFD
FSVEQLVLSV LSGHKLIVPP AEFVADDEFY RMASTHGLSY LSGTPSLLQK
IDLARLDHLQ VVTAAGEELH ATQYEKMRRR FNGPIYNAYG VTETTVYNII
AEFTTNSIFE NALREVLPGT RAYVLNAALQ PVPFDAVGEL YLAGDSVTRG
YLNQPLLTDQ RFIPNPFCKE EDIAMGRFAR LYKTGDLVRS RFNRQQQPQL
EYLGRGDLQI KMRGYRIEIS EVQNVLTSSP GVREGAVVAK YENNDTYSRT
AHSLVGYYTT DNETVSEADI LTFMKARLPT YMVPSHLCCL EGALPVTING
KLDVRRLPEI INDSAQSSYS PPRNIIEAKM CRLWESALGM ERCGIDDDLF
KLGGDSITSL HLVAQIHNQV GCKITVRDIF EHRTARALHD HVFMKDSDRS
NVTQFRTEQG PVIGEAPLLP IQDWFLSKAL QHPMYWNHTF YVRTPELDVD
SLSAAVRDLQ QYHDVFRMRL KREEVGFVQS FAEDFSPAQL RVLNVKDVDG
SAAVNEILDG WQSGFNLENG PIGSIGYLHG YEDRSARVWF SVHHMAIDTV
SWQILVRDLQ TLYRNGSLGS KGSSFRQWAE AIQNYKASDS ERNHWNKLVM
ETASSISALP TSTGSRVRLS RSLSPEKTAS LIQGGIDRQD VSVYDSLLTS
VGLALQHIAP TGPSMVTIEG HGREEVDQTL DVSRTMGWFT TMYPFEIPRL
STENIVQGVV AVSERFRQVP ARGVGYGTLY GYTQHPLPQV TVNYLGQLAR
KQSKPKEWVL AVGDNEFEYG LMTSPEDKDR SSSAVDVTAV CIDGTMIIDV
DSAWSLEESE QFISSIEEGL NKILDGRASQ QTSRFPDVPQ PAETYTPYFE
YLEPPRQGPT LFLLPPGEGG AESYFNNIVK RLRQTNMVVF NNYYLHSKRL
RTFEELAEMY LDQVRGIQPH GPYHFIGWSF GGILAMEMSR RLVASDEKIG
FLGIIDTYFN VRGATRTIGL GDTEILDPIH HIYNPDPANF QRLPSATDRI
VLFKAMRPNN KYESENQRRL YEYYDGTRLN GLDSLLPSDS DVQLVPLTDD
THFSWVGNPQ QVEQMCATIK EHLARYLIAW RASRGGPVPV EKMSKGEELF
TGVVPILVEL DGDVNGHKFS VSGEGEGDAT YGKLTLKFIC TTGKLPVPWP
TLVTTLTYGV QCFSRYPDHM KRHDFFKSAM PEGYVQERTI FFKDDGNYKT
RAEVKFEGDT LVNRIELKGI DFKEDGNILG HKLEYNYNSH NVYIMADKQK
NGIKANFKTR HNIEDGGVQL ADHYQQNTPI GDGPVLLPDN HYLSTQSALS
KDPNEKRDHM VLLEFVTAAG ITHGMVELYK HHHHHH
```

Amino acid coverage showing parts of the ACVS reference protein sequence (8-1) are identified by detected peptides from MS/MS. Segments with identified peptides are marked in yellow, and modified amino acids are in green. The sequence coverage percentages of 8-1 is 20%.

**Fig. S34**

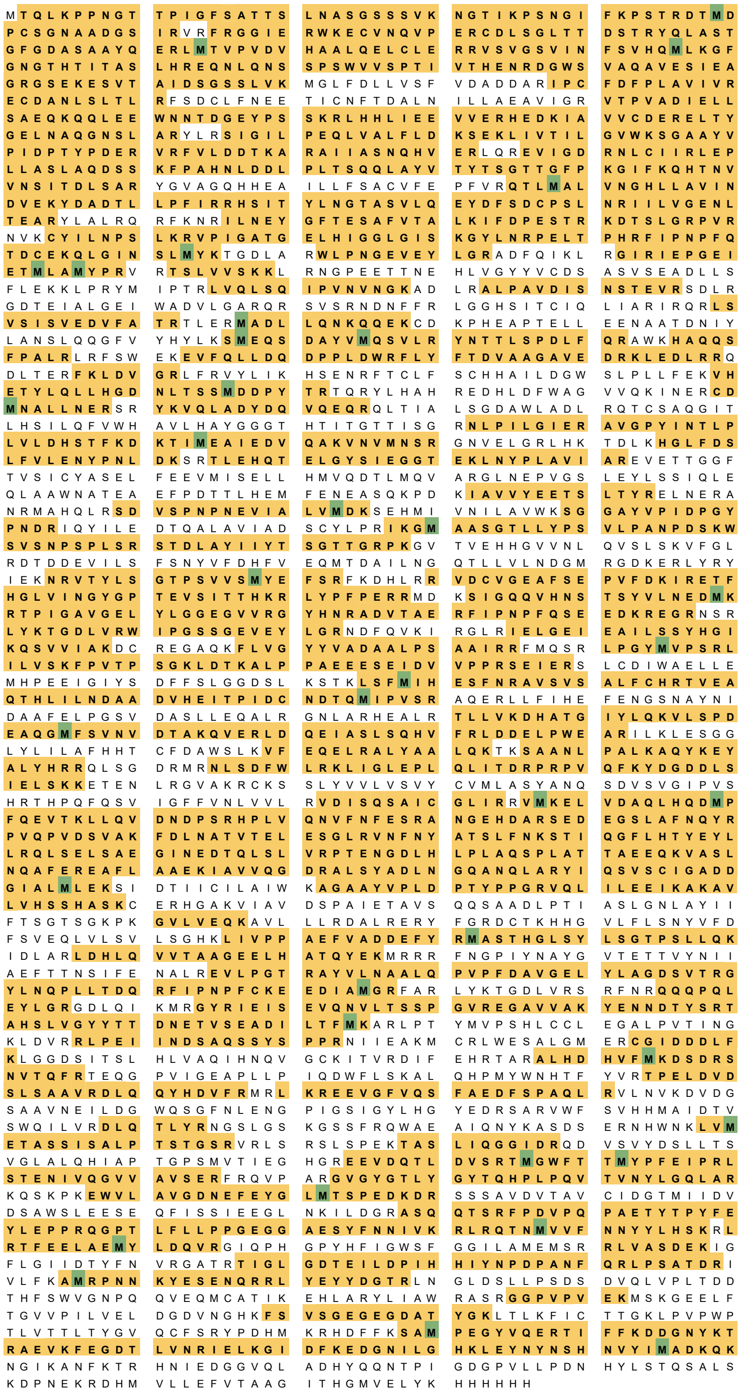

Amino acid coverage showing parts of the ACVS reference protein sequence (8-2) are identified by detected peptides from MS/MS. Segments with identified peptides are marked in yellow, and modified amino acids are in green. The sequence coverage percentages of 8-2 is 60%.

**Fig. S35**

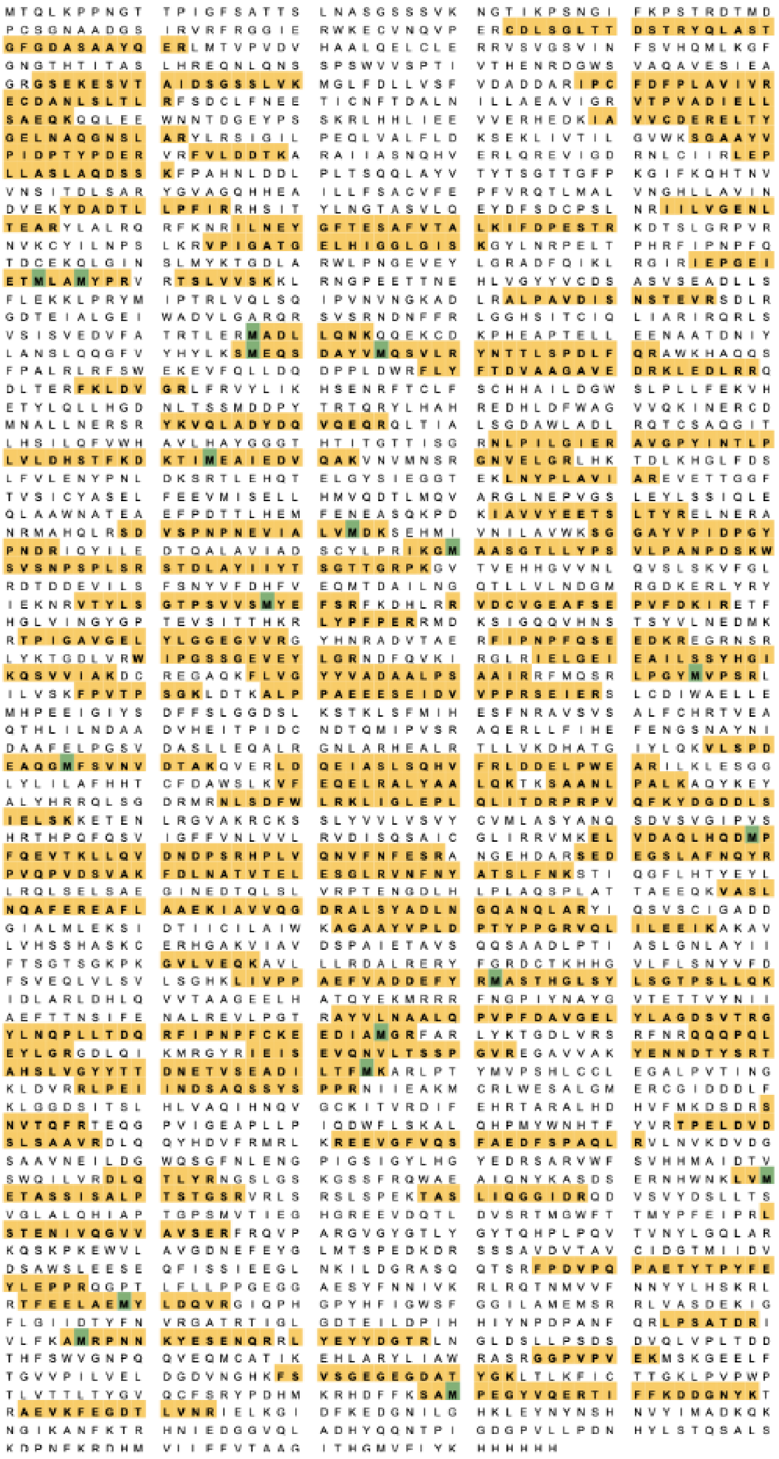

Amino acid coverage showing parts of the ACVS reference protein sequence (10-1) are identified by detected peptides from MS/MS. Segments with identified peptides are marked in yellow, and modified amino acids are in green. The sequence coverage percentage of 10-1 is 33%.

**Fig. S36.**

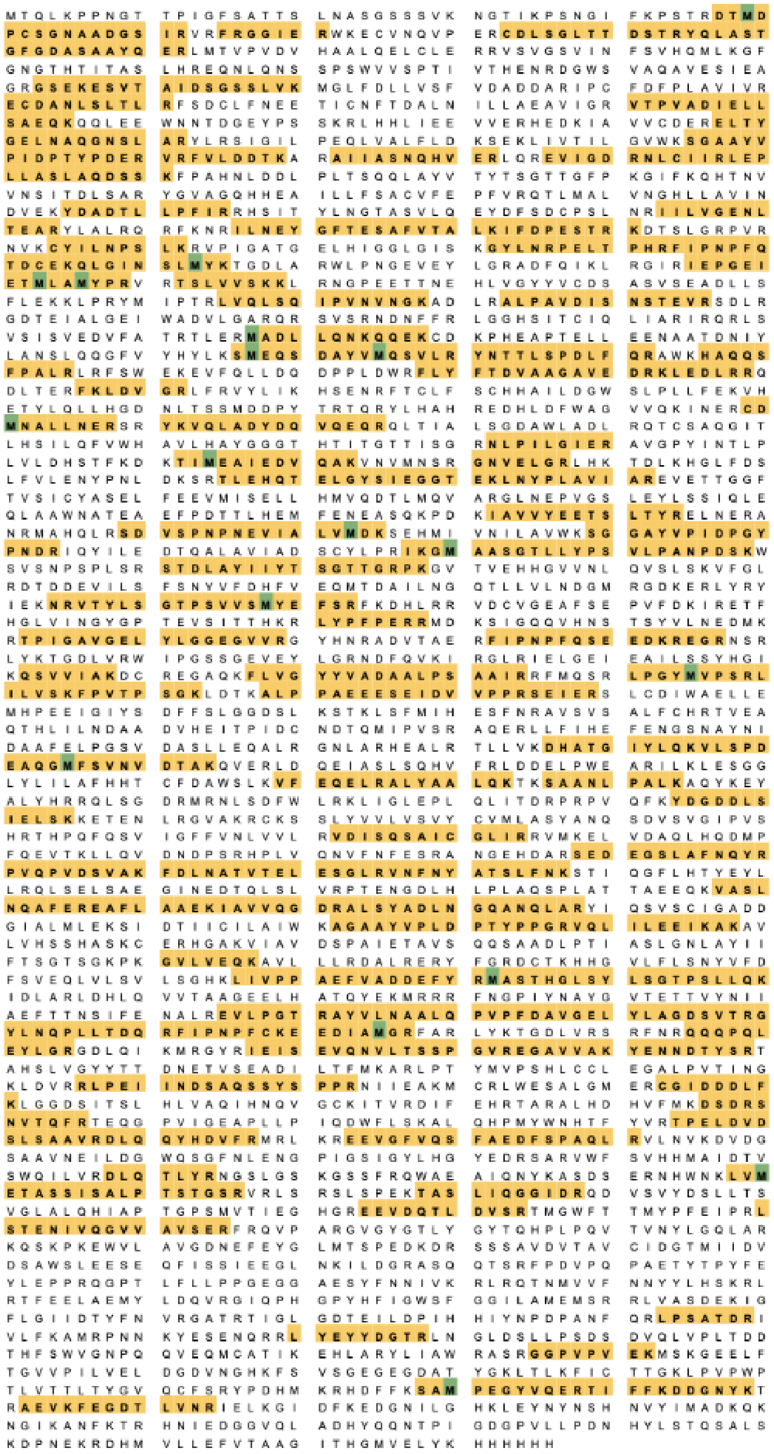

Amino acid coverage showing parts of the ACVS reference protein sequence (10-2) are identified by detected peptides from MS/MS. Segments with identified peptides are marked in yellow, and modified amino acids are in green. The sequence coverage percentages of 10-2 is 31%.

**Fig. S37**

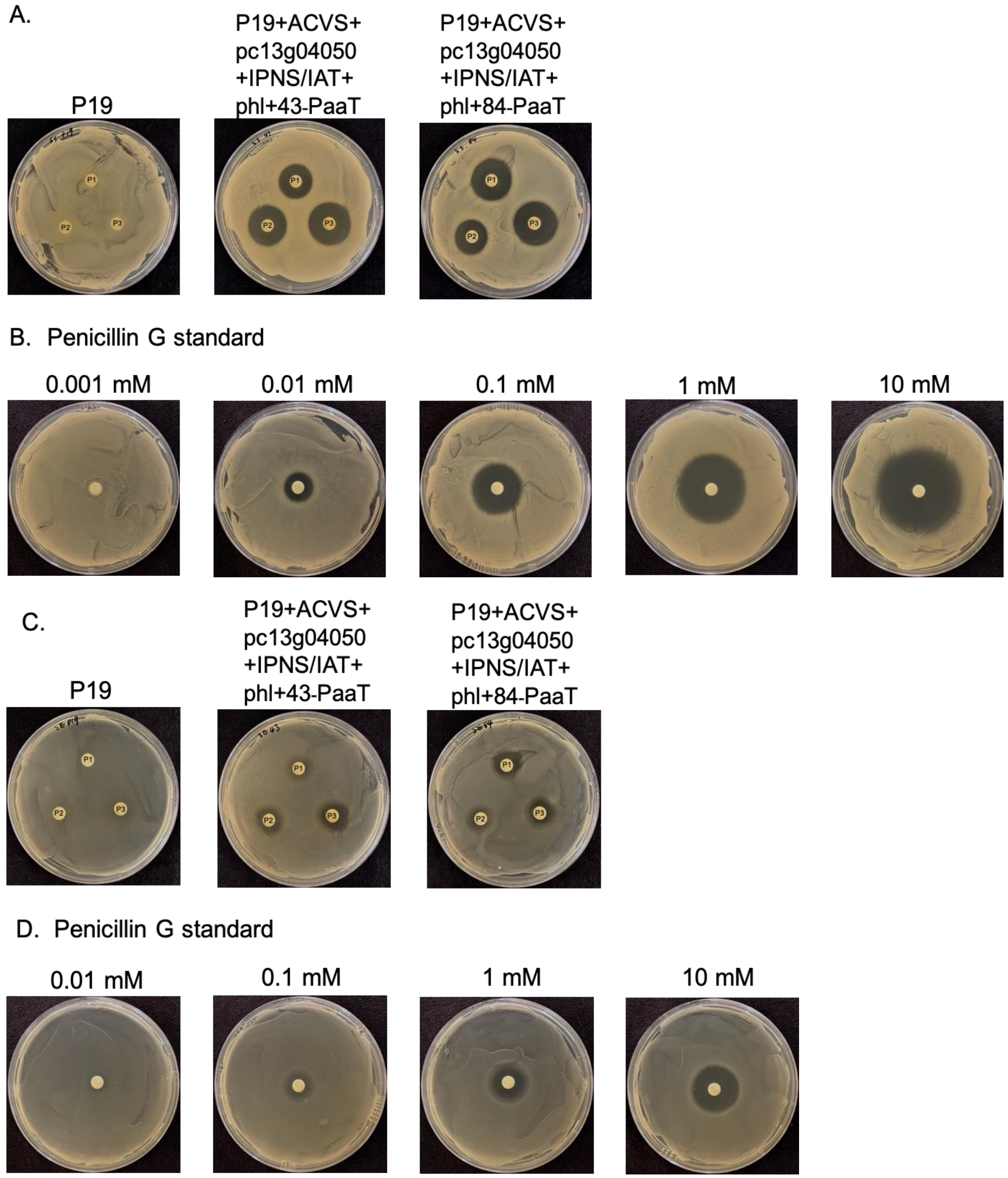

**Growth inhibition assay on *Staphylococcus saprophyticus* and *Staphylococcus epidermidis*.** Plant extracts were extracted from three infiltrated leaves of three individual *N. benthamiana* plants expressing the penicillin G biosynthetic genes with the PaaT transporter tagged with GFP at N-terminus (43-PaaT) or C-terminus (84-PaaT) along with expressing P19 alone. Plant extracts and Penicillin G standards with different concentrations were applied to the bacterial lawn of *Staphylococcus saprophyticus* (A and B) and *Staphylococcus epidermidis* (C and D), respectively. P1, P2, and P3: Plant extracts from three individual plants.

**Fig. S38**

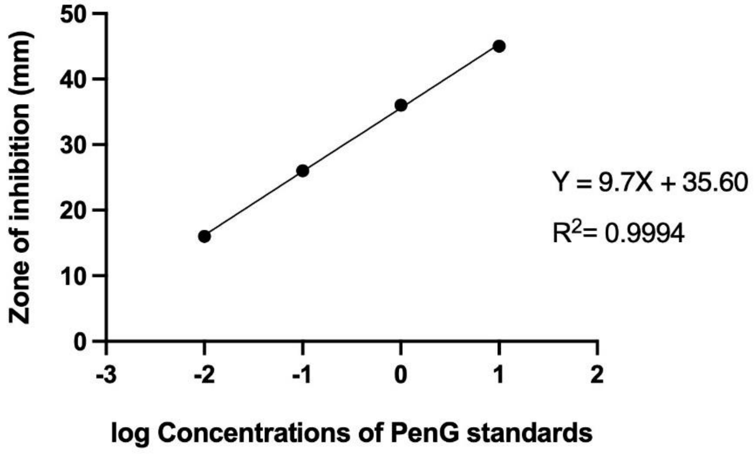

**Linear regression analysis of inhibition zone diameters against log-transformed penicillin G concentrations derived from the *Staphylococcus saprophyticus* bioassay.** The diameters of the zone of inhibition (mm) were measured against a bacterial lawn of *Staphylococcus saprophyticus* (Fig.37B). Data points represent the relationship between the measured zones and the $\log_{10}$ concentrations of penicillin G. The solid line represents the linear regression model (Y = 9.700X + 35.60) with a coefficient of determination ($R^2$) of 0.9994.

**Table S1.**

| **Name** | **Cultivation medium and conditions** | **Reference / Supplier** |
|---|---|---|
| *E. coli* TOP 10 | LB agar + appropriate antibiotic (16 hours, 37ºC) | Invitrogen |
| *E. coli ccdB* survival | LB agar + appropriate antibiotic (16 hours, 37ºC) | Invitrogen |
| *E. coli* EPI400 | LB agar + appropriate antibiotic (16 hours, 37ºC) | Biosearch Technologies |
| *S. cerevisiae* YPH 499 | YPAD agar (2 -3 days, 28ºC)<br>YPAD medium (16 hours, 30 ºC, 200 rpm)<br>SM-ura agar (2 – 3 days, 30 ºC)<br>SM-ura medium (16 hours, 30 ºC, 200 rpm) | ATCC |
| *P. rubens* NRRL 1951 | PDB (4 days, 25oC, 120 rpm) | ARS Culture Collection (NRRL) |
| *Agrobacterium* GV3101 | LB agar + 25µg/ml Rifampicin + 50 µg/ml Gentamicin (48 hours, 28ºC) | GoldBio |
| *Staphylococcus saprophyticus* ATCC 49007 | LB agar (16 hours, 37ºC) | ATCC |
| *Staphylococcus epidermidis* ATCC 1228 | LB agar (16 hours, 37ºC) | ATCC |

Microorganisms used in this study and culture conditions.

**Table S2.**

| **Medium Name** | **Ingredients** |
| --- | --- |
| LB agar | 5 g/L yeast extract, 10 g/L tryptone, 5 g/L NaCl, 15 g/L agar |
| YPAD agar | 10 g/L yeast extract, 10 g/L tryptone, 0.3 g/L adenine, 20 g/L D (+)-glucose monohydrate, 15 g/L agar |
| YPAD medium | 10 g/L yeast extract, 10 g/L tryptone, 0.3 g/L adenine, 20 g/L D (+)-glucose monohydrate |
| SM-ura agar | 1,7 g/L yeast nitrogen base, 20 g/L D (+)-glucose monohydrate, 5 g/L ammonium sulphate, 0.77 g/L complete supplement mixture minus uracil (Q biogene), 25 g/L agar |
| SM-ura media | 1,7 g/L yeast nitrogen base, 20 g/L D (+)-glucose monohydrate, 5 g/L ammonium sulphate, 0.77 g/L complete supplement mixture minus uracil (Q biogene) |
| PDB agar | 24 g/L potato dextrose broth, 15 g/L agar |
| PDB medium | 24 g/L potato dextrose broth |
| LB Broth | 10 g/L NaCl, 10 g/L Tryptone, 5 g/L Yeast Extract (Sigma-Aldrich) |
| *N. benthamiana* infiltration buffer | 10 mM $MgCl_2$, 10 mM MES (pH 5.6), 500 µM acetosyringone, 0.5% glucose, and poloxamer 188 (1%, w/v) dissolved in sterile water |

Culture media recipes used in this study.

**Table S3.**

| Primer Name | Sequence (5' – 3') | Description |
|---|---|---|
| M13-F | GTTGTAAAACGACGGCCAGT | General use |
| M13-R | CCAGGAAACAGCTATGACCATG | General use |
| pcbAB_PR_F1-F | ttaacagctatacaagtttgtacaaaaaactgatgactcaactgaagccac | Forward primer to amplify pcbAB-native-F1 cDNA with an overhang homologous to the pDEST/URA/Ub10 vector |
| pcbAB_PR_F1-R | cgaatgatcgagtaccag | Reverse primer to amplify pcbAB-native-F1 cDNA with an overhang homologous to the pcbAB-native-F2 |
| pcbAB_PR_F2-F | caacactctaccgctggt | Forward primer to amplify pcbAB-native-F2 cDNA with an overhang homologous to the pcbAB-native-F1 |
| pcbAB_PR_F2-R | cgtcgtctgccccaatac | Reverse primer to amplify pcbAB-native-F2 cDNA with an overhang homologous to the pcbAB-native-F3 |
| pcbAB_PR_F3-F | catacagtccgtgtcctgta | Forward primer to amplify pcbAB-native-F3 cDNA with an overhang homologous to the pcbAB-native-F2 |
| pcbAB_PR_F3-R | gcccccccctcgaggcgcgccaagctatcaaatagcgagcgaggtgttc | Reverse primer to amplify pcbAB-native-F3 cDNA with an overhang homologous to the linker region between pcbAB-native-F3 and eGFP |
| pcbAB_F1 | ATGACTCAACTGAAGCCACCGAACG | RT-PCR confirmation of pcbAB expression |
| PcbAB_R1 | TACGAGCATCGTCTGCATCG | |
| pcbAB_F2 | ACTTCATCGGATGGAGCTTC | RT-PCR confirmation of pcbAB expression |
| pcbAB_R2 | TTCACCCTCTCCACTGACAG | |
| IPNS-F | gtcaattccggagaaaaaggcag | RT-PCR confirmation of IPNS expression |
| IPNS-R | ctaggatcccaaggttgtacag | |
| IAT-F | gggaaggttggatttaacagtgc | RT-PCR confirmation of IAT expression |
| IAT-R | gcgttgagcgcagacc | |
| NpgA-F | caagagatctttcttcacc | RT-PCR confirmation of npgA expression |
| NpgaA-R | cttcttgaaactcttcaacc | |
| PaaT_F | TCCACGGTCAGATCAGGCTC | RT-PCR confirmation of paaT expression |
| PaaT_R | ACTGGTCACCAAGACGTTCG | |

| | | |
|---|---|---|
| PenM_F | AAGGATGGTGAGGAGACACC | RT-PCR confirmation of penM expression |
| PenM_R | GAATTTCCCCAACCAAGACC | |
| Pc13g04050_F | ATGGCTTCTATCCAGGTGTG | RT-PCR confirmation of pc13g04050 expression |
| Pc13g04050_R | AGCTTGGTAAATCCCGTAGG | |
| NbEF_F | ACTGCACTGTGATTGATGCC | RT-PCR confirmation of *EF-1 α* expression |
| NbEF_R | GACACCAGTTTCCACACGAC | |
| GFPtot-R | GTATAGTTCATCCATGCCATG | RT-PCR confirmation of eGFP expression |
| Linker+eGFP-F | ttgatagcttggcgcgcc | Forward primer to amplify the linker region +eGFP from the pMDC84 vector with an overhang homologous to the linker region +eGFP |
| NYV_pcbAB_egfp-R | tgaattataccactttgtacaagaaagctgttagtggtggtggtg | Reverse primer to amplify linker region +eGFP from the pMDC84 vector with an overhang homologous to the pDEST/URA/Ub10 vector |
| AscI_phl_F | TTAATATGGCGCGCCTATGGTTTTTTACCTCCAAAGG | Forward and reverse primers were used to amplify coding regions and add two restriction enzyme recognition sites on both ends of the amplicons |
| PacI_phl_R | TTAATTATTAATTAATTAGATCTTGCTACCAGCCTTCTCC | |
| PacI_pc13g04050_F | TTGGAATTTAATTAAATGGCTTCTATCCAGGTGTGG | Forward and reverse primers were used to amplify coding regions and add two restriction enzyme recognition sites on both ends of the amplicons |
| AscI_pc13g04050_R | TTGGTTAAGGCGCGCCAAAGCTTGGTAAATCCCGTAGG | |
| AscI_NpgA_F | TTAAGGTTGGCGCGCCATGGTTCAAGATACTTCTTCTGC | Forward and reverse primers were used to amplify coding regions and add two restriction enzyme recognition sites on both ends of the amplicons |
| PacI_NpgA_R | TTGGTTATATTAATTAAGAAAGACAATTACAAACACCAGTAGC | |

Oligonucleotides used in this study.

**Table S4.**

| Enzyme Name | Protein Sequence | (Codon optimized) coding sequence |
|---|---|---|
| ACVS (native) | MTQLKPPNGTTPIGFSATTSLN<br>ASGSSSVKNGPSNPAMAYFKPS<br>TRDTMDPCSGNAADGSIRVRF<br>RGGIERWKECVNQVPERCDLS<br>GLTTDSTRYQLASTGFGDASA<br>AYQERLMTVPVDVHAALQEL<br>CLERRVSVGSVINFSVHQMLK<br>GFGNGTHTITASLHREQNLQNS<br>SPSWVVSPTIVTHENRDGWSV<br>AQAVESIEAGRGSEKESVTAID<br>SGSSLVKMGLFDLLVSFVDAD<br>DARIPCFDFPLAVIVRECDANL<br>SLTLRFSDCLFNEETICNFTDAL<br>NILLAEAVIGRVTPVADIELLSA<br>EQKQQLEEWNNTDGEYPSSKR<br>LHHLIEEVVERHEDKIAVVCDE<br>RELTYGELNAQGNSLARYLRSI<br>GILPEQLVALFLDKSEKLIVTIL<br>GVWKSGAAYVPIDPTYPDERV<br>RFVLDDTKARAIIASNQHVERL<br>QREVIGDRNLCIIRLEPLLASLA<br>QDSSKFPAHNLDDLPLTSQQLA<br>YVTYTSGTTGFPKGIFKQHTN<br>VVNSITDLSARYGVAGQHHEAI<br>LLFSACVFEPFVRQTLMALVN<br>GHLLAVINDVEKYDADTLLPFI<br>RRHSITYLNGTASVLQEYDFSD<br>CPSLNRIILVGENLTEARYLALR<br>QRFKNRILNEYGFTESAFVTAL<br>KIFDPESTRKDTSLGRPVRNVK<br>CYILNPSLKRVPIGATGELHIGG<br>LGISKGYLNRPELTPHRFIPNPF<br>QTDCEKQLGINSLMYKTGDLA | See Table S5 for fragment sequences used in this study |

| | |
|---|---|
| | RWLPNGEVEYLGRADFQIKLR<br>GIRIEPGEIETMLAMYPRVRTSL<br>VVSKKLRNGPEETTNEHLVGY<br>YVCDSASVSEADLLSFLEKKLP<br>RYMIPTRLVQLSQIPVNVNGKA<br>DLRALPAVDISNSTEVRSDLRG<br>DTEIALGEIWADVLGARQRSV<br>SRNDNFFRLGGHSITCIQLIARI<br>RQRLSVSISVEDVFATRTLERM<br>ADLLQNKQQEKCDKPHEAPTE<br>LLEENAATDNIYLANSLQQGF<br>VYHYLKSMEQSDAYVMQSVL<br>RYNTTLSPDLFQRAWKHAQQS<br>FPALRLRFSWEKEVFQLLDQD<br>PPLDWRFLYFTDVAAGAVEDR<br>KLEDLRRQDLTERFKLDVGRL<br>FRVYLIKHSENRFTCLFSCHHA<br>ILDGWSLPLLFEKVHETYLQLL<br>HGDNLTSSMDDPYTRTQRYLH<br>AHREDHLDFWAGVVQKINER<br>CDMNALLNERSRYKVQLADY<br>DQVQEQRQLTIALSGDAWLAD<br>LRQTCSAQGITLHSILQFVWHA<br>VLHAYGGGTHTITGTTISGRNL<br>PILGIERAVGPYINTLPLVLDHS<br>TFKDKTIMEAIEDVQAKVNVM<br>NSRGNVELGRLHKTDLKHGLF<br>DSLFVLENYPNLDKSRTLEHQT<br>ELGYSIEGGTEKLNYPLAVIAR<br>EVETTGGFTVSICYASELFEEV<br>MISELLHMVQDTLMQVARGL<br>NEPVGSLEYLSSIQLEQLAAWN<br>ATEAEFPDTTLHEMFENEASQ<br>KPDKIAVVYEETSLTYRELNER<br>ANRMAHQLRSDVSPNPNEVIA<br>LVMDKSEHMIVNILAVWKSGG<br>AYVPIDPGYPNDRIQYILEDTQ<br>ALAVIADSCYLPRIKGMAASGT<br>LLYPSVLPANPDSKWSVSNPSP<br>LSRSTDLAYIIYTSGTTGRPKG |

| | |
|---|---|
| | VTVEHHGVVNLQVSLSKVFGL<br>RDTDDEVILSFSNYVFDHFVEQ<br>MTDAILNGQTLLVLNDGMRG<br>DKERLYRYIEKNRVTYLSGTPS<br>VVSMYEFSRFKDHLRRVDCVG<br>EAFSEPVFDKIRETFHGLVING<br>YGPTEVSITTHKRLYPFPERRM<br>DKSIGQQVHNSTSYVLNEDMK<br>RTPIGSVGELYLGGEGVVRGY<br>HNRADVTAERFIPNPFQSEEDK<br>REGRNSRLYKTGDLVRWIPGSS<br>GEVEYLGRNDFQVKIRGLRIEL<br>GEIEAILSSYHGIKQSVVIAKDC<br>REGAQKFLVGYYVADAALPSA<br>AIRRFMQSRLPGYMVPSRLILV<br>SKFPVTPSGKLDTKALPPAEEE<br>SEIDVVPPRSEIERSLCDIWAEL<br>LEMHPEEIGIYSDFFSLGGDSL<br>KSTKLSFMIHESFNRAVSVSAL<br>FCHRTVEAQTHLILNDAADVH<br>EITPIDCNDTQMIPVSRAQERL<br>LFIHEFENGSNAYNIDAAFELP<br>GSVDASLLEQALRGNLARHEA<br>LRTLLVKDHATGIYLQKVLSPD<br>EAQGMFSVNVDTAKQVERLD<br>QEIASLSQHVFRLDDELPWEA<br>RILKLESGGLYLILAFHHTCFD<br>AWSLKVFEQELRALYAALQKT<br>KSAANLPALKAQYKEYALYHR<br>RQLSGDRMRNLSDFWLRKLIG<br>LEPLQLITDRPRPVQFKYDGDD<br>LSIELSKKETENLRGVAKRCKS<br>SLYVVLVSVYCVMLASYANQS<br>DVSVGIPVSHRTHPQFQSVIGF<br>FVNLVVLRVDISQSAICGLIRRV<br>MKELVDAQLHQDMPFQEVTK<br>LLQVDNDPSRHPLVQNVFNFE<br>SRANGEHDARSEDEGSLAFNQ<br>YRPVQPVDSVAKFDLNATVTE<br>LESGLRVNFNYATSLFNKSTIQ |

| | |
|---|---|
| | GFLHTYEYLLRQLSELSAEGIN<br>EDTQLSLVRPTENGDLHLPLA<br>QSPLATTAEEQKVASLNQAFER<br>EAFLAAEKIAVVQGDRALSYA<br>DLNGQANQLARYIQSVSCIGA<br>DDGIALMLEKSIDTIICILAIWK<br>AGAAYVPLDPTYPPGRVQLILE<br>EIKAKAVLVHSSHASKCERHG<br>AKVIAVDSPAIETAVSQQSAAD<br>LPTIASLGNLAYIIFTSGTSGKP<br>KGVLVEQKAVLLLRDALRERY<br>FGRDCTKHHGVLFLSNYVFDF<br>SVEQLVLSVLSGHKLIVPPAEF<br>VADDEFYRMASTHGLSYLSGT<br>PSLLQKIDLARLDHLQVVTAA<br>GEELHATQYEKMRRRFNGPIY<br>NAYGVTETTVYNIIAEFTTNSIF<br>ENALREVLPGTRAYVLNAALQ<br>PVPFDAVGELYLAGDSVTRGY<br>LNQPLLTDQRFIPNPFCKEEDIA<br>MGRFARLYKTGDLVRSRFNRQ<br>QQPQLEYLGRGDLQIKMRGYR<br>IEISEVQNVLTSSPGVREGAVVA<br>KYENNDTYSRTAHSLVGYYTT<br>DNETVSEADILTFMKARLPTY<br>MVPSHLCCLEGALPVTINGKL<br>DVRRLPEIINDSAQSSYSPPRNI<br>IEAKMCRLWESALGMERCGID<br>DDLFKLGGDSITSLHLVAQIHN<br>QVGCKITVRDIFEHRTARALHD<br>HVFMKDSDRSNVTQFRTEQGP<br>VIGEAPLLPIQDWFLSKALQHP<br>MYWNHTFYVRTPELDVDSLSA<br>AVRDLQQYHDVFRMRLKREE<br>VGFVQSFAEDFSPAQLRVLNV<br>KDVDGSAAVNEILDGWQSGFN<br>LENGPIGSIGYLHGYEDRSARV<br>WFSVHHMAIDTVSWQILVRDL<br>QTLYRNGSLGSKGSSFRQWAE<br>AIQNYKASDSERNHWNKLVM |

| | | |
|---|---|---|
| | ETASSISALPTSTGSRVRLSRSL<br>SPEKTASLIQGGIDRQDVSVYD<br>SLLTSVGLALQHIAPTGPSMVT<br>IEGHGREEVDQTLDVSRTMGW<br>FTTMYPFEIPRLSTENIVQGVV<br>AVSERFRQVPARGVGYGTLYG<br>YTQHPLPQVTVNYLGQLARK<br>QSKPKEWVLAVGDNEFEYGL<br>MTSPEDKDRSSSAVDVTAVCID<br>GTMIIDVDSAWSLEESEQFISSI<br>EEGLNKILDGRASQQTSRFPDV<br>PQPAETYTPYFEYLEPPRQGPT<br>LFLLPPGEGGAESYFNNIVKRL<br>RQTNMVVFNNYYLHSKRLRTF<br>EELAEMYLDQVRGIQPHGPYH<br>FIGWSFGGILAMEMSRRLVASD<br>EKIGFLGIIDTYFNVRGATRTIG<br>LGDTEILDPIHHIYNPDPANFQR<br>LPSATDRIVLFKAMRPNNKYES<br>ENQRRLYEYYDGTRLNGLDSL<br>LPSDSDVQLVPLTDDTHFSWV<br>GNPQQVEQMCATIKEHLARY | |
| IPNS | MASTPKANVPKIDVSPLFGDN<br>MEEKMKVARAIDAASRDTGFF<br>YAVNHGVDVKRLSNKTREFHF<br>SITDEEKWDLAIRAYNKEHQD<br>QIRAGYYLSIPEKKAVESFCYL<br>NPNFKPDHPLIQSKTPTHEVNV<br>WPDEKKHPGFREFAEQYYWD<br>VFGLSSALLRGYALALGKEED<br>FFSRHFKKEDALSSVVLIRYPY<br>LNPYPPAAIKTAEDGTKLSFEW<br>HEDVSLITVLYQSDVANLQVE<br>MPQGYLDIEADDNAYLVNCGS<br>YMAHITNNYYPAPIHRVKWVN<br>EERQSLPFFVNLGFNDTVQPW<br>DPSKEDGKTDQRPISYGDYLQ<br>NGLVSLINKNGQT | ATGGCTTCAACTCCAAAAGCCAATGTACCAAAAATAGATGTGTCTCCCCTTTTCGGGGACAATA<br>TGGAGGAAAAGATGAAGGTTGCCCGAGCTATTGATGCTGCCAGCCGTGACACCGGGTTCTTCT<br>ACGCAGTCAATCATGGTGTGGATGTGAAACGACTATCCAATAAGACAAGGGAATTCCACTTTTC<br>TATCACAGATGAAGAAAAGTGGGATCTCGCTATTAGAGCCTACAATAAAGAACACCAAGATCA<br>GATAAGGGCAGGATACTACCTGTCAATTCCGGAGAAAAAGGCAGTTGAATCATTCTGCTACCTT<br>AACCCCAACTTTAAACCTGATCATCCTCTTATCCAGTCTAAAACTCCCACTCATGAAGTAAATGT<br>TTGGCCGGACGAGAAGAAACATCCTGGTTTCCGCGAGTTTGCAGAGCAATATTACTGGGATGT<br>GTTCGGGCTCTCGTCTGCTTTGCTGAGAGGTTATGCTCTTGCGTTAGGAAAGGAGGAAGATTTT<br>TTTAGCAGGCACTTTAAGAAGGAAGACGCTTTATCCAGTGTTGTTCTTATTCGTTATCCTTACCT<br>TAATCCATATCCACCTGCAGCAATTAAGACGGCTGAGGATGGCACCAAATTGAGTTTTGAATGG<br>CATGAAGATGTTTCTTTGATTACTGTCCTGTATCAGTCAGATGTTGCTAACTTGCAGGTTGAGAT<br>GCCACAGGGTTACCTCGATATTGAGGCGGACGACAACGCTTACCTTGTAAATTGTGGTAGCTAT<br>ATGGCACATATCACAAACAATTATTATCCAGCTCCGATACACCGGGTCAAGTGGGTGAATGAGG<br>AGAGACAATCCTTGCCTTTTTTTGTCAATTTGGGATTTAATGATACTGTACAACCTTGGGATCCT<br>AGTAAGGAAGACGGTAAGACCGATCAAAGACCAATCTCTTATGGCGATTATCTACAGAACGGA<br>TTAGTTAGTCTAATTAACAAAAATGGACAAACATGA |
| IAT | MLHILCQGTPFEIGYEHGSAAK<br>AVIARSIDFAVDLIRGKTKKTD | ATGTTACACATACTCTGTCAAGGTACTCCCTTTGAAATTGGCTACGAACATGGATCTGCTGCCA<br>AAGCCGTTATAGCTAGAAGTATTGACTTCGCCGTCGATTTGATTCGAGGGAAGACAAAGAAGA |

| | | |
|---|---|---|
| | EELKQVLSQLGRVIEERWPKY<br>YEEIRGIAKGAERDVSEIVMLN<br>TRTEFAYGLKAARDGCTTAYC<br>QLPNGALQGQNWDFFSATKEN<br>LIRLTIRQAGLPTIKFITEAGIIG<br>KVGFNSAGVAVNYNALHLQG<br>LRPTGVPSHIALRIALESTSPSQ<br>AYDRIVEQGGMAASAFIMVGN<br>GHEAFGLEFSPTSIRKQVLDAN<br>GRMVHTNHCLLQHGKNEKEL<br>DPLPDSWNRHQRMEFLLDGFD<br>GTKQAFAQLWADEDNYPFSIC<br>RAYEEGKSRGATLFNIIYDHAR<br>REATVRLGRPTNPDEMFVMRF<br>DEEDERSALNARL | CTGATGAAGAGCTTAAACAGGTATTGTCGCAACTGGGGAGAGTTATTGAGGAAAGATGGCCAA<br>AATACTATGAAGAGATTCGTGGTATTGCAAAGGGAGCTGAAAGAGATGTCTCCGAGATTGTTAT<br>GCTTAATACACGCACGGAATTTGCATACGGTCTAAAGGCAGCACGTGATGGATGTACCACTGCT<br>TATTGTCAACTTCCAAATGGAGCCCTGCAAGGTCAAAACTGGGATTTCTTTTCTGCTACAAAAG<br>AAAATCTGATAAGATTAACGATCCGTCAGGCAGGATTACCGACAATCAAATTCATAACCGAGGC<br>TGGAATCATTGGGAAGGTTGGATTTAACAGTGCTGGGGTAGCAGTTAATTATAACGCCCTTCAT<br>CTTCAGGGTCTTCGACCCACCGGAGTTCCTTCACATATTGCTTTGAGGATAGCGCTCGAAAGTA<br>CTTCTCCTTCCCAGGCTTATGACCGGATTGTGGAGCAAGGTGGAATGGCAGCTAGCGCTTTTAT<br>TATGGTGGGAAATGGTCACGAGGCATTTGGTTTGGAATTCTCACCTACATCTATTCGAAAGCAG<br>GTGTTGGATGCTAATGGTAGGATGGTTCATACAAACCATTGCTTGCTTCAACACGGCAAAAATG<br>AGAAGGAACTCGATCCATTACCGGATTCATGGAATCGGCACCAGAGAATGGAGTTCCTATTGG<br>ATGGGTTCGATGGTACTAAACAAGCATTTGCCCAGCTATGGGCTGACGAAGATAATTATCCTTTT<br>TCAATCTGCCGCGCTTACGAGGAAGGCAAGAGCAGAGGCGCGACTCTGTTCAATATCATCTAC<br>GACCATGCACGTAGAGAAGCAACTGTGAGGTTGGGCCGGCCAACCAACCCTGATGAGATGTTT<br>GTCATGAGGTTTGACGAGGAAGATGAGAGGTCTGCGCTCAACGCTAGGCTTTGA |
| NpgA | MVQDTSSASTSPILTRWYIDTR<br>PLTASTAALPLLETLQPADQISV<br>QKYYHLKDKHMSLASNLLKY<br>LFVHRNCRIPWSSIVISRTPDPH<br>RRPCYIPPSGSQEDSFKDGYTG<br>INVEFNVSHQASMVAIAGTAFT<br>PNSGGDSKLKPEVGIDITCVNE<br>RQGRNGEERSLESLRQYIDIFS<br>EVFSTAEMANIRRLDGVSSSSL<br>SADRLVDYGYRLFYTYWALKE<br>AYIKMTGEALLAPWLRELEFS<br>NVVAPAAVAESGDSAGDFGEP<br>YTGVRTTLYKNLVEDVRIEVA<br>ALGGDYLFATAARGGGIGASS<br>RPGGGPDGSGIRSQDPWRPFK<br>KLDIERDIQPCATGVCNCLS | ATGGTTCAAGATACTTCTTCTGCTTCTACTTCTCCTATTCTTACTAGATGGTATATTGATACTAGA<br>CCTCTTACTGCTTCTACTGCTGCTCTTCCTCTTCTTGAAACTCTTCAACCTGCTGATCAAATTTC<br>TGTTCAAAAGTATTATCATCTTAAGGATAAGCATATGTCTCTTGCTTCTAATCTTCTTAAGTATCT<br>TTTTGTTCATAGAAATTGTAGAATTCCTTGGTCTTCTATTGTTATTTCTAGAACTCCTGATCCTCA<br>TAGAAGACCTTGTTATATTCCTCCTTCTGGTTCTCAAGAAGATTCTTTTAAGGATGGTTATACTG<br>GTATTAATGTTGAATTTAATGTTTCTCATCAAGCTTCTATGGTTGCTATTGCTGGTACTGCTTTTA<br>CTCCTAATTCTGGTGGTGATTCTAAGCTTAAGCCTGAAGTTGGTATTGATATTACTTGTGTTAATG<br>AAAGACAAGGTAGAAATGGTGAAGAAAGATCTCTTGAATCTCTTAGACAATATATTGATATTTTT<br>TCTGAAGTTTTTTCTACTGCTGAAATGGCTAATATTAGAAGACTTGATGGTGTTTCTTCTTCTTC<br>TCTTTCTGCTGATAGACTTGTTGATTATGGTTATAGACTTTTTTATACTTATTGGGCTCTTAAGGA<br>AGCTTATATTAAGATGACTGGTGAAGCTCTTCTTGCTCCTTGGCTTAGAGAACTTGAATTTTCTA<br>ATGTTGTTGCTCCTGCTGCTGTTGCTGAATCTGGTGATTCTGCTGGTGATTTTGGTGAACCTTAT<br>ACTGGTGTTAGAACTACTCTTTATAAGAATCTTGTTGAAGATGTTAGAATTGAAGTTGCTGCTCT<br>TGGTGGTGATTATCTTTTTGCTACTGCTGCTAGAGGTGGTGGTATTGGTGCTTCTTCTAGACCTG<br>GTGGTGGTCCTGATGGTTCTGGTATTAGATCTCAAGATCCTTGGAGACCTTTTAAGAAGCTTGA<br>TATTGAAAGAGATATTCAACCTTGTGCTACTGGTGTTTGTAATTGTCTTTCTTGA |
| Pc13g04<br>050<br>(native) | MASIQVWMLRIQGANIRDHNA<br>MVDPSVSGINDIPKDQPTLVR<br>WYMDVRRWDEKYFDLPLLET<br>LTQPDQAAVKKYYQTSDKRLS<br>LASQLLKYYYIHQATGTPWSKI<br>EIQRTPMPENRPFYDSSLDFNV<br>SHQAGLTLFAGTRAATAHSLSG<br>GPQTLPRVGIDVACVDEPSRRR<br>ANRPPKTLADLATFVDVFSDV | ATGGCTTCTATCCAGGTGTGGATGCTCCGCATCCAAGGTGCGAATATACGAGATCACAATGCAA<br>TGGTAGACCCCAGTGTGTCTGGAATTAATGACATCCCCAAGGACCAGCCCACGTTGGTCCGAT<br>GGTACATGGATGTCAGACGTTGGGATGAAAAATACTTTGATCTCCCTTTGCTTGAAACCTTAAC<br>ACAGCCTGATCAGGCAGCTGTCAAGAAGTACTATCAAACATCGGACAAGCGCCTGTCCTTGGC<br>CTCCCAGTTGCTGAAATATTACTACATTCACCAAGCCACTGGCACTCCCTGGAGCAAGATTGAG<br>ATCCAGCGTACTCCGATGCCCGAAAATCGACCATTCTACGATTCAAGCCTGGATTTCAACGTCA<br>GCCATCAGGCTGGTCTCACTCTGTTCGCAGGCACGCGTGCCGCAACAGCCCACTCCTTATCCG<br>GTGGACCTCAAACATTGCCTCGCGTGGGAATTGACGTTGCGTGTGTTGATGAACCCTCTCGTC<br>GTCGTGCTAATCGTCCCCCGAAGACACTTGCCGACCTTGCAACCTTCGTGGATGTCTTCAGTGA |

| | | |
|---|---|---|
| | LSLRELATIKNPYATLKLAREL<br>GLNKSDPSKDDQEVLAAYGIR<br>LFYSIWALKEAYLKMTGDGLL<br>ASWIKDLEFTNVVPPEPVQTV<br>GFAGDPSATHAPSVQNWGRPY<br>SDVKISLRGIPDHSVRVQLVGF<br>ESDYIVATAASGPNIGSVSRQV<br>VVNDSDHHLPGRITAFDSETGL<br>QNVRIPPIALRSIGDGDPWRVD<br>SKISDPWLPMQEVDIEIDIRPCA<br>DGRCEHLRDLPSF | CGTTCTCTCACTCCGTGAGCTTGCGACCATCAAGAACCCGTACGCGACTCTTAAATTGGCTCGT<br>GAGCTTGGTCTGAATAAAAGTGACCCGAGCAAAGACGACCAGGAAGTCCTTGCTGCCTACGG<br>CATTCGGCTGTTCTACTCGATTTGGGCTCTCAAGGAGGCTTACTTGAAAATGACCGGAGACGG<br>CCTTCTGGCCTCTTGGATAAAGGATCTGGAATTCACAAACGTTGTTCCCCCCGAACCAGTTCAA<br>ACAGTCGGATTTGCTGGTGATCCTTCTGCCACTCACGCGCCCTCGGTCCAAAATTGGGGCCGG<br>CCTTACTCCGATGTCAAAATCTCCTTGCGTGGCATTCCTGACCATTCTGTGCGCGTTCAGCTCGT<br>CGGCTTCGAGTCCGACTACATAGTTGCCACGGCCGCGTCGGGCCCCAATATTGGATCCGTTTCG<br>CGGCAGGTAGTCGTGAATGACAGCGATCACCATCTGCCAGGGCGTATCACAGCCTTCGACTCT<br>GAGACTGGACTCCAGAACGTCCGCATTCCCCCAATCGCGCTTCGATCAATTGGCGATGGGGAC<br>CCCTGGCGTGTGGACTCGAAAATCAGCGACCCCTGGCTCCCCATGCAGGAGGTCGATATTGAA<br>ATCGATATCCGGCCCTGTGCGGATGGTCGTTGCGAGCACCTACGGGATTTACCAAGCTTTTAA |
| PaaT | MVLVTCTMEAPRSDQAHTDAT<br>TPMEAIRTTSLGTNNYGPVPDD<br>YLDLPVREVNDGADLREYITE<br>TRTGEIIKPIKSNVTGKTEDWK<br>MVTFTIDDPENPKNWSKAFK<br>WYCTMVVAFTCFVVAFCSSVI<br>TADVEGPIEEFGIGREASLVVIT<br>VFVIGFGLGPMVFAPMSEIVGR<br>RPVYALTLALAVIFVIPCAVSKN<br>IGTLIVCRLIDGIAFSAPMTLVG<br>GTLADLWKSEERGVPMAAFSA<br>APFIGPAIGPLVGGYLADNCGW<br>RWLYWIQLILAFVAWVMITFT<br>VPETFAPILLKKRAQKLRKAED<br>DPKYTTETELDARPMGEKLRIF<br>LFRPFQLLFLEPIVLFISLYMSVI<br>YGLLYMFFVAYPIVYMGGKG<br>WSASNTGLMFIPLAIGVIFSAC<br>CAPFVNNHYLKVSVAYGGKPP<br>AEKRLIPMMWACWCIPSGLFV<br>FAWTSYPDLHWMGPAMGGFLI<br>GVGVILLYNSANNYLVDTYQH<br>QAASALAAKTFIRSIWGACTV<br>LFTEQMYERLGDQWASTLLAF<br>IGLACCAIPYVFYFKGESIRRFS<br>KFAFSDDEEKAIKA | ATGGAGGCTCCACGGTCAGATCAGGCTCATACAGATGCTACCACGCCAATGGAGGCTATACGG<br>ACAACTTCTCTCGGGACTAACAACTATGGACCAGTACCTGATGATTACCTCGACCTACCAGTAC<br>GCGAAGTAAATGATGGCGCCGATTTACGAGAATACATAACCGAGACGAGGACAGGTGAAATCA<br>TCAAGCCGATCAAGTCCAATGTTACTGGGAAAACAGAAGATTGGAAAATGGTGACTTTCACAA<br>TAGACGATCCTGAGAATCCAAAGAACTGGTCGAAGGCTTTTAAGTGGTATTGTACCATGGTGGT<br>TGCATTCACCTGCTTTGTTGTTGCCTTTTGCAGTAGTGTCATTACAGCTGATGTTGAGGGACCTA<br>TTGAGGAATTTGGTATCGGTAGGGAAGCATCATTGGTTGTTATAACGGTTTTCGTTATTGGATTT<br>GGTCTAGGACCCATGGTCTTTGCTCCGATGTCAGAGATCGTAGGGAGACGACCTGTCTACGCTC<br>TGACACTTGCATTGGCTGTTATTTTCGTAATTCCTTGTGCGGTTTCAAAGAACATTGGAACTTTA<br>ATTGTATGCAGGTTGATAGACGGAATCGCTTTTAGTGCTCCAATGACTTTGGTCGGTGGTACTCT<br>AGCTGATTTGTGGAAAAGTGAAGAGCGTGGGGTTCCGATGGCAGCATTCAGCGCTGCTCCATT<br>CATTGGTCCTGCTATTGGCCCACTTGTTGGTGGCTACCTGGCTGACAATTGTGGTTGGAGGTGG<br>CTATATTGGATACAACTTATTTGGCCTTTGTGGCCTGGGTGATGATCACATTCACTGTGCCCGA<br>AACTTTTGCGCCAATTTTGTTGAAAAAGCGAGCACAGAAACTTAGAAAGGCTGAGGATGACCC<br>TAAGTATACAACTGAAACCGAACTCGATGCTAGACCCATGGGAGAGAAGCTGAGAATCTTTTT<br>ATTTCGCCCTTTCCAACTTCTGTTCCTCGAGCCCATTGTATTATTCATTTCTCTATACATGTCAGT<br>GATCTATGGGTTGCTTTACATGTTTTTCGTTGCCTATCCGATTGTTTATATGGGCGGCAAAGGATG<br>GAGCGCGTCTAATACAGGTCTCATGTTCATTCCTCTGGCGATTGGTGTGATATTCAGCGCATGTT<br>GTGCTCCTTTTGTCAACAATCACTACCTCAAGGTCTCTGTGGCATACGGTGGCAAGCCTCCAGC<br>TGAAAAAAGATTGATTCCAATGATGTGGGCTTGTTGGTGCATACCTTCTGGGCTTTTTGTGTTC<br>GCTTGGACATCCTATCCCGATCTTCACTGGATGGGACCAGCGATGGGAGGGTTTCTTATTGGAG<br>TCGGCGTTATTTTATTGTATAATTCTGCAAATAATTACCTTGTTGATACCTATCAGCATCAAGCAG<br>CATCTGCTTTAGCAGCCAAAACCTTTATCAGGTCTATATGGGGTGCTTGCACTGTTCTGTTCACT<br>GAACAAATGTACGAACGTCTTGGTGACCAGTGGGCCAGTACGCTTCTTGCATTTATCGGATTGG<br>CATGTTGCGCCATTCCTTATGTGTTTTATTTCAAGGGAGAGTCCATTAGACGTTTTTCGAAATTT<br>GCCTTTTCAGACGATGAGGAAAAAGCAATAAAGGCATGA |
| PenM | MKDGEETPSVDGSTSASNREK<br>LGTDLEIGPVDLSDGGKEEKV<br>KDPNLVDWDGPDDPENPLNW | ATGAAGGATGGTGAGGAGACACCTTCAGTTGATGGTTCAACATCAGCTTCTAATAGAGAGAAA<br>CTTGGAACAGATTTGGAAATTGGACCAGTTGATCTTTCTGATGGAGGAAAAGAAGAAAAAGTT<br>AAGGATCCAAATCTCGTTGATTGGGATGGACCAGATGATCCTGAGAATCCTCTTAATTGGACTT |

| | | |
|---|---|---|
| | TSKRKITATCSIALITFLTPLGSS<br>MFAPGVGQLVKDFNVTSTELS<br>SFVVSVYLLGYCFGPLIIAPLSE<br>LYGRQYVYHVCNILYVIWTIA<br>CAFAPEIGSLVVFRFFAGLAGS<br>CPLTIGAGSIADMFVQEQRGG<br>AMAAWALGPLIGPVVGPVAGA<br>YLAQAKGWRWSFYVLAMAA<br>GAITISSLFSIRESYAPTLLARKT<br>KKLQKETGNMNLRSALDTGRT<br>PKELFLYSIVRPTKMLFRSPIVF<br>LLSLYVGVIYGYLYLLFTTITSV<br>FQQQYNFSQGAVGLTYLGLGV<br>GSLIGLFLIGATSDRLLNYLAA<br>KNGEKKPEYRLPPMVPGAIFV<br>PISLFMYGWTAYYQTHWIVPII<br>GTSFLGTGMMITFMCVSTYLV<br>DAFTNYAASVMAANTVFRSLA<br>GALLPLAGPKMYAVLGLGWG<br>NSLLGFIALAFCALPVIFWIYG<br>ERIRTSPKFQVTF | CAAAGAGAAAAATTACCGCTACATGTTCAATTGCTTTGATTACTTTTCTCACTCCTTTGGGTTCA<br>TCAATGTTTGCTCCAGGAGTTGGACAATTGGTTAAAGATTTTAATGTGACTTCCACTGAGCTTT<br>CATCATTTGTTGTTTCTGTTTATCTCCTTGGTTATTGTTTTGGTCCACTTATTATTGCTCCACTTTC<br>AGAACTTTATGGAAGACAATATGTTTACCATGTTTGTAATATCCTCTACGTTATTTGGACTATTGC<br>TTGTGCTTTTGCTCCTGAGATTGGATCTCTTGTTGTTTTTAGATTTTTCGCTGGACTTGCTGGTTC<br>ATGTCCTTTGACAATTGGAGCTGGTTCTATTGCTGATATGTTTGTTCAAGAGCAAAGAGGTGGT<br>GCTATGGCTGCTTGGGCTCTTGGACCATTGATTGGTCCAGTTGTTGGTCCTGTTGCTGGTGCTTA<br>TCTTGCTCAAGCTAAGGGTTGGAGATGGTCTTTTTATGTTCTTGCTATGGCTGCAGGTGCTATTA<br>CTATTTCTTCTCTTTTTAGTATCAGGGAATCATATGCTCCAACTCTTTTGGCTAGAAAGACTAAG<br>AAATTGCAAAAGGAAACAGGTAATATGAACTTGAGATCAGCTCTTGATACTGGTAGAACACCT<br>AAAGAGTTGTTTTTGTATTCAATCGTTAGGCCAACTAAGATGCTTTTTAGATCACCAATTGTTTT<br>CCTTTTGTCACTTTATGTTGGAGTTATATATGGTTACTTGTACCTTTTGTTCACAACTATTACTTCA<br>GTTTTCCAACAACAATACAACTTTTCACAAGGAGCTGTTGGACTTACATATCTTGGACTTGGAG<br>TTGGTTCTTTGATTGGATTGTTTTTGATTGGAGCTACTTCAGATAGACTTTTGAATTATCTTGCTG<br>CTAAGAATGGAGAAAAGAAACCTGAGTATAGACTTCCACCAATGGTTCCAGGAGCTATTTTTGT<br>TCCTATTTCATTGTTTATGTACGGATGGACTGCTTATTATCAAACACATTGGATTGTTCCAATCAT<br>TGGAACATCTTTTCTTGGTACTGGTATGATGATTACTTTTATGTGTGTTTCAACCTACCTTGTTGA<br>TGCTTTTACTAATTATGCTGCTTCAGTTATGGCTGCTAATACAGTTTTTAGATCATTGGCTGGAGC<br>TCTTCTTCCTCTTGCTGGACCAAAGATGTATGCTGTTTTGGGTCTTGGTTGGGGAAATTCTTTGC<br>TTGGTTTTATTGCTCTTGCTTTTTGTGCTTTGCCAGTTATTTTCTGGATATATGGAGAAAGAATCA<br>GAACTTCACCTAAATTTCAAGTTACTTTCTAA |
| PCL<br>(native) | MVFLPPKESGQLDPIPDNIPISE<br>FMLNERYGRVRHASSRDPYTC<br>GITGKSYSSKEVANRVDSLARS<br>LSKEFGWAPNEGSEWDKTLAV<br>FALNTIDSLPLFWAVHRLGGVL<br>TPANASYSAAELTHQLLDSKA<br>KALVTCVPLLSISLEAAAKAGL<br>PKNRIYLLDVPEQLLGGVKPPA<br>GYKSVSELTQAGKSLPPVDEL<br>RWSAGEGARRTAFVCYSSGTS<br>GLPKGVMISHRNVIANTLQIKA<br>FEQNYRDGGGTKPASTEVALG<br>LLPQSHIYALVVIGHAGAYRGD<br>QTIVLPKFELKSYLNAIQQYKI<br>SALFLVPPIIIHMLGTQDVCSKY<br>DLSSVTSLFTGAAPLGMETAA<br>DFLKLYPNILIRQGYGLTETCT<br>VVSSTHPHDIWLGSSGALLPG<br>VEARIVTPENKEITTYDSPGEL | ATGGTTTTTTTACCTCCAAAGGAGTCCGGTCAATTGGACCCAATTCCCGACAATATTCCAATCA<br>GCGAGTTTATGCTCAATGAGAGATATGGACGAGTGCGACACGCCAGCTCCCGGGACCCATACA<br>CCTGTGGTATTACCGGGAAGTCATACTCGTCGAAAGAGGTAGCCAATCGCGTCGACTCGCTGG<br>CTCGTAGTCTATCAAAGGAATTTGGTTGGGCGCCGAATGAAGGGTCAGAATGGGATAAGACAT<br>TGGCCGTGTTTGCCCTCAACACTATCGATTCCTTACCCCTATTCTGGGCCGTTCACAGACTGGG<br>CGGTGTTCTCACTCCCGCCAACGCATCATACTCCGCCGCCGAGCTGACGCATCAGCTGCTTGAT<br>TCCAAGGCCAAGGCCCTTGTGACTTGTGTTCCTCTCCTCTCCATCTCACTGGAAGCTGCAGCCA<br>AAGCTGGTCTCCCGAAGAACAGAATCTACTTACTCGATGTACCTGAGCAGCTTCTTGGCGGAG<br>TCAAGCCTCCAGCAGGATACAAGTCCGTTTCCGAACTGACCCAGGCTGGGAAGTCTCTCCCGC<br>CAGTGGATGAATTGCGATGGAGCGCGGGTGAAGGTGCCCGGCGAACAGCATTTGTGTGCTACT<br>CAAGTGGAACGTCTGGATTGCCGAAAGGAGTCATGATCTCACACCGCAACGTGATCGCCAATA<br>CCCTTCAGATCAAGGCGTTTGAGCAGAACTACCGGGATGGTGGGGGCACAAAGCCTGCGAGT<br>ACTGAGGTTGCTCTTGGTCTCCTTCCGCAGAGCCATATCTATGCTCTTGTGGTCATTGGCCATGC<br>TGGGGCATACCGAGGCGACCAAACAATCGTTCTCCCCAAATTCGAATTGAAATCCTACCTGAA<br>CGCCATCCAACAGTACAAGATCAGTGCGCTGTTCCTGGTACCTCCGATCATCATTCACATGCTG<br>GGCACTCAAGACGTGTGCTCCAAGTATGACCTGAGTTCCGTGACGTCTCTGTTCACGGGAGCG<br>GCACCCCTGGGTATGGAGACAGCTGCCGATTTCCTCAAACTCTACCCGAACATTTTGATCCGCC<br>AAGGATACGGTCTGACAGAGACATGCACGGTCGTAAGCTCGACCCACCCGCACGATATCTGGC<br>TAGGTTCATCCGGCGCTTTGCTCCCTGGAGTCGAGGCACGAATTGTGACGCCTGAAAACAAGG |

| | | |
|---|---|---|
| | VVRSPSVVLGYLNNEKATAETFVDGWMRTGDEAVIRRSPKGIEHVFIVDRIKELIKVKGLQVAPAELEAHILAHPDVSDCAVIAIPDDRAGEVPKAIVVKSASAGSDESVSQALVKYVEDHKARHKWLKGGIRFVDAIPKSPSGKILRRLIRDQEKEARRKAGSKI | AAATCACAACGTACGACTCACCGGGCGAATTGGTGGTCCGAAGCCCAAGCGTCGTCCTGGGCTATTTGAACAACGAAAAAGCCACCGCAGAGACATTTGTGGACGGATGGATGCGTACGGGAGACGAGGCTGTCATCCGTAGAAGCCCGAAGGGCATCGAGCACGTGTTTATTGTCGATCGGATCAAGGAGTTGATCAAGGTCAAGGGTCTGCAAGTCGCGCCTGCCGAACTCGAAGCCCATATCCTCGCCCACCCCGATGTCTCGGACTGTGCTGTCATCGCTATTCCGGATGATCGTGCAGGAGAAGTACCCAAGGCCATTGTTGTGAAGTCCGCCAGCGCAGGATCGGACGAATCTGTCTCCCAGGCTCTCGTGAAGTATGTTGAGGACCACAAGGCTCGTCACAAGTGGTTGAAGGGAGGTATCAGATTTGTGGATGCCATTCCCAAGAGCCCGAGTGGTAAGATTCTTCGTCGGTTGATCCGTGACCAAGAGAAGGAGGCACGGAGAAAGGCTGGTAGCAAGATCTAA |

Gene and protein sequences used in this study.

**Table S5.**

| Name | Sequence |
| --- | --- |
| pcbAB native-F1 | ttaacagctatacaagtttgtacaaaaaactgATGACTCAACTGAAGCCACCGAACGGAACCACGCCGATAGGCTTCTCGGCCACTACATCCCTGAACGCCA<br>GTGGGAGCTCGAGTGTGAAAAATGGACCATCAAACCCAGCAATGGCATCTTCAAGCCCAGCACTAGGGACACCATGGACCCTTGCAG<br>TGGGAATGCGGCCGATGGCAGTATCCGCGTCCGTTTCCGTGGAGGAATCGAACGGTGGAAGGAGTGCGTCAACCAGGTCCCCGAGCGC<br>TGCGACCTGAGTGGTCTGACAACCGACTCCACGCGATATCAGCTCGCATCGACTGGGTTCGGTGACGCGAGCGCTGCCTACCAGGAGCG<br>CTTGATGACGGTCCCTGTTGACGTACATGCCGCGCTCCAAGAGCTGTGCCTAGAACGCCGTGTGAGCGTGGGATCCGTCATTAATTTCTC<br>CGTGCACCAGATGCTGAAAGGGTTTGGAAATGGCACACACACTATCACCGCCTCTCTGCACCGTGAGCAGAATTTGCAGAATTCTTCGC<br>CATCCTGGGTAGTCTCCCCCACAATCGTCACCCATGAGAACAGAGACGGATGGTCCGTCGCGCAGGCGGTCGAGAGTATCGAAGCGGG<br>GCGCGGTTCCGAGAAGGAGTCAGTGACTGCGATTGACTCCGGGTCAAGTCTCGTGAAAATGGGGTTATTTGACTTACTCGTCAGCTTTG<br>TCGATGCAGACGATGCTCGTATTCCATGTTTCGACTTTCCCCTCGCAGTGATAGTGCGTGAGTGTGATGCCAACCTCTCGCTGACTCTGC<br>GTTTCTCCGACTGTCTCTTCAACGAGGAGACGATATGCAATTTTACCGATGCCCTAAACATCTTGCTCGCCGAAGCAGTGATAGGAAGA<br>GTGACCCCGGTTGCCGATATCGAACTACTATCCGCGGAGCAGAAGCAGCAGCTGGAAGAGTGGAACAACACGGATGGCGAGTACCCTT<br>CATCAAAGCGACTGCACCATCTCATTGAAGAGGTGGTTGAACGGCATGAAGACAAAATAGCCGTTGTCTGCGACGAGCGAGAGCTCAC<br>TTACGGCGAGCTCAATGCCCAAGGCAACAGCCTCGCACGCTATCTCCGTTCCATTGGTATCCTGCCCGAGCAGCTAGTCGCATTGTTTCT<br>AGATAAGAGCGAGAAGCTCATTGTTACCATCCTCGGCGTGTGGAAATCCGGCGCCGCCTACGTGCCCATCGACCCGACTTATCCGGATG<br>AGCGAGTGCGCTTCGTGCTGGATGACACCAAGGCACGGGCCATCATCGCCAGTAATCAACATGTGGAGAGGCTCCAGCGAGAGGTCAT<br>CGGCGATAGAAACCTATGCATTATCCGTCTGGAGCCCTTGTTGGCCTCCCTTGCTCAGGATTCCTCAAAATTCCCCGCGCATAACTTGGA<br>CGACCTACCCCTCACAAGCCAGCAGCTCGCCTATGTGACTTACACCTCTGGGACCACTGGCTTCCCAAAGGGCATATTTAAACAACACA<br>CCAATGTGGTGAACAGTATCACCGACCTGTCTGCAAGGTACGGGGTGGCCGGGCAGCACCACGAAGCCATTCTGCTTTTCTCGGCCTGC<br>GTGTTCGAGCCGTTCGTTCGACAGACGCTCATGGCACTCGTGAATGGCCATCTCCTCGCAGTTATCAATGACGTGGAAAAATATGATGC<br>CGATACGCTCCTGCCGTTCATACGCAGACACAGCATCACCTACCTCAATGGTACTGCCTCTGTCCTGCAAGAGTACGACTTTTCCGACTG<br>CCCATCACTGAATCGGATAATCCTGGTGGGTGAGAACCTGACAGAAGCCCGGTATCTGGCGCTGCGCCAGCGGTTCAAGAATCGCATCC<br>TCAACGAGTATGGTTTTACCGAGTCAGCCTTTGTAACGGCCCTCAAGATTTTCGACCCGGAGTCGACCCGTAAGGACACGAGTCTGGGG<br>AGACCGGTGCGCAACGTCAAGTGCTACATCCTCAATCCATCCCTTAAACGTGTCCCGATTGGAGCTACGGGTGAGTTGCATATCGGAGG<br>GTTGGGCATTTCCAAGGGATACCTCAACCGCCCCGAACTCACGCCGCACCGCTTCATTCCCAACCCCTTCCAAACGGATTGCGAGAAGC<br>AGCTCGGGATCAACAGCTTGATGTACAAGACCGGTGACCTGGCCCGCTGGCTTCCGAACGGCGAGGTTGAGTATCTCGGACGCGCAGAT<br>TTCCAGATCAAACTGCGAGGTATTCGAATTGAACCTGGTGAAATTGAGACGATGCTGGCTATGTACCCTAGGGTCCGGACCAGTTTAGT<br>GGTGTCCAAAAAGCTCCGCAACGGTCCAGAGGAAACTACCAACGAGCACCTCGTGGGTTATTATGTTTGTGATAGCGCCTCAGTGTCCG<br>AGGCAGACCTGCTGTCATTTTTAGAGAAGAAACTGCCTCGATACATGATTCCCACGCGGTTGGTACAGCTGTCGCAGATCCCAGTGAAT<br>GTGAACGGGAAGGCGGACCTACGCGCCTTGCCGGCCGTCGATATCTCCAATTCCACGGAGGTCCGTTCCGACCTTCGAGGCGATACGGA<br>AATCGCCCTCGGGGAAATCTGGGCCGACGTGTTGGGAGCCCGCCAGAGATCCGTCTCTCGCAACGACAACTTCTTCCGCCTAGGAGGGC<br>ACAGCATCACCTGCATCCAACTGATCGCTCGCATCCGACAACGACTCTCGGTCAGCATCTCCGTCGAAGATGTTTTTGCAACAAGGACA<br>CTTGAGCGCATGGCAGACCTTCTACAGAACAAGCAGCAGGAGAAATGCGACAAACCCCATGAGGCGCCGACAGAGCTGCTTGAGGAGA<br>ATGCAGCAACGGACAATATCTATCTGGCAAACAGTCTTCAGCAGGGCTTCGTCTACCATTACCTCAAGAGCATGGAACAATCCGACGCC<br>TATGTAATGCAGTCCGTTCTTCGGTACAACACCACATTGTCTCCAGATCTGTTTCAGAGAGCCTGGAAGCATGCACAGCAGTCCTTTCCA |

| | |
|---|---|
| | GCGCTGCGGCTGCGGTTCTCATGGGAAAAGGAGGTTTTCCAACTGCTCGATCAGGATCCACCATTGGACTGGCGTTTCCTCTACTTCACC<br>GACGTTGCCGCGGGTGCTGTCGAGGACCGGAAATTGGAAGACTTGCGGCGCCAAGACCTTACGGAGAGATTCAAGCTGGATGTTGGCA<br>GACTGTTCCGCGTCTATCTGATTAAACACAGCGAGAATCGCTTCACGTGTCTTTTCAGCTGCCACCATGCAATCCTCGATGGTTGGAGTC<br>TGCCACTCTTGTTCGAAAAGGTTCACGAGACCTACCTGCAACTGCTGCATGGGGACAATCTCACTTCGTCCATGGATGACCCTTACACTC<br>GCACCCAGCGGTATCTCCACGCTCACCGTGAGGATCACCTCGACTTTTGGGCCGGTGTGGTTCAAAAGATCAACGAACGGTGTGATATG<br>AACGCCTTGTTGAACGAGCGCAGTCGTTACAAAGTCCAGCTGGCAGACTATGACCAGGTGCAGGAGCAGCGACAGCTGACAATTGCTC<br>TCTCTGGAGACGCATGGCTAGCAGACCTTCGTCAGACCTGCTCCGCCCAGGGTATTACCTTACATTCGATTCTCCAATTTGTTTGGCACG<br>CCGTGCTGCACGCTTATGGCGGTGGCACCCACACCATAACCGGCACGACCATTTCTGGAAGGAACCTGCCCATCTTGGGAATTGAACGA<br>GCAGTTGGTCCGTATATCAACACTCTACCGCTGGTACTCGATCATTCG |
| pcbAB native-F2 | CAACACTCTACCGCTGGTACTCGATCATTCGACGTTCAAGGATAAGACAATCATGGAGGCCATCGAGGATGTGCAGGCCAAGGTAAAC<br>GTCATGAACAGCCGGGGCAATGTGGAACTGGGCCGTTTGCACAAAACCGACTTAAAGCACGGATTATTCGATTCTTTATTCGTGCTTGA<br>AAACTACCCGAATTTGGACAAATCGCGAACACTTGAGCACCAGACTGAACTGGGGTATTCGATTGAAGGCGGCACTGAGAAGCTGAAT<br>TATCCACTGGCTGTCATCGCGCGCGAAGTCGAGACGACTGGCGGATTCACAGTATCCATCTGCTACGCCAGTGAGCTATTTGAGGAGGT<br>TATGATCTCCGAGCTTCTTCATATGGTCCAGGACACACTGATGCAGGTTGCCCGAGGTTTGAATGAACCCGTCGGCAGCCTGGAGTATCT<br>CTCATCTATCCAATTGGAGCAACTCGCCGCGTGGAATGCCACGGAAGCTGAGTTTCCCGATACCACGCTTCATGAGATGTTTGAAAACG<br>AAGCGAGCCAGAAGCCGGACAAGATAGCAGTGGTCTATGAGGAGACGTCCTTGACTTACCGCGAGTTGAATGAGCGGGCGAACCGTAT<br>GGCACATCAGCTAAGGTCCGACGTCAGCCCCAACCCCAACGAGGTCATTGCGCTGGTGATGGACAAGAGCGAGCATATGATCGTCAAC<br>ATTCTGGCCGTATGGAAGAGCGGCGGTGCCTATGTCCCCATTGACCCTGGATATCCTAACGACCGCATTCAATACATCCTAGAGGACAC<br>ACAAGCCCTCGCAGTCATCGCGGACTCCTGCTATCTGCCTCGCATCAAGGGAATGGCTGCCTCCGGCACGCTTCTTTATCCCTCTGTCTT<br>GCCTGCCAATCCGGATTCCAAGTGGAGCGTATCGAACCCTTCACCGTTGAGTCGGAGCACGGACTTAGCTTATATCATCTATACCTCTGG<br>AACGACAGGTCGGCCCAAGGGCGTCACGGTAGAGCATCATGGAGTGGTCAACCTGCAGGTGTCGCTATCCAAAGTATTCGGACTACGG<br>GATACTGACGACGAGGTAATTCTCTCCTTTTCCAACTATGTGTTCGACCATTTCGTGGAGCAGATGACCGACGCCATTCTCAATGGCCAA<br>ACCCTCCTGGTCCTCAACGATGGAATGCGCGGGGACAAAGAGCGACTCTACAGATACATTGAGAAGAACCGAGTGACCTACTTGTCTG<br>GCACCCCATCCGTGGTCTCCATGTACGAATTTAGCCGGTTCAAGGACCATCTACGCCGTGTGGACTGCGTGGGGGAGGCGTTCAGCGAA<br>CCGGTCTTCGACAAGATCCGCGAAACGTTCCATGGCCTCGTTATCAACGGCTACGGCCCAACTGAAGTTTCCATCACCACCCACAAGCG<br>GCTCTATCCATTCCCAGAGCGGCGAATGGACAAAAGTATTGGCCAACAGGTCCACAATAGCACGAGCTATGTGCTGAACGAGGACATG<br>AAGCGCACCCCCATAGGGGCTGTCGGCGAGCTCTACCTGGGTGGTGAAGGAGTGGTACGGGGATATCACAATCGCGCAGATGTGACCG<br>CGGAGCGTTTTATTCCTAATCCATTCCAGTCGGAAGAAGATAAGCGAGAAGGTCGTAACTCCCGTTTGTACAAGACCGGTGACCTGGTA<br>CGCTGGATTCCTGGAAGCAGCGGGGAGGTCGAGTATCTAGGTCGTAATGACTTCCAGGTCAAGATTCGCGGACTGCGCATCGAACTAGG<br>CGAGATTGAGGCCATCCTATCGTCTTATCACGGAATCAAACAGTCTGTGGTGATTGCCAAGGATTGCAGAGAAGGGGCCCAGAAATTCC<br>TGGTTGGTTACTATGTCGCCGATGCAGCGCTGCCGTCCGCTGCCATTCGGCGCTTCATGCAGTCTCGGCTCCCTGGCTACATGGTGCCCT<br>CTCGTCTCATTCTCGTCAGCAAGTTCCCCGTCACTCCTAGTGGAAAATTAGACACCAAGGCTTTGCCCCCAGCCGAGGAAGAGAGCGAG<br>ATTGACGTGGTGCCGCCGCGTAGTGAAATCGAACGCTCCTTGTGTGACATCTGGGCGGAACTACTCGAGATGCACCCAGAGGAGATCGG<br>CATTTACAGCGATTTCTTCAGCCTGGGAGGTGACAGCCTAAAGAGCACAAAGCTTTCCTTCATGATTCACGAGTCCTTTAACCGCGCCGT<br>CTCAGTCAGCGCCCTTTTCTGTCACCGGACAGTTGAAGCCCAGACGCACTTGATCCTGAACGATGCTGCAGATGTGCACGAAATTACTC<br>CCATAGATTGCAATGATACGCAGATGATTCCCGTGTCCCGTGCCCAGGAGCGACTCCTCTTCATCCACGAATTTGAGAATGGCAGCAAT<br>GCATACAATATCGACGCTGCATTTGAACTGCCTGGCTCGGTTGACGCGTCGCTTCTCGAGCAGGCGCTGCGTGGAAACCTTGCTCGACA<br>TGAGGCGTTGAGAACTTTACTGGTCAAGGATCACGCAACCGGCATCTATCTTCAGAAGGTATTGAGTCCCGATGAAGCCCAGGGCATGT<br>TCTCCGTCAACGTGGACACAGCCAAGCAGGTGGAGCGGCTGGACCAGGAGATAGCCAGTCTATCCCAGCATGTTTTCCGCCTCGATGAT<br>GAACTGCCTTGGGAGGCCCGCATCCTTAAACTCGAATCCGGCGGCCTGTATCTCATTCTGGCGTTCCACCATACCTGCTTCGATGCATGG |

| | |
|---|---|
| | TCATTGAAAGTCTTCGAGCAAGAGCTTCGGGCCTTGTACGCAGCGCTCCAGAAAACCAAAAGTGCAGCGAACTTACCAGCCCTCAAAGC<br>GCAGTACAAGGAATACGCGCTCTACCATCGCCGGCAGCTGTCTGGCGATCGCATGCGCAACCTGTCAGACTTTTGGCTGCGGAAACTCA<br>TTGGCTTGGAACCATTGCAGCTGATCACGGACCGCCCACGTCCTGTGCAATTCAAATACGACGGTGACGACCTCAGTATCGAACTGAGC<br>AAGAAGGAAACGGAGAACCTGAGGGGGGTGGCCAAACGTTGCAAGTCGAGTCTGTACGTCGTGTTGGTTTCCGTTTATTGCGTTATGCT<br>AGCCTCGTACGCGAACCAGTCCGATGTTTCCGTGGGTATCCCAGTCAGCCACCGAACGCATCCTCAGTTCCAATCGGTCATTGGATTCTT<br>CGTCAACCTTGTGGTGCTAAGGGTGGATATTTCTCAGTCAGCCATTTGCGGGCTCATCAGAAGGGTAATGAAAGAGCTCGTGGACGCCC<br>AACTGCACCAAGACATGCCGTTCCAGGAAGTGACGAAGCTGCTGCAGGTGGATAATGACCCCAGCCGGCATCCGCTGGTACAGAACGT<br>GTTCAACTTCGAATCCCGTGCGAACGGAGAACACGATGCCAGGTCGGAGGATGAAGGATCGCTTGCATTCAATCAATACCGGCCGGTTC<br>AGCCCGTGGATTCCGTTGCGAAGTTCGATCTGAACGCAACGGTCACGGAATTGGAGTCGGGATTGAGAGTCAACTTCAACTATGCGACC<br>AGCCTATTCAACAAAAGCACGATCCAGGGTTTTTGCATACCTATGAGTATCTCCTGCGCCAGCTGTCCGAACTGAGTGCAGAAGGGAT<br>CAATGAGGATACGCAGCTGTCGTTAGTTCGCCCGACAGAGAATGGCGATCTGCACTTGCCATTGGCACAGTCCCCGCTTGCGACGACTG<br>CTGAGGAGCAGAAAGTAGCGTCGTTGAACCAGGCCTTTGAGCGCGAAGCTTTCCTTGCCGCAGAGAAGATTGCCGTCGTGCAGGGAGA<br>TAGAGCACTTAGTTATGCTGATCTTAACGGGCAGGCTAACCAGCTCGCCCGGTACATACAGTCCGTGTCCTGTATTGGGGCAGACGACG |
| pcbAB native-F3 | CATACAGTCCGTGTCCTGTATTGGGGCAGACGACGGAATAGCTTTGATGCTGGAAAAGAGTATCGACACGATTATTTGCATTCTCGCGA<br>TTTGGAAGGCTGGTGCAGCATACGTGCCCTTGGATCCGACTTACCCACCCGGACGCGTCCAGCTGATTCTGGAGGAGATTAAAGCGAAG<br>GCTGTCCTTGTGCACTCCAGTCATGCTTCGAAATGTGAACGCCATGGCGCGAAGGTGATTGCAGTCGACTCGCCCGCCATCGAGACGGC<br>GGTCAGCCAACAGTCAGCTGCTGACCTGCCCACAATTGCTAGCCTCGGCAATCTAGCGTATATAATCTTTACTTCAGGCACTTCCGGTAA<br>GCCAAAGGGAGTCCTAGTTGAGCAAAAGGCAGTTCTTCTTCTACGCGATGCCCTCCGGGAGCGGTATTTCGGTCGAGACTGTACCAAGC<br>ATCATGGCGTCCTGTTCCTGTCCAACTACGTCTTCGACTTCTCCGTCGAACAACTTGTGTTGTCGGTGCTCAGCGGACACAAGCTGATCG<br>TTCCCCCAGCTGAGTTCGTCGCAGATGATGAATTTTACAGAATGGCCAGCACGCACGGTCTCTCCTATCTCAGCGGCACACCATCCTTAC<br>TGCAGAAGATCGATCTGGCACGACTGGACCATCTGCAGGTTGTTACCGCCGCGGGCGAAGAGCTTCACGCCACCCAGTACGAGAAGAT<br>GCGCCGCCGATTCAACGGTCCCATCTACAATGCCTATGGTGTCACCGAGACCACGGTGTACAACATTATCGCGGAATTCACAACGAATT<br>CGATATTTGAGAATGCTCTTCGGGAAGTGCTCCCTGGTACCCGAGCGTATGTGCTGAACGCGGCACTTCAGCCCGTCCCCTTCGATGCTG<br>TCGGAGAACTCTATCTTGCCGGCGACAGCGTTACGCGTGGTTATCTCAACCAACCTCTTCTAACGGATCAGCGATTCATTCCCAACCCTT<br>TCTGCAAAGAGGAGGACATCGCTATGGGGCGCTTCGCGCGGCTCTACAAGACCGGCGACCTGGTTCGATCGCGTTTCAACCGTCAGCAG<br>CAGCCGCAGCTGGAATACCTAGGAAGAGGCGATCTGCAGATCAAGATGAGGGGATACCGGATCGAGATTTCTGAAGTTCAGAACGTGC<br>TCACTTCAAGTCCCGGTGTCCGGGAGGGTGCAGTCGTTGCCAAGTATGAGAACAACGATACCTATTCCCGGACCGCTCACTCTCTGGTC<br>GGTTACTATACCACGGACAATGAAACAGTATCGGAAGCCGATATTCTCACTTTCATGAAAGCAAGGCTTCCAACGTACATGGTGCCAAG<br>CCACCTCTGCTGTCTGGAAGGCGCACTGCCTGTGACGATTAACGGAAAGCTCGACGTCCGGAGATTGCCGGAGATTATCAACGACTCCG<br>CGCAGTCCTCGTACAGCCCACCAAGGAACATAATCGAGGCCAAGATGTGCAGACTGTGGGAATCCGCCTTGGGAATGGAGCGATGCGG<br>TATCGACGACGACCTGTTCAAACTGGGTGGCGACAGCATCACATCTTTGCATCTCGTGGCCCAGATTCACAACCAGGTGGGCTGCAAGA<br>TCACCGTTCGGGATATATTTGAACATCGTACCGCCCGAGCCCTCCATGATCACGTCTTCATGAAGGACTCCGACCGGAGTAATGTGACTC<br>AGTTCCGAACCGAACAAGGGCCGGTCATCGGCGAGGCGCCCCTACTGCCGATTCAAGACTGGTTTTTGTCAAAGGCTCTGCAGCATCCG<br>ATGTATTGGAATCACACTTTCTACGTCCGAACGCCAGAGCTGGATGTTGATTCCTTAAGCGCTGCTGTCAGGGACTTGCAACAGTATCAC<br>GATGTTTTCCGCATGCGACTCAAGCGCGAGGAAGTCGGATTCGTGCAGTCCTTTGCTGAGGACTTCTCTCCTGCCCAGCTTCGGGTGCTG<br>AACGTAAAAGATGTTGACGGGTCCGCGGCCGTCAACGAGATATTGGATGGGTGGCAGTCTGGCTTCAACCTTGAGAACGGACCCATTG<br>GTTCCATTGGCTACCTACATGGGTATGAAGACCGATCCGCGCGAGTCTGGTTCTCCGTTCACCATATGGCCATTGACACCGTCAGCTGGC<br>AGATCCTTGTCCGTGACCTGCAGACGCTGTACCGAAATGGAAGCCTCGGAAGCAAGGGCAGCAGTTTCCGGCAGTGGGCTGAAGCCAT<br>CCAAAATTACAAGGCGTCAGACTCTGAGAGGAACCATTGGAATAAGCTCGTCATGGAAACAGCTTCCAGCATATCCGCATTGCCTACGT<br>CAACCGGTTCGCGCGTGCGCCTGAGCAGAAGTTTGAGCCCTGAGAAGACAGCCTCACTGATCCAAGGAGGAATCGATCGACAGGATGT |

| | |
|---|---|
| | CTCCGTGTACGACTCCCTCCTGACTTCAGTTGGATTGGCGCTCCAACATATCGCTCCAACCGGCCCAAGTATGGTTACGATCGAGGGACA<br>TGGCCGTGAAGAAGTGGATCAGACACTGGATGTGAGCCGCACCATGGGTTGGTTCACCACCATGTATCCATTTGAAATTCCCCGTCTCA<br>GCACCGAGAACATTGTTCAAGGAGTCGTCGCTGTGAGCGAACGGTTCAGACAGGTGCCTGCCCGTGGCGTCGGGTATGGAACCTTGTAC<br>GGCTATACTCAACACCCGCTGCCCCAGGTGACCGTCAACTACCTGGGCCAGCTCGCCCGCAAGCAATCGAAGCCAAAGGAATGGGTCCT<br>CGCGGTGGGCGACAACGAATTTGAATACGGACTCATGACTAGCCCAGAGGACAAAGACCGGAGCTCTTCTGCCGTCGACGTCACGGCC<br>GTGTGTATTGACGGCACTATGATCATCGATGTGGACAGTGCTTGGAGCCTTGAGGAGAGCGAGCAATTCATCTCGAGCATCGAGGAAGG<br>ACTGAACAAGATCCTCGACGGCAGGGCAAGTCAGCAAACCTCGCGATTCCCGGATGTTCCTCAACCGGCGGAGACATATACGCCGTATT<br>TCGAGTATCTGGAACCTCCACGACAGGGACCGACGCTGTTCCTGCTGCCGCCGGGCGAAGGAGGCGCCGAGAGTTACTTCAACAACATC<br>GTCAAGCGCCTGCGTCAGACAAATATGGTGGTCTTCAACAACTACTACTTGCACAGCAAACGCCTGCGCACGTTCGAGGAGCTGGCGGA<br>AATGTATCTCGACCAAGTACGCGGCATCCAACCACACGGACCGTACCACTTCATCGGATGGAGCTTCGGAGGAATTCTCGCAATGGAAA<br>TGTCGCGGCGACTGGTAGCCTCGGACGAGAAGATTGGCTTCCTCGGTATTATCGACACCTATTTCAACGTGCGGGGAGCGACACGCACC<br>ATTGGCTTGGGGGACACTGAGATTCTGGACCCGATCCATCACATCTACAATCCCGATCCGGCCAACTTCCAACGCCTGCCCTCTGCAAC<br>AGATCGCATTGTGCTGTTCAAGGCCATGAGGCCGAACAACAAGTACGAATCCGAGAACCAGCGTCGCCTGTACGAGTACTATGACGGC<br>ACTCGACTCAACGGACTGGACAGCTTGTTACCAAGCGATTCCGACGTCCAGCTGGTCCCGCTTACGGACGATACACACTTTTCCTGGGTC<br>GGAAATCCACAACAGGTGGAGCAGATGTGTGCGACTATCAAGGAACACCTCGCTCGCTATTTGATAGCTTGGCGCGCCTCGAGGGGGG<br>GC |
| eGFP<br>(F4) | TGCGACTATCAAGGAACACCTTGCGACTATCAAGGAACACCTCGCTCGCTATTTGATAGCTTGGCGCGCCCGGTACCGGTAGAAAAAAT<br>GAGTAAAGGAGAAGAACTTTTCACTGGAGTTGTCCCAATTCTTGTTGAATTAGATGGTGATGTTAATGGGCACAAATTTTCTGTCAGTGG<br>AGAGGGTGAAGGTGATGCAACATACGGAAAACTTACCCTTAAATTTATTTGCACTACTGGAAAACTACCTGTTCCATGGCCAACCCTGG<br>TCACCACCCTGACCTACGGCGTGCAGTGCTTCTCCCGTTACCCTGATCATATGAAGCGGCACGACTTCTTCAAGAGCGCCATGCCTGAGG<br>GATACGTGCAGGAGAGGACCATCTTCTTCAAGGACGACGGGAACTACAAGACACGTGCTGAAGTCAAGTTTGAGGGAGACACCCTCGT<br>CAACAGGATCGAGCTTAAGGGAATCGATTTCAAGGAGGACGGAAACATCCTCGGCCACAAGTTGGAATACAACTACAACTCCCACAAC<br>GTATACATCATGGCCGACAAGCAAAAGAACGGCATCAAAGCCAACTTCAAGACCCGCCACAACATCGAAGACGGCGGCGTGCAACTCG<br>CTGATCATTATCAACAAAATACTCCAATTGGCGATGGCCCTGTCCTTTTACCAGACAACCATTACCTGTCCACACAATCTGCCCTTTCGA<br>AAGATCCCAACGAAAAGAGAGACCACATGGTCCTTCTTGAGTTTGTAACAGCTGCTGGGATTACACATGGCATGGATGAACTATACAAA<br>CACCACCACCACCACCACTAAcagctttcttgtacaaagtggtataattca |

Nucleotide sequences of the fragments used to re-assemble *pcbAB*. Lowercase nucleotides are overhangs homologous to the destination vector.

**Table S6.**

| Plasmid Name | Description | Reference |
|---|---|---|
| pE-YA | Entry vector for assembling megasynthases in yeast; yeast – *E. coli* shuttle vector | (*52*) |
| pMDC84 | Plant expression plasmid; *E. coli* – agrobacteria shuttle vector | (*53*) |
| pTWIST | Entry vector for synthesized genes purchased from TWIST Bioscience | Twist Bioscience, South San Francisco, CA |
| pDONR | Donor vector for synthesized genes purchased from Gene Universal | Invitrogen |
| pDEST | Destination vector | Invitrogen |
| pDGB3 | Plant expression vector | (*54*) |
| pUPD2 | Entry vector for cloning in *E. coli* | (*54*) |
| pDEST-URA-Ub10 | *E. coli* / yeast / *A. tumefaciens* shuttle vector for expression of genes in plant hosts. | This study |
| pDONR-NpgAstop | Donor vector containing NpgA +/- a stop codon | This study |
| pEY-pcbAB | Entry vector containing assembled *pcbAB* | This study |
| pUPD2-IPNS | Entry vector containing IPNS | This study |
| pUPD2-IAT | Entry vector containing IAT | This study |
| pUPD2-PenV | Entry vector containing PenV | This study |
| pUPD2-PenM | Entry vector containing PenM | This study |
| pUPD2-PaaT | Entry vector containing PaaT | This study |
| pMDC83-NpgA-GFP | Plant expression plasmid containing NpgA fused to eGFP under the control of the CaMV35S promoter | This study |
| pMDC83-NpgA | Plant expression plasmid containing NpgA | This study |
| pMDC32-NpgA | Plant expression plasmid containing NpgA under the control of the CaMV35S promoter | This study |
| pMDC84-Pc1304050-GFP | Plant expression plasmid containing Pc1304050 fused to eGFP under the control of the CaMV35S promoter | This study |
| pMDC32-Pc13g04050 | Plant expression plasmid containing Pc1304050 under the control of the CaMV35S promoter | This study |
| pDEST-URA-Ub10-native | Plant expression plasmid containing pcbAB fused to eGFP under the control of the UBQ10 promoter | This study |

| | | |
|---|---|---|
| pcbAB-GFP | | |
| pDGB3-alpha1-GFP-IPNS | Plant expression plasmid containing IPNS fused to eGFP under the control of the CaMV35S promoter | This study |
| pMDC84-IPNS-GFP | Plant expression plasmid containing IPNS fused to eGFP under the control of the CaMV35S promoter | This study |
| pMDC84-IPNS | Plant expression plasmid containing IPNS | This study |
| pDGB3-alpha1-IPNS | Plant expression plasmid containing IPNS under the control of the CaMV35S promoter | This study |
| pDGB3-alpha1-GFP-IAT | Plant expression plasmid containing IAT and eGFP under the control of the CaMV35S promoter | This study |
| pMDC84-IAT | Plant expression plasmid containing IAT | This study |
| pDGB3-alpha1-IAT | Plant expression plasmid containing IAT under the control of the CaMV35S promoter | This study |
| pDGB3-alpha2-IAT | Plant expression plasmid containing IAT | This study |
| pDGB3-omega1-IAT/IPNS | Plant expression plasmid containing IPNS and IAT under the control of the CaMV35S promoter | This study |
| pDGB3-alpha1-GFP-PenM | Plant expression plasmid containing PenM and eGFP under the control of the CaMV35S promoter | This study |
| pMDC84-PaaT-GFP | Plant expression plasmid containing PaaT and eGFP under the control of the CaMV35S promoter | This study |

Plasmids used in this study

**Table S7.**

| Experiment/ Collection | Treatment | Rep. | Total Leaf Fresh Weight (g) | Total Leaf Dry Weight (DW; mg) | Sample DW Used For Extraction (mg) | Detection Status | | | | Absolute Quantification (µg/g dw) | | | Normalized Peak Area |
|---|---|---|---|---|---|---|---|---|---|---|---|---|---|
| | | | | | | ACV | IPN | PenG | PenV | ACV | Pen G | POA | (IPN) |
| 2/21 Infiltration 2/25 collection | Mock | 1 | 1.362 | 92 | 25.2 | No | No | No | No | 0.024 | 0.023 | 0.749 | 0.0009 |
| | | 2 | 1.533 | 122 | 21.7 | No | No | No | No | 0.034 | 0.018 | 0.579 | 0.0016 |
| | | 3 | 1.392 | 88 | 23.6 | No | No | No | No | 0.003 | 0.012 | 0.623 | 0.0016 |
| | | 4 | 1.579 | 96 | 24.3 | No | No | No | No | 0.001 | 0.009 | 0.633 | 0.0013 |
| | P19 | 5 | 1.469 | 119 | 24.1 | No | No | No | No | 0.047 | 0.014 | 0.582 | 0.0009 |
| | | 6 | 1.456 | 113 | 22.2 | No | No | No | No | 0.043 | 0.015 | 0.478 | 0.0011 |
| | | 7 | 1.758 | 127 | 24.4 | No | No | No | No | 0.038 | 0.012 | 0.436 | 0.0012 |
| | | 8 | 1.586 | 122 | 24.9 | No | No | No | No | 0.011 | 0.015 | 0.546 | 0.0016 |
| | **p19+ACVS+ pc13g04050 +IPNS/IAT (All genes)** | 9 | 2.717 | 226 | 23.1 | **Yes** | **Yes** | **Yes** | No | 0.404 | 4.005 | 0.922 | 0.354 |
| | | 10 | 1.692 | 145 | 24.9 | **Yes** | **Yes** | **Yes** | No | 0.644 | 4.599 | 1.563 | 0.478 |
| | | 11 | 2.178 | 160 | 22.2 | **Yes** | **Yes** | **Yes** | No | 0.557 | 12.124 | 1.046 | 0.464 |
| | | 12 | 2.08 | 164 | 24.8 | **Yes** | **Yes** | **Yes** | No | 0.515 | 3.655 | 19.321 | 0.429 |
| | | 13 | 2.897 | 242 | 24.9 | **Yes** | **Yes** | **Yes** | No | 0.386 | 2.815 | 0.694 | 0.288 |
| | p19+ACVS+pc13g04050+ IPNS/IAT+**POA** **(All genes+POA)** | 14 | 2.48 | 185 | 25.3 | **Yes** | **Yes** | **Yes** | No | 0.760 | 9.327 | 525.641 | 1.016 |
| | | 15 | 1.592 | 133 | 24.2 | **Yes** | **Yes** | **Yes** | No | 0.517 | 2.216 | 392.044 | 0.135 |
| | | 16 | 2.441 | 183 | 23.7 | **Yes** | **Yes** | **Yes** | No | 0.953 | 6.173 | 442.119 | 0.750 |
| | | 17 | 1.942 | 147 | 23.2 | **Yes** | **Yes** | **Yes** | No | 0.647 | 8.964 | 495.526 | 0.746 |
| | | 18 | 2.261 | 199 | 24.6 | **Yes** | **Yes** | **Yes** | No | 0.937 | 8.500 | 426.949 | 1.451 |

| Experiment/ Collection | Treatment | Rep. | Sample Label for LC-MS analysis | Leaf Fresh Weight (FW; gram) | Leaf Dry Weight (DW; gram) | Sample DW Used For Extraction (mg) | Detection Status | | | Absolute Quantification (µg/g dw) | | Normalized Peak Area (IPN) |
|---|---|---|---|---|---|---|---|---|---|---|---|---|
| | | | | | | | ACV | IPN | PenG | ACV | PenG | |
| 2/27 infiltration 3/3 collection | Mock | 1 | 1 | 2.102 | 0.148 | 19.5 | No | No | No | 0.001 | 0.007 | 0.0012 |
| | | 2 | 2 | 1.978 | 0.131 | 19.4 | No | No | No | 0.011 | 0.024 | 0.0012 |
| | | 3 | 3 | 2.066 | 0.126 | 20.4 | No | No | No | 0.000 | 0.012 | 0.0001 |
| | P19 | 1 | 4 | 1.516 | 0.135 | 19.4 | No | No | No | 0.002 | 0.027 | 0.0022 |
| | | 2 | 5 | 1.671 | 0.112 | 19.7 | No | No | No | 0.006 | 0.012 | 0.0007 |
| | | 3 | 6 | 1.546 | 0.113 | 19.7 | No | No | No | 0.0042 | 0.031 | 0.0014 |
| | p19+ACVS+pc13g04050 +IPNS/IAT **(All genes)** | 1 | 7 | 1.967 | 0.173 | 19.7 | Yes | Yes | Yes | 0.720 | 5.299 | 0.648 |
| | | 2 | 8 | 2.205 | 0.187 | 19.7 | Yes | Yes | Yes | 0.793 | 4.867 | 0.750 |
| | | 3 | 9 | 2.005 | 0.183 | 19.2 | Yes | Yes | Yes | 0.916 | 10.452 | 0.849 |
| | All genes+penM | 1 | 10 | 2.185 | 0.176 | 19.3 | Yes | Yes | Yes | 0.723 | 12.189 | 0.501 |
| | | 2 | 11 | 1.873 | 0.153 | 19.5 | Yes | Yes | Yes | 0.753 | 5.329 | 0.591 |
| | | 3 | 12 | 2.089 | 0.167 | 20.8 | Yes | Yes | Yes | 0.727 | 6.106 | 0.463 |
| | All-genes+43PaaT | 1 | 13 | 1.692 | 0.139 | 19.9 | Yes | Yes | Yes | 0.928 | 21.239 | 0.802 |
| | | 2 | 14 | 1.37 | 0.105 | 19.6 | Yes | Yes | Yes | 0.467 | 24.046 | 0.460 |
| | | 3 | 15 | 1.771 | 0.144 | 19.9 | Yes | Yes | Yes | 0.801 | 32.175 | 1.239 |
| | All genes+84PaaT | 1 | 16 | 1.049 | 0.097 | 20.3 | Yes | Yes | Yes | 0.440 | 18.033 | 0.401 |
| | | 2 | 17 | 1.227 | 0.108 | 19.6 | Yes | Yes | Yes | 0.679 | 20.732 | 0.582 |
| | | 3 | 18 | 1 | 0.091 | 20.4 | Yes | Yes | Yes | 0.607 | 33.891 | 0.758 |
| | All-genes+ penM+43PaaT | 1 | 19 | 1.599 | 0.144 | 20 | Yes | Yes | Yes | 0.559 | 13.200 | 0.436 |
| | | 2 | 20 | 1.728 | 0.15 | 19.8 | Yes | Yes | Yes | 0.908 | 19.997 | 0.753 |
| | | 3 | 21 | 1.696 | 0.151 | 20.3 | Yes | Yes | Yes | 1.323 | 39.046 | 1.192 |
| | All-genes+ penM+84PaaT | 1 | 22 | 1.349 | 0.163 | 19.3 | Yes | Yes | Yes | 0.590 | 10.967 | 0.652 |
| | | 2 | 23 | 1.729 | 0.15 | 19.2 | Yes | Yes | Yes | 0.649 | 14.024 | 0.600 |
| | | 3 | 24 | 1.708 | 0.138 | 19.4 | Yes | Yes | Yes | 0.387 | 10.590 | 0.150 |

Batch infiltration raw data for quantification of ACV, IPN, and PenG from dried *N. benthamiana* leaf extracts.

**Table S8.**

| Treament Group | Sample ID | Mean ZOI Diameter (mm) | Calculated Conc. in 25 µL (mM) | Calculated Mass in 25 µL extract (µg) | Total Mass in 1 mL Extract (µg) | Sample Dry Weight (g) | Final Conc. In leaf tissue (µg/g dw) |
|---|---|---|---|---|---|---|---|
| P19+ACVS+pc13g04050+IPNS/IAT+phl+43-PaaT | P1 | 17.5 | 0.0136 | 0.1138 | 4.55 | 0.264 | 17.23 |
| | P2 | 21 | 0.0312 | 0.2612 | 10.43 | 0.366 | 28.51 |
| | P3 | 23 | 0.0502 | 0.4199 | 16.79 | 0.338 | 49.67 |
| P19+ACVS+pc13g04050+IPNS/IAT+phl+84-PaaT | P1 | 23.5 | 0.0566 | 0.4729 | 18.92 | 0.185 | 102.25 |
| | P2 | 20 | 0.0246 | 0.206 | 8.24 | 0.316 | 26.08 |
| | P3 | 22.5 | 0.0446 | 0.3729 | 14.92 | 0.342 | 43.62 |

**Determination of penicillin G concentration in different plant extract treatments based on bioassay zone of inhibition.** The penicillin G concentrations of the six samples across two treatments were determined based on the linear regression model derived from the *Staphylococcus saprophyticus* bioassay. By applying the measured zone of inhibition diameters to the regression equation ($Y = 9.700X + 35.60$), the millimolar concentrations in the 25 µL were calculated. The final concentration (µg/g) in each plant extract was determined after normalizing the amounts to the respective dry weights of leaf tissue. The ZOI diameter for each sample was measured in two technical replicates. Abbreviations: ZOI: Zone of inhibition; dw: dry weight.
